# Overcoming the “feast or famine” effect: improved interaction testing in genome-wide association studies

**DOI:** 10.1101/2024.02.13.580168

**Authors:** Huanlin Zhou, Mary Sara McPeek

## Abstract

In genetic association analysis of complex traits, detection of interaction (either GxG or GxE) can help to elucidate the genetic architecture and biological mechanisms underlying the trait. Detection of interaction in a genome-wide interaction study (GWIS) can be methodologically challenging for various reasons, including a high burden of multiple comparisons when testing for epistasis between all possible pairs of a set of genomewide variants, as well as heteroscedasticity effects occurring in the presence of GxG or GxE interaction. In this paper, we address the problem of an even more striking phenomenon that we call the “feast or famine” effect that occurs when testing interaction in a genomewide context. We show that in any given GxE GWIS, the type 1 error of standard interaction tests performed genomewide can vary widely from the nominal level, where the actual type 1 error in any given GWIS varies as a predictable function of the observed trait and environmental values. Using standard methods, some GWISs will have systematically underinflated p-values (“feast”), and others will have systematically overinflated p-values (“famine”), which can lead to false detection of interaction, reduced power, inconsistent results across studies, and failure to replicate true signal. This startling phenomenon is specific to detection of interaction in a GWIS, and it may partly explain why such detection has often proved challenging and difficult to replicate. We show that the feast or famine effect occurs across a wide range of GxE analysis methods, including but not limited to (1) testing interaction in a linear or linear mixed model (LMM) using standard approaches such as t-tests/Wald tests, likelihood ratio tests, or score tests; (2) doing a combined interaction-association test in a linear model or LMM using standard approaches; (3) testing interaction with multiple environments or multiple SNPs, where these are modeled as random effects in a LMM using standard approaches; (4) performing tests of interaction in a GWIS where significance is assessed using permutation of the trait residuals. We show theoretically that the key cause of this phenomenon is which variables are conditioned on in the analysis. Using this insight, we have developed (i) a diagnostic ratio to detect which GWASs are subject to a strong “feast or famine” effect and (ii) the TINGA method to adjust the interaction test statistics to make their p-values approximately uniform under the null hypothesis. In simulations we show that TINGA both controls type 1 error and improves power. TINGA allows for covariates and population structure through use of a linear mixed model and accounts for heteroscedasticity. We apply TINGA to detection of epistasis in a study of flowering time in *Arabidopsis thaliana*.

**Author summary:** Testing for interactions in GWAS can lead to insight into biological mechanisms, but poses greater challenges than ordinary genetic association GWAS. When testing for interaction in a GWAS setting with one fixed SNP or environmental variable, the standard test statistics may not have the expected statistical properties under the null hypothesis, which can lead to false detection of interaction, inconsistent results across studies, reduced power, and failure to replicate true signal. We propose the TINGA method to adjust the test statistics so that the null distribution of their p-values is closer to uniform. Through simulations and real data analysis, we illustrate the problems with the standard analysis and the improvement of our proposed method.

## Introduction

It is well-known that the effects of a genetic variant on a trait can be different for individuals with different environments, such as age [1], sex [2–5], lifestyle [6] and other exposures [7]. The genetic effects can also depend on other variants, either from the same genome [8, 9] or the genome of another species (such as pathogen and host [10], mother and offspring [11]). Detection of such interaction effects can enhance the ability to identify genetic effects that would otherwise be reduced or masked [12]; they are considered as one of the reasons why results of marginal association studies are sometimes hard to replicate [13]; they are believed to account for a large part of missing heritability [14–16] ; and they can elucidate the genetic architecture of complex traits and diseases [12, 17, 18] and benefit many areas such as public health [19] and agriculture [20, 21]. Much previous work has been done to develop appropriate methods for detecting interactions in GWAS, aiming to improve computational efficiency, reduce false positives and increase power [4, 22–29].

One challenge specific to epistasis detection is that, because of the large number of tests, exhaustive search for epistatic effects in a GWAS context has a larger computational burden and lower statistical power than ordinary trait-variant association studies. To deal with this issue, various methods have been developed that correct for multiple testing while still remaining powerful [30, 31]. Another option is to reduce the number of tests by a two-stage approach: first select a subset of SNPs that are more likely to be involved in interaction and then test for interaction among them [22, 26, 32, 33].

Previous work [34–36] has found that it can be hard to replicate interactions in GWAS. This can occur for a variety of reasons. For example, in some cases, an apparent epistatic effect that is detected could be due to an unsequenced causal variant [34, 37, 38]. Another important issue that has been identified is heteroscedasticity [39–41] that can result under the null model when, for example, interaction is present between one of the two tested variables and some other variable not included in the model or when the null model is misspecified in some other way. If not accounted for, this heteroscedasticity can lead to excess type 1 error [39–41].

Many scenarios of testing for GxG or GxE in a GWAS context involve fixing one genetic variant or environmental factor and performing an interaction GWAS by testing the fixed variable for interaction with each genetic variant across the genome. Systematically inflated or deflated p-values in such an interaction GWAS have been previously reported, based on both data and simulations [38–40]. Even under simplified assumptions, in the absence of problems such as heteroscedasticity, it has been noted that type 1 error rates and genomic control inflation factors are highly variable across such interaction GWASs [39, 40]. In this paper, we develop a deeper and more detailed understanding of this phenomenon, which we call the “feast or famine” effect in interaction GWAS. We frame this problem as resulting from the choice of variables to condition on and show how changing this choice has the potential to resolve the problem. Our framework also explains clearly why the “feast or famine” effect only occurs in interaction GWAS, not in ordinary association GWAS. We implement our ideas in a method we call TINGA (Testing INteraction in GWAS with test statistic Adjustment), in which we adjust the t-statistic for interaction by re-centering and re-scaling it using the null conditional mean and conditional variance of its numerator, with a more appropriate choice of conditioning variables. In simulations, we demonstrate the ability of TINGA to greatly reduce or eliminate the “feast or famine” effect while controlling type 1 error and increasing power. We also develop a useful diagnostic that accurately predicts the magnitude and direction of the “feast or famine” effect in any given data set. We apply the methods to detect epistasis in a GWAS for flowering time in *Arabidopsis thaliana*.

## Materials and methods

We consider the problem of testing for interaction, either *G × E* or *G × G*, in a GWAS context. In a sample of *n* individuals, let *Y* be an *n ×* 1 trait vector, and let *G* be an *n × m* matrix of genotypes for a set of genome-wide variants. Let *Z* be an *n ×* 1 vector that, in the case of *G × E* testing, represents the environmental variable that we wish to test interaction with and in the case of *G × G* testing, represents the genotype at a particular variant that we wish to test interaction with (where we assume that *Z* is removed from the matrix *G* in that case). In addition, we can allow for an *n × k* matrix *U* of covariates (including intercept), where these are implicitly taken as fixed and are conditioned on throughout the analysis. By “testing interaction in a GWAS context,” we mean that for each *j* in {1, …, *m*}, we test for interaction between *G*_*j*_ and *Z* in a linear or linear mixed model (LMM) for *Y*, where *G*_*j*_ is the *j*th column of *G*.

In this section, we first describe what we call the “feast or famine” effect for testing interaction in a GWAS context. We explain how the “feast or famine” effect can result in some GWASs having systematically overinflated interaction p-values, reducing power, while others have systematically underinflated p-values, resulting in excess type 1 error. In what follows, we focus our exposition on the t-statistic for testing interaction, but the “feast or famine” effect is very general. We show that the feast or famine effect occurs across a wide range of GxE analysis methods, including but not limited to (1) testing interaction in a linear or linear mixed model (LMM) using standard approaches such as t-tests/Wald tests, likelihood ratio tests, or score tests; (2) doing a combined interaction-association test in a linear model or LMM using standard approaches; (3) testing interaction with multiple environments or multiple SNPs, where these are modeled as random effects in a LMM using standard approaches [22, 28]; (4) performing tests of interaction in a GWIS where significance is assessed using permutation of the trait residuals. We show that the “feast or famine” effect does not occur in ordinary GWAS for testing association between a trait and each genetic variant, but only when testing interaction in a GWAS context. Next we describe our TINGA method to correct the interaction test statistics to greatly reduce or eliminate this effect.

In the simplest setting in which there are no covariates and no population sub-structure, we let *T*_*j*_ denote the *t*-statistic for testing interaction between *G*_*j*_ and *Z*, i.e., for testing *H*_0_ : *δ* = 0, in the following linear model:

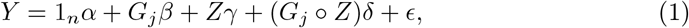

where 1_*n*_ is a vector of length *n* with every entry equal to 1, *α, β, γ* and *δ* are unknown scalar parameters, 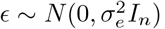, where 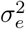 is unknown and *I*_*n*_ is the *n × n* identity matrix, and where, for any two vectors *a* and *b*, both of length *n*, we define *a* ° *b* to be the vector of length *n* with *i*th element 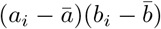, where, e.g., 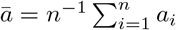. _(Note that the test statistics *Tj*_ would remain exactly the same if we replaced *G*_*j*_ ° *Z* in (1) by the element-wise product of the vectors *G*_*j*_ and *Z*, but choosing to center the variables before multiplying them has various advantages such as reducing potential collinearity and making the coefficients more interpretable.)

### The “feast or famine” effect: what we thought we knew about testing interaction in a GWAS context was wrong

For simplicity, we first focus the exposition on *G × E* interaction testing. An essential feature of testing *G × E* interaction in a GWAS context is that we obtain a set of *m* test statistics *T*_*j*_, *j* ∈ {1, …, *m*}, where *T*_*j*_ ≡ *T*_*j*_ (*G*_*j*_, *Z, Y*), with the same *Y* and *Z* used in all the test statistics and only *G*_*j*_ varying. As a thought experiment, imagine the simplest possible null scenario in which *Y, Z* and the columns of *G* are mutually independent, with the elements of *Y* drawn as i.i.d. *N*(*µ, σ*^2^), the elements of *Z* drawn as i.i.d. from some distribution *F*_*Z*_, and the elements of *G*_*j*_ drawn as i.i.d. from some distribution *F*_*Gj*_, for *j* = 1, …, *m*. What would be the distribution of (*T*_1_, …, *T*_*m*_) in this case? It is well-known that for any given *j*, the distribution of *T*_*j*_ in this case is the (central) Student’s *t* distribution on *n* − 4 df, which we denote by 𝒯_*n*−4_. Thus, it is tempting to assume that *T*_1_, …, *T*_*m*_ must be approximately i.i.d. draws from 𝒯_*n*−4_, but that is (perhaps surprisingly) incorrect.

In this simple scenario, we show that it is most appropriate to think of *T*_1_, …, *T*_*m*_ as i.i.d. draws from some distribution whose mean is 0 and whose variance is a function of (*Y, Z*). For some choices of (*Y, Z*), the variance of the resulting *T*_*j*_ ‘s is larger than 1 (where 1 is the approximate variance of 𝒯_*n*−4_ for large *n*), while for other choices of (*Y, Z*), the variance of the resulting *T*_*j*_ ‘s is smaller than 1. Thus, if we used 𝒯_*n*−4_ to calculate p-values *p*_1_, …, *p*_*m*_ for *T*_1_, …, *T*_*m*_, respectively, which would be the standard approach, then in one GWAS these p-values might be systematically too big on average, in a second GWAS these p-values might be systematically too small on average, and in a third GWAS, they might be about right (where by “about right” we mean approximately i.i.d. uniform under the null).

This can easily be observed in simulations (see also [39, 40]). Fig 1 shows four histograms, each of which depicts the p-values *p*_1_, …, *p*_*m*_ for a *G × E* GWAS obtained as described above, where *n* is 1,000, *m* is 5,000, *F*_*Z*_ is taken to be Bernoulli(.2), and *F*_*Gj*_ is taken to be Bernoulli(*f*_*j*_) for *j* = 1, …, *m*, where *f*_1_, …, *f*_*m*_ are drawn as i.i.d. Unif(.1, .9), to mimic unlinked genotypes from a haploid organism or an inbred line. In Panel A of Fig 1, the p-values are seen to be systematically overinflated, while in Panel B of Fig 1, the p-values are seen to be systematically underinflated. The information in Table 1 supports this conclusion, where we can see that for Panel A, the s.d. of the interaction t-statistics is *<* 1 and the genomic control inflation factor is *<* 1, while for Panel B the opposite holds. We repeated this experiment 400 times, and in each replicate, we tested whether the 5,000 p-values were i.i.d. Uniform(0,1) distributed under the null hypothesis (which is equivalent to testing whether the 5,000 interaction t-statistics are i.i.d. 𝒯_*n*−4_ distributed) using the two-sided equal local levels (ELL) test as implemented in qqconf [42]. (See S1 Text for an R script to perform this test.) In 190 out of 400, i.e., 47.5%, of the replicates, the two-sided ELL test for uniformity was rejected at level .05, clearly showing that the t-statistics for interaction in a GWAS are not i.i.d. 𝒯_*n*−4_distributed under the null hypothesis.

This effect seems to be very general and also occurs when, e.g., *F*_*Z*_ and *F*_*Gj*_ are taken to be Gaussian or Binomial, as we show later. Furthermore, if instead of a t-test for interaction, we apply a likelihood ratio chi-squared test or F-test for interaction to the same simulated data sets, we get essentially indistinguishable histograms to those in Fig 1 (which is perhaps not surprising since they are asymptotically equivalent tests), and the same 190 replicates out of 400 are rejected by the ELL test for uniformity of the p-values, showing that the likelihood ratio chi-squared test and F-test for interaction are also subject to the “feast or famine” effect.

**Fig 1.**
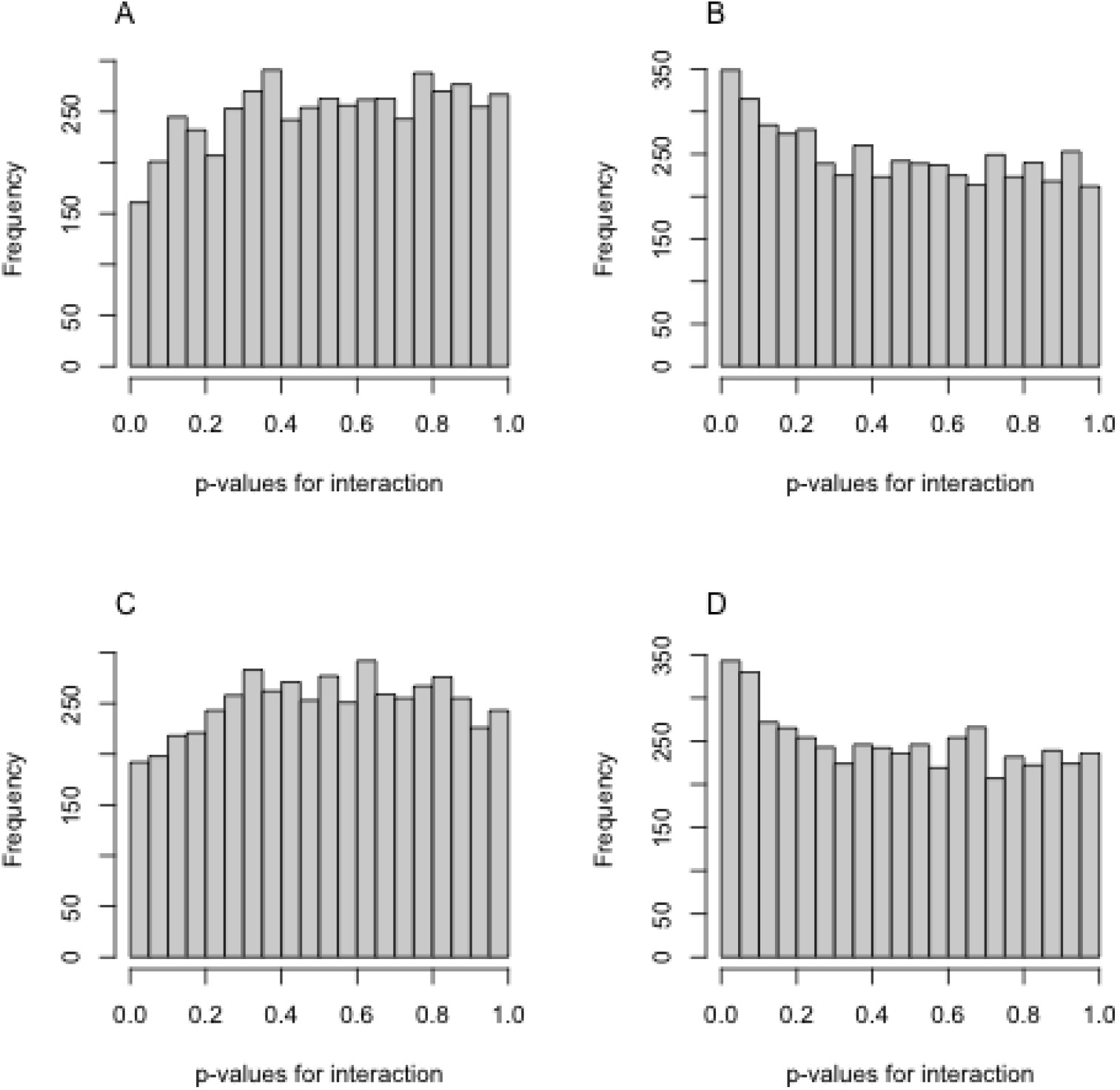
Histograms of p-values for t-tests for interaction in a GWAS when the null hypothesis is true. Each histogram is based on a replicate of (*Y, Z*) and 5,000 genotypes, *G*_1_, …, *G*_5000_. In each histogram, interaction is tested between *Z* and *G*_*j*_ in the linear model in (1) for *j* = 1, … 5,000, as described in the text, and the 5,000 p-values are computed using the the 𝒯_*n*−4_ distribution and are displayed in the histogram. Panels A and B represent two different replicates of a null simulation as described in the text. In Panel C, the same (*Y, Z*) replicate is used as in Panel A, and a new set of 5,000 genotypes is simulated and used in the interaction tests. Similarly, in Panel D, the same (*Y, Z*) replicate is used as in Panel B, and a new set of 5,000 genotypes is simulated and used in the interaction tests.

Many standard methods are affected by the “feast or famine” effect. In Figs. S4-S8, we show that the feast or famine effect occurs across a wide range of GxE analysis methods, including but not limited to (1) testing interaction in a linear or linear mixed model (LMM) using standard approaches such as t-tests/Wald tests, likelihood ratio tests, or score tests; (2) doing a combined interaction-association test in a linear model or LMM using standard approaches; (3) testing interaction with multiple environments or multiple SNPs, where these are modeled as random effects in a LMM using standard approaches; (4) performing tests of interaction in a GWIS where significance is assessed using permutation of the trait residuals.

**Table 1.**
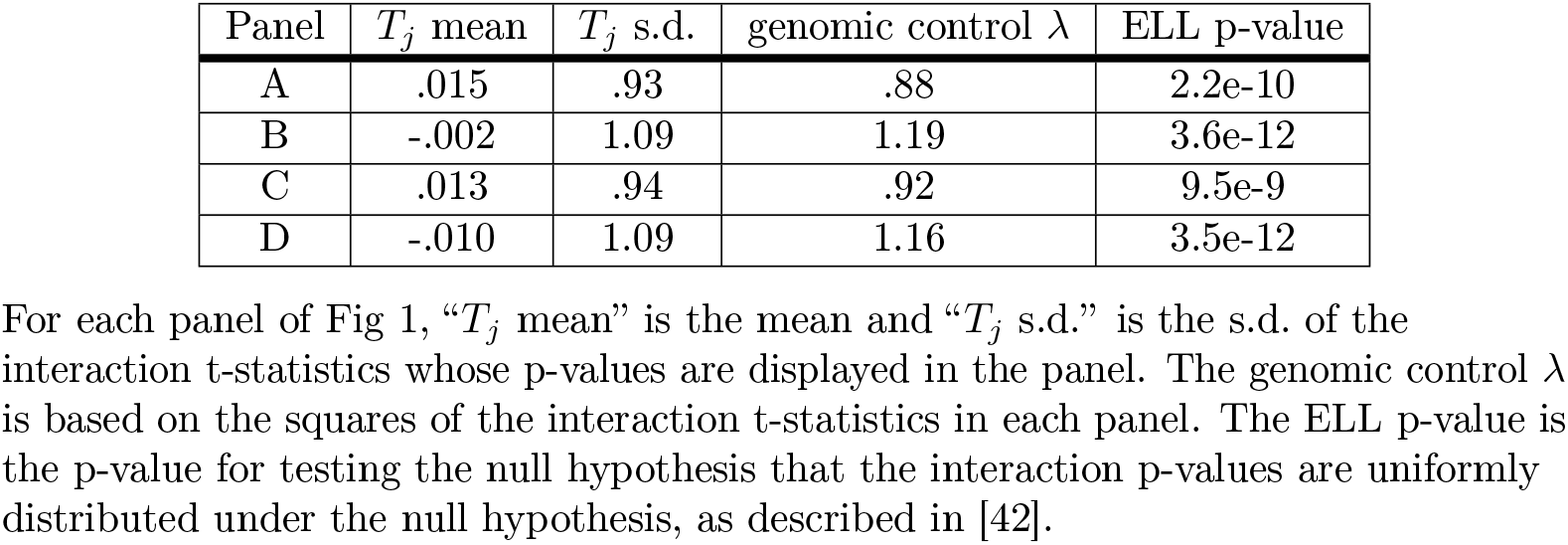
Summary statistics for the examples in Fig 1.

### A deeper understanding

We want to emphasize that we are not simply saying that the interaction p-values *p*_1_, …, *p*_*m*_ from a given GWAS are positively correlated. A further key point is that for a particular *G × E* GWAS, i.e., for a particular choice of (*Y, Z*), it is, in principle, predictable based on (*Y, Z*) whether the p-values *p*_1_, …, *p*_*m*_ will be systematically too large, systematically too small or about right. For example, in Fig 1, when we keep (*Y, Z*) the same as in Panel A and simulate a completely new and independent set of genotypes *G* for testing interaction, as in Panel C, we again see overinflation of the p-values. Similarly, when we keep (*Y, Z*) the same as in Panel B and simulate a completely new and independent set of genotypes *G* for testing interaction, as in Panel D, we again see underinflation of the p-values. This is further supported by the information in Table 1. Thus, use of standard methods would be expected to result in loss of power (“famine”) in some GWASs (e.g., the (*Y, Z*) used in Panels A and C) and excessive type 1 (“feast”) error in other GWASs (e.g., the (*Y, Z*) used in Panels B and D).

To understand why this happens, it is helpful to think about which variables we are conditioning on. The ordinary *t*-statistic for interaction was developed in a non-GWAS context in which it made sense to condition on *G*_*j*_ and *Z* and treat *Y* as random, and in that case, the null conditional distribution of *T*_*j*_ can be proven to be *T*_*n*−4_ in the simple setting described above. As a direct consequence of this, it is also true that the unconditional distribution of *T*_*j*_ is 𝒯_*n*−4_. In other words, if we randomly choose a *G × E* GWAS (i.e., randomly choose (*Y, Z*)), and then randomly choose a null SNP *j* from that GWAS, then *T*_*j*_ has distribution 𝒯_*n*−4_. However, in any particular *G × E* GWAS, *Z* and *Y* are fixed, and only *G*_*j*_ is varying, so it is more appropriate to consider the null conditional distribution of the *t*-statistic for interaction where we condition on *Z* and *Y* and treat *G*_*j*_ as random [39]. We show that even in the simple case described above, conditional on (*Y, Z*), the distribution of *T*_*j*_ depends on (*Y, Z*) and is not 𝒯_*n*−4_. In fact, in the slightly more general null hypothesis scenario when *G*_*j*_ has some marginal effect on *Y* but no interaction with *Z*, we show that not only the null conditional variance of *T*_*j*_ but even its null conditional mean depends on (*Z, Y*).

These same ideas apply to testing *G × G* interaction in a GWAS context if we think of setting *Z* to be the genotype of one particular variant, we exclude *Z* from the columns of *G*, and we consider a GWAS in which we test for interaction between *Z* and *G*_*j*_ for *j* = 1, …, *m* in model (1) using a t-test for interaction. The upshot is that for some *G × E* or *G × G* GWASs, i.e., for some realizations of (*Y, Z*), use of a 𝒯_*n*−4_ distribution to assess significance of interaction will systematically overstate the evidence for interaction (“feast”), while for other *G × E* or *G × G* GWASs, it will systematically understate the evidence for interaction (“famine”). Whether there is feast or famine will depend on the luck of what value of (*Y, Z*) is observed. This statistical phenomenon could be an important explanation of the difficulty in detecting and replicating epistasis and gene-environment interaction that has long been observed.

With this conditioning explanation in mind, one way of thinking of the “feast or famine” effect is that if we average across many interaction GWASs, then the t-statistic for interaction has correct type 1 error, but its false positives are excessively concentrated in some GWASs, and its false negatives are excessively concentrated in some other GWASs. The good news is that, as we show below, (i) we can accurately predict, based on the observed (*Y, Z*), whether the GWAS will be “feast” or “famine” or neither, and (ii) our conditioning explanation implies that by doing conditional calculations, such as we describe below, we should in principle be able to alleviate or entirely eliminate this effect.

### Why doesn’t ordinary (non-interaction) GWAS have the “feast or famine” phenomenon?

We have argued that when testing interaction in a GWAS context, we are actually conditioning on *Y* and *Z* and letting *G*_*j*_ be random, and that the t-statistic for interaction does not have a t-distribution under the null hypothesis when we condition on (*Y, Z*). By a similar argument, we could point out that in an ordinary (non-interaction) GWAS, we are conditioning on *Y* and letting *G*_*j*_ be random, rather than the reverse. Does this also cause a problem for the t-statistic for association? The answer is no. The problem we describe does not occur for ordinary (non-interaction) GWAS, but is specific to interaction GWAS, as we now explain.

First, consider the t-statistic for association in an ordinary GWAS. We consider a slightly more general scenario than before in which there may be additional covariates *U* in the model (where *U* includes an intercept). Suppose the model we use for testing association is

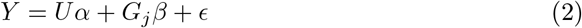

where *Y* is *n ×* 1, *U* is *n × k*, and *G*_*j*_ is *n ×* 1, all as defined before, *α* is an (unknown) *k ×* 1 vector, *β* is the unknown scalar parameter of interest, and 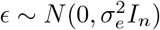, where 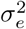 eis unknown.

Define *P*_*U*_ = *I* − *U*(*U*^*T*^*U*)^−1^*U*^*T*^, an *n × n* symmetric matrix. We note that the t-statistic for testing *H*_0_ : *β* = 0 in the model in (2) can be written as

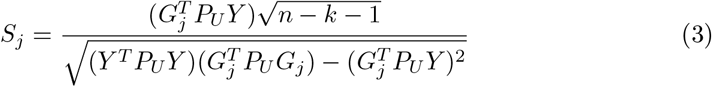

From this formula, it is clear that the t-statistic is symmetric in *G*_*j*_ and *Y*. The symmetry between *G*_*j*_ and *Y* in the ordinary (non-interaction) t-statistic for association means that in large samples, the distribution of the t-statistic under the null hypothesis of no association would be approximately the same regardless of whether we conditioned on *G*_*j*_ and let *Y* be random or conditioned on *Y* and let *G*_*j*_ be random. The only difference would be that *G*_*j*_ would typically be a Binomial or Bernoulli random variable (genotype) and *Y* might commonly be a conditionally normal random variable (phenotype). In very small sample sizes, the difference between the underlying distributions of *G*_*j*_ and *Y* would change the conditional distribution of the t-statistic for association depending on which one you conditioned on, but in typical GWAS sample sizes, the central limit theorem will take effect, and the conditional distribution of the t-statistic for association will be approximately the same in both cases.

This difference between ordinary (non-interaction) GWAS and interaction GWAS can be seen in simulations. We performed *r* = 5,000 replicates of a null simulation similar to that in the previous subsection, except that instead of *F*_*Z*_ being Bernoulli(.2), we made *F*_*Z*_ Bernoulli(*f*_*Zk*_) in replicate *k*, where *f*_*Z*1_, …, *f*_*Zk*_ are i.i.d. Unif(.1, .9). In replicate *k*, we tested interaction between *Z* and *G*_*j*_ (*H*_0_ : *δ* = 0 in Model (1)) for *j* = 1, … *m*, obtaining interaction t-statistics 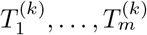. We also tested association between *G*_*j*_ and *Y* in a model with no other covariates except intercept, obtaining ordinary association t-statistics 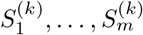 as in (3). We obtain the interaction p-values for 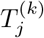 using the 𝒯_*n*−4_ distribution and the ordinary association p-values for 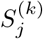 using the 𝒯_*n*−2_ distribution. In this simulation, when we apply the two-sided ELL _test for uniformity at level .05 to the interaction p-values from each replicate, we reject_ 29.3% of the 5,000 replicates as being significantly non-uniform. In contrast, when we apply the same ELL test to the ordinary association p-values from each replicate, we reject just 4.8% of the 5,000 replicates, which is not significantly different from the nominal 5% rate. This verifies that the ordinary GWAS p-values are showing the expected behavior, while the “feast or famine” effect is only showing up in the interaction p-values. This can be seen also in Fig 2 Panel A which depicts a histogram of the genomic control inflation factors for each replicate for the interaction GWASs in red and for the ordinary (non-interaction) GWASs in blue. The narrower blue histogram reflects the expected sampling variability of the GCIF based on 5,000 i.i.d. test statistics. In contrast, the wider red histogram reflects the additional spread due to the “feast or famine effect”, i.e., the fact that conditional on (*Y, Z*) the p-values may be systematically over- or under-inflated compared to uniform. Fig 2 Panel B is similar but for a simulation in which *F*_*Z*_ is Binomial(2, *f*_*Zk*_) in replicate *k* instead of Bernoulli(*f*_*Zk*_) and *F*_*Gj*_ is Binomial(2, *f*_*Gj*_) instead of Bernoulli(*f*_*Gj*_). In S1 Text, a similar pair of histograms can be seen for the case when both *Z* and *G* are normally distributed. Consider the case when *Y* follows a LMM, i.e., the model is as in (1)except that

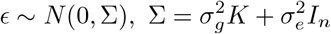

where *K* is a GRM. In this framework, it is also true that the Wald test statistic for association (i.e., the Wald test for *H*_0_ : *β* = 0) is symmetric between *G*_*j*_ and *Y* when Σ is known. Thus, in this case also, ordinary GWAS association testing is essentially not affected by whether we condition on *G*_*j*_ and let *Y* be random or condition on *Y* and let *G*_*j*_ be random.

### TINGA method for correcting t-statistics for interaction in a GWAS

To address the “feast or famine” effect in interaction GWAS, we propose to correct the interaction t-statistics for a given GWAS by subtracting off the null conditional means of their numerators and dividing by the conditional s.d.s given the (*Y, Z*) observed for that GWAS. We call this approach TINGA for “Testing INteraction in GWAS with test statistic Adjustment.”

In the most general case, we consider testing for interaction in the model

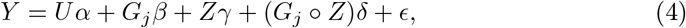

where *Y, U, G*_*j*_, *β, Z, γ*, (*G*_*j*_ *° Z*) and *δ* are as defined before, *α* is a *k ×* 1 vector of unknown coefficients, and 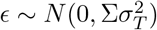, where either Σ = *I*_*n*_ in the case of a linear model, or else Σ = *h*^2^*K* + (1 − *h*^2^)*I*_*n*_ where *K* is as defined before and *h*^2^is an unknown heritability parameter, in the case of a LMM, and where 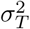 is an unknown parameter. Then the t-statistic for interaction can be written as

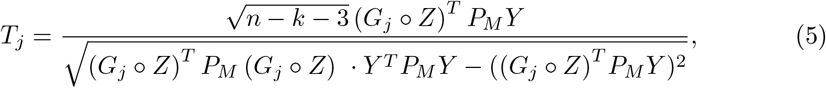

where the “M” in *P*_*M*_ stands for “marginal”, and *P*_*M*_ is a symmetric matrix that removes the marginal effects of *G*_*j*_, *Z*, and *U*, where in the simplest case *U* represents just the intercept, but it may contain additional covariates as needed. We let *M* be the *n ×* (*k* + 2) matrix *M* whose columns are *G*_*j*_, *Z*, and the columns of *U*. Then in the case of a linear model, we have *P*_*M*_ = *I*_*n*_ − *M*(*M*^*T*^*M*)^−1^*M*^*T*^, and in the case of a LMM, we have 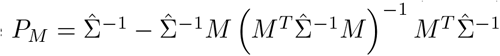, where 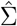 is Σ with the estimated value of *h*^2^ plugged in.

**Fig 2.**
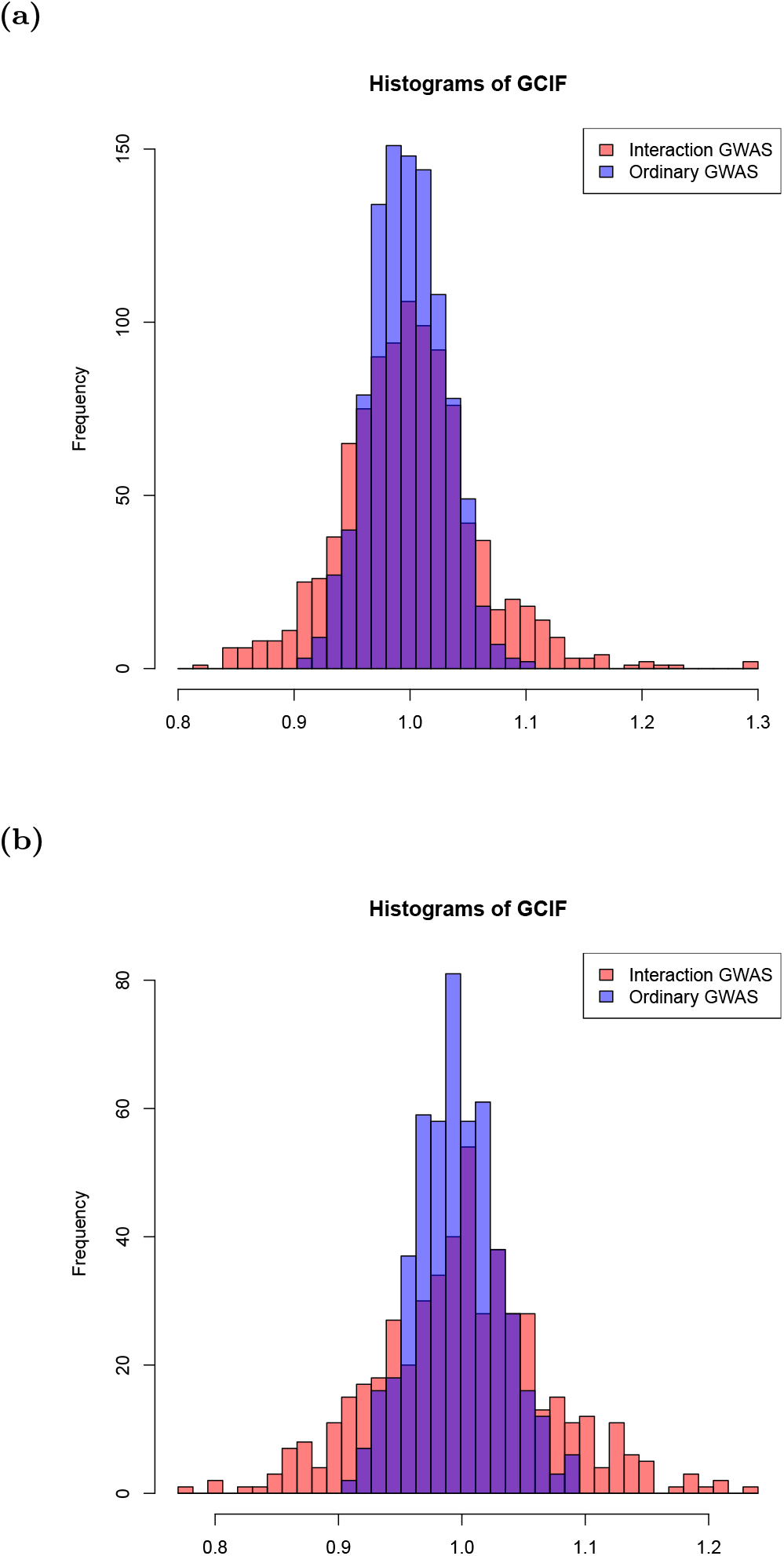
Histograms of GCIFs for interaction GWAS and for ordinary, non-interaction. GWAS Each panel is based on *r* = 5,000 simulated null GWASs in which *Y, Z* and *G* are simulated independently, with the elements of *Y* i.i.d. normal. For each GWAS, two different GCIFs are calculated, each based on *m* = 5,000 test statistics. The GCIF for ordinary (non-interaction) GWAS uses the *m* genetic association tests between *Y* and the *G*_*j*_ s, and the GCIF for interaction GWAS uses the *m* interaction tests based on Model (1). In each panel, the blue histogram represents the *r* resulting GCIFs for ordinary (non-interaction) association testing, and the red histogram represents the *r* resulting GCIFs for interaction testing. In Panel A, both *Z* and the *G*_*j*_ s are Bernoulli distributed, and in Panel B, both *Z* and the *G*_*j*_ s are Binomial(2) distributed.

In the LMM context, the test based on *T*_*j*_ is commonly called the “Wald test.” In fact, the ordinary t-test for interaction is also a Wald test, so this term is not a useful way of distinguishing the LMM-based test from the ordinary one. We refer to the test based on *T*_*j*_ as the “t-test” in both cases, and, when needed, we specify whether it is performed in an LMM or a linear model.

For both the linear and LMM cases, we define the numerator of the t-statistic to be

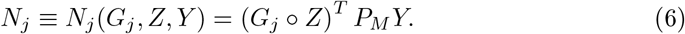

Then the interaction t-statistic in (5) can be rewritten as

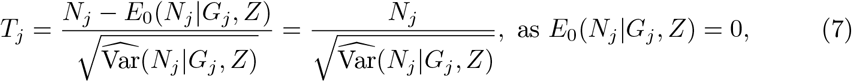

where both *E*_0_(*N*_*j*_ |*G*_*j*_, *Z*) and Var(*N*_*j*_ |*G*_*j*_, *Z*) are calculated based on Model (4), *E*_0_(*N*_*j*_ |*G*_*j, Z*) has the additional assumption *δ* = 0, and 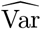_ d _denotes estimated variance._ For testing interaction in a GWAS context, we propose to replace *T*_*j*_ by a “corrected” statistic

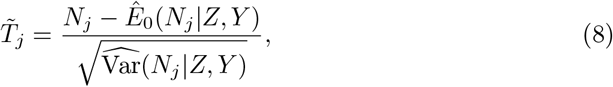

where the difference from Eq (7) is that we condition on (*Z, Y*) instead of on (*G*_*j*_, *Z*). The remaining challenge of the methods development is to obtain appropriate estimators *Ê*_0_(*N*_*j* |*Z, Y*) and 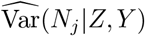_. We perform the following steps:

1. We approximate *N*_*j* by *Ñj* ≡ *Ñj*_ (*G*_*j*_, *Z, Y*), where *Ñj* is quadratic in *G*_*j*_.
2. We calculate *E*_0_(*Ñj* |*Z, Y*) and approximate Var(*Ñj* |*Z, Y*) as functions of *E*_0_(*G*_*j*_ |*Z, Y*) and Var(*G*_*j*_ |*Z, Y*).
3. We calculate *E*_0_(*G*_*j*_ |*Z, Y*) and Var(*G*_*j*_ |*Z, Y*) theoretically based on a suitable model.
4. We obtain estimates *Ê*0(*G*_*j*_ |*Z, Y*) and 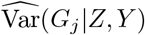 for the quantities in step 3.
5. We plug the estimates from step 4 into the expressions for *E*_0(*Ñj*_ |*Z, Y*) and Var(*Ñj* |*Z, Y*) from step 2 to obtain *Ê*_0_(*N*_*j*_ |*Z, Y*) and 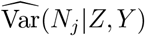, respectively, and calculate 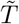 in (8).

The quadratic approximation in step 1 is is based on an asymptotic approximation and is detailed in S1 Text. The calculation of *E*_0_(*Ñj* |*Z, Y*) in step 2 is completely straightforward. To approximate Var(*Ñj* |*Z, Y*), we perform a variance calculation that is exact for the case when *Gj* |*Z, Y* has a normal distribution and is otherwise approximate (see S1 Text). Step 5 is completely straightforward given the other steps. Here, we give more details on steps 3 and 4.

For the conditional moment calculations in step 3, to model *G*_*j*_ |*Z*, we consider two different modeling approaches: a normal approximation and a discrete model. For the normal approximation, we assume a normal regression model for *G*_*j*_ |*Z*, i.e., we take *G*_*j*_ = *a*1_*n*_ + *bZ* + *η*, where 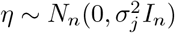, or, more generally, where *Ũ* consists of the intercept and any confounding covariates that are in *U*, we take *G*_*j* = *aŨ*_ + *bZ* + *η*, with *a, b*, and 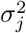 unknown. For the discrete model approach, we instead assume a discrete model for *G*_*j*_ |*Z*, where we assume that conditional on *Z*, the *n* entries of the vector *G*_*j*_, call them *G*_1*j*_, …, *G*_*nj*_, are independent with *P*(*G*_*ij*_ = *k*|*Z*_*i*_ = *z*) = *p*_*k*|*z*_ for all choices of (*i, k, z*), where these may also depend on *Ũ* as needed. Since *G*_*j*_ is a genotype, we will have *k* ∈ {0, 1, 2} when the genotypes are from a diploid organism or *k* ∈ {0, 1} when the genotypes are from a haploid organism or inbred line. For the latter case, we can use a logistic regression model for *G*_*j*_ |*Z*, and for the former case a binomial regression model. In the *A. thaliana* dataset we analyze, both *G*_*j*_ and *Z* are binary and there are no additional confounding covariates, in which case the discrete model can simply be specified in terms of the two parameters *p*_1|0_ and *p*_1|1_, without the need for a logistic model.

For the conditional moment calculations in step 3, we also consider two different modeling approaches for *Y* |(*G*_*j*_, *Z*). The first approach is to assume that Model (4) holds, which we call the homoscedastic model. The second approach assumes a more general and robust version of Model (4) in which we allow a specific type of heteroscedasticity, namely, we allow 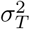 to depend quadratically on *Z*, and we call this the heteroscedastic model. In an interaction GWAS, it can potentially be important to consider this specific type of heteroscedasticity, because it arises naturally in a model in which *Z* interacts with some other variable in a linear model or LMM for *Y*, even if it doesn’t interact with *G*_*j*_ [39–41, 43]. That is, suppose the true model for *Y* could be written

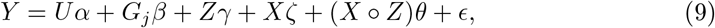

where *Y, U, α, G*_*j*_, *β, Z, γ*, and *ϵ* are as before, *ζ* and *θ* are unknown scalar coefficients, and *X* is some additional variable that might or might not be observed, is independent of (*G*_*j*_, *Z*), and that interacts with *Z*. In other words, from the point of view of testing for interaction between *Z* and *G*_*j*_, this is a null model, but it allows for the possibility that *Z* does interact with some other variable, *X*, such as a SNP on another chromosome, or a non-genetic variable. Then in this model, if we calculate Var(*Y* |*G*_*j*_, *Z*), we find that it depends on *Z* quadratically. In other words, we have the specific type of heteroscedasticity described above. This motivates the heteroscedastic model for *Y* |(*G*_*j*_, *Z*).

Given the modeling assumptions described above, we now consider the calculation of *E*_0_(*G*_*j*_ |*Z, Y*) and Var(*G*_*j*_ |*Z, Y*) in step 3. When the normal approximation is used for *G*_*j*_ |*Z*, then with either the homo- or heteroscedastic model for *Y* |(*G*_*j*_, *Z*), we obtain a multivariate normal distribution for (*G*_*j*_, *Y*)|*Z*, from which *E*_0_(*G*_*j*_ |*Z, Y*) and Var(*G*_*j*_ |*Z, Y*) can be easily computed using standard properties of multivariate normal. When a discrete model is used for *G*_*j*_ |*Z*, then with either the homo- or heteroscedastic model for *Y* |(*G*_*j*_, *Z*), we can apply a Bayes rule calculation to obtain the discrete distribution Pr(*G*_*j*_ |*Z, Y*). For example, if we assume unrelated individuals, then conditional on (*Z, Y*), *G*_1*j*_, … *G*_*nj*_ are independent with

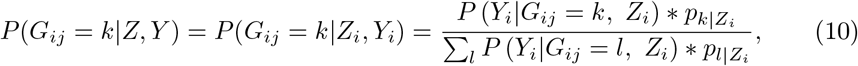

where *P* (*Y*_*i*_|*G*_*ij*_ = *k, Z*_*i*_) is a univariate normal density function.

### Approximate null conditional mean and variance of interaction t-statistic numerator

To better understand the surprising behavior of the t-statistic for interaction in a GWAS setting under the null hypothesis, it can be helpful to examine approximate analytical formulas for the null conditional mean and variance of the t-statistic numerator given (*Z, Y*), where *Z* and *Y* are the variables that remain fixed for the GWAS. If we instead took the more common approach of conditioning on (*G*_*j*_, *Z*), we would obtain zero for the null conditional mean and 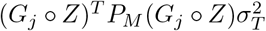 for the null conditional variance.

When we use the normal approximation for *G*_*j*_ |*Z*, use a linear model for *Y* instead of an LMM, and assume no covariates, then it becomes possible to obtain approximate analytical formulas for *E*_0(*Ñ*_|*Z, Y*) and Var_0(*Ñ*_|*Z, Y*). We obtain

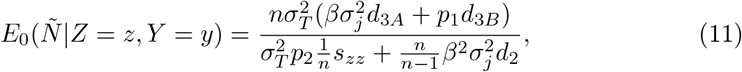

Where 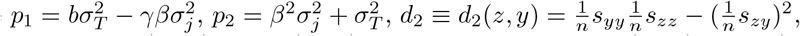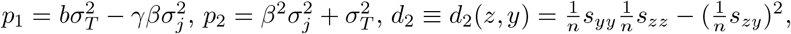 and where for any 3 vectors *u, v* and *w* of length *n*, we define 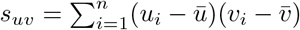 and 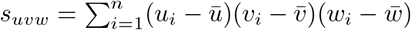.

The motivation for this notation is that “p” denotes “parameters”, and *p*_1_ and *p*_2_ are functions only of parameters; “d” denotes data, and the subscript “2” in *d*_2_ denotes that *d*_2_ is a function of only the observed sample second moments of (*Z* = *z, Y* = *y*) and not of any parameters. The subscript “3” in *d*_3*A*_ and *d*_3*B*_ denotes that they are functions of only the observed sample third and second moments of (*Z* = *z, Y* = *y*) and not of any parameters. In the special case when *β* = 0, we get

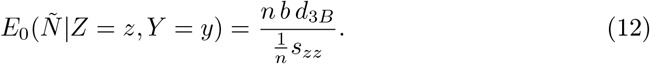

These approximate formulas can serve as useful heuristics about when the null conditional expectation of the interaction t-statistic might or might not be approximately zero. From this approximation, we get that if both *β* and *b* are 0, which would happen if *G*_*j*_ is independent of (*Z, Y*), then the null conditional expectation should be 0. We can also see that if (1) (*Y, Z*) is multivariate normal with arbitrary correlation or (2) *Y* and *Z* are independent (with any distribution), then in sufficiently large samples, *d*_3*A*_ and *d*_3*B*_ will both be close to 0, so we expect the null conditional mean to be close to 0 in sufficiently large samples. However, if *Y* is heteroscedastic with respect to *Z*, or if *Z* has a skewed distribution and *Z* and *Y* are correlated, then the null conditional mean could be non-zero when *G*_*j*_ is correlated with *Z* or *Y*, even in large samples.

Using the normal approximation, the approximate null conditional variance is

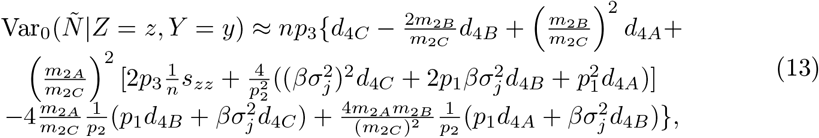

where 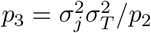 (with *p*_2_ as defined in Eq (11)) is a function of only parameters, *d*_4*A*_, *d*_4*B*_, and *d*_4*C*_ are functions of only the observed sample 4th and second moments of (*Z* = *z, Y* = *y*), with 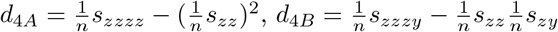 and 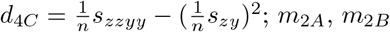, and *m*_2*C*_ are “mixed” terms that are functions of both parameters and data, but that depend on the data only through the observed sample 2nd moments of (*Z* = *z, Y* = *y*), with 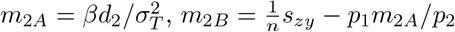 and 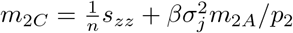. When *β* = 0, we further get

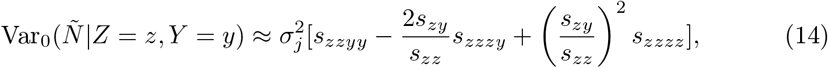

which is almost exclusively a function of the observed second and fourth sample moments of (*Z* = *z, Y* = *y*), except for the parameter 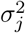.

The above formulas can be useful as heuristics, but when *G*_*j*_ has a discrete distribution, we instead use a discrete model for *G*_*j*_ |*Z*, and the null conditional mean and conditional variance based on that do not lend themselves to a simple closed-form expression. Furthermore, with covariates or in a LMM, the results are also more involved. Finally, the variance expression we give above is the one we obtain in the special case when we assume *δ* = 0, and, more generally, we usually prefer to do a Wald test, in which case we need an estimate of the conditional variance under the alternative model, which is also a more involved calculation.

### Estimation step

In step 4, we need to obtain estimates *Ê*0(*G*_*j*_ |*Z, Y*) and 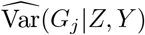 of the quantities we derived theoretically in step 3. In the case that the model for *Y* |*G*_*j*_, *Z* is the linear, homoscedastic model with Σ = *I*_*n*_, then when we use the normal approximation for *G*_*j*_ |*Z*, we can fit ordinary least squares (OLS) regression of *G*_*j*_ on (*U, Z, Y*) and use the fitted values as 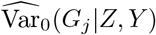 and RSS*/*(*n* − *k* − 2) as Var d_0_(*G*_*j*_ |*Z, Y*). Similarly, when *Y* |*G*_*j*_, *Z* is the linear, homoscedastic model with Σ = *I*_*n*_, *G*_*j*_ |*Z* is the discrete model, and *G*_*j*_ is binary (or binomial), we can use logistic (or binomial) regression of *G*_*j*_ on (*U, Z, Y*) to obtain *Ê*_0_(*G*_*j*_ |*Z, Y*) and 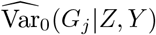. However, to obtain 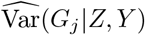 under the alternative (which can allow us to do a more powerful Wald-type test instead of a score-type test), and for all other modeling cases, we instead use some version of the Bayes rule calculation, where we fit the model for *G*_*j*_ |*Z* to obtain its parameters, fit the model for *Y* |*G*_*j*_, *Z* to obtain its parameters, and then plug the estimated parameters into the Bayes rule calculation. To allow for heteroscedasticity in step 4, we first regress *G*_*j*_ out of *Y* to obtain *r*_*Y* |*Gj*_, and then we replace *Y* by *r*_*Y* |*Gj*_ in the model for interaction, and allow for heteroscedasticity in the model, where the variance can depend on *Z* and *Z*^2^.

### Diagnostic ratio for identifying GWISs in which the “feast or famine” effect occurs

When we consider how to predict the likely “feast or famine” effect for a GWAS based on a given (*Y, Z*) observed in data, a natural starting point is to consider the mean and variance adjustments to the t-statistic obtained using the TINGA method. These adjustments change with the variant *G*_*j*_ as we scan the genome, and they depend on some modeling choices. For the diagnostic, we could choose rather simple modeling assumptions (normal regression models for *Y* on (*G*_*j*_, *Z*) and for *G*_*j*_ on *Z*) and consider a hypothetical *G*_*j*_ that is completely independent of (*Y, Z*), so that the mean adjustment reduces to 0, and we are left with only a variance adjustment *V* that is now a function of various sample second and fourth central moments (including cross-moments) of (*Y, Z*). In this case, the ratio of *V* to the square of the denominator of the t-statistic, call it *R*, would seem to be a natural measure of the feast or famine effect (and the TINGA adjustment to the test statistic would be to multiply the t-statistic by *R*^−1*/*2^), where large *R* would be indicative of “feast” and small *R* of “famine”. However, the denominator of the t-statistic also involves *G*_*j*_, so we approximate it by taking a first-order approximation to its conditional expectation given (*Y, Z*) under the simple modeling assumptions we chose above.

The resulting ratio *R* is given by

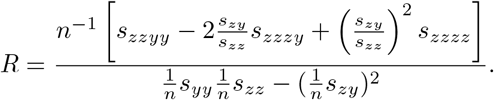

This ratio turns out to have a direct interpretation as one of the 5 components of co-kurtosis between *Z* and *r*_*Y* |*Z*_, where *r*_*Y* |*Z*_ denotes the vector of residuals from simple linear regression of *Y* on *Z*. If we abbreviate *r*_*Y* |*Z*_ as *r*, then we have that *R* = *n* ∗ *s*_*zzrr*_*/*(*s*_*zz*_*s*_*rr*_). Informally, the sample co-kurtosis is generally considered to measure the extent to which the variables *z* and *r* are observed to have extreme observations or outliers, and whether the extreme observations for *z* and *r* tend to co-occur in the same individuals. However, as we show in the **Results** section, even just the ordinary sampling variability in this ratio that occurs in different (*Z, Y*) replicates is highly predictive of the FoF effect for an interaction GWAS using the given (*Z, Y*). Note that the diagnostic ratio does not contain any information on interaction between *Z* and and any specific *G*_*j*_.

For the case when there are additional covariates *U* in the linear model for *Y*, beyond just intercept, *G*_*j*_, and *Z*, we can extend the definition of *R* to

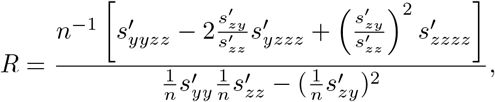

where 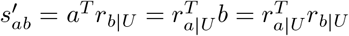 is the residual of *a* after regressing out *U* where *U* consists of the intercept and any additional covariates but does not include *G*_*j*_ or *Z*, and where 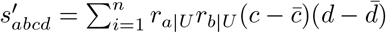.

### Additional methodological considerations

In the special case when at least one of *Z* and *X* is discrete, it is natural to place certain constraints on when one would or would not perform any sort of interaction test. For example, if both *Z* and *X* are binary and are perfectly correlated, then there would typically be zero information in the data on interaction between them as a predictor of *Y*, and if they are almost perfectly correlated, then the amount of information available on interaction would be quite low. In the case when *Z* and *X* are both binary, we can think of constructing a 2 *×* 2 table of counts of the four possible observed values of (*Z, X*) in the data, and we require the minimum cell count (MCC), i.e., the smallest of the counts of the four possible observed values, to be at least 5 in order to perform the interaction t-test.

Step 4 of the TINGA method requires some additional parameter estimation compared to the interaction t-test. If all variables were continuous, then with typical GWAS sample sizes, the estimation of a handful of additional parameters would pose little problem for the inference. When *X* and *Z* are both binary, however, then we require MCC ≥ 20 in order to perform the additional estimation in step 4. Therefore, our TINGA method uses a mixed strategy in that case, in which, when 5 ≤ MCC *<* 20 we use the interaction t-test, and when MCC ≥ 20, we use the adjustment strategy. All the TINGA results for the case when both *X* and *Z* are binary use this “mixed” strategy.

Specifically for the problem of epistasis detection, it has been noted that in the presence of an untyped causal variant, two typed variants in strong linkage disequilibrium that form a haplotype that tags the untyped variant could exhibit false epistasis [34]. Therefore, in detection of epistasis, we only test for epistasis between variants *X* and *Z* if their sample correlation is close to 0. (In our data analysis we use a cut-off of .1 for absolute value of correlation.)

For the problem of epistasis detection, for a given pair (*G*_1_, *G*_2_) of SNPs, there are two possible adjustments, one based on conditioning the test on (*G*_1_, *Y*) and the other based on conditioning the test on (*G*_2_, *Y*). We propose the strategy of conditioning on the less polymorphic of *G*_1_ and *G*_2_, because that should result in more information available for the statistical test leading to a more powerful test. We test this strategy in simulations.

## Simulations

### Type 1 error simulations I

For the first set of type 1 error simulations, we simulate 10^5^replicate GWASs each with *n* = 10^3^individuals. In each replicate, we simulate the elements of *Z* as *n* i.i.d draws from Bin(2, 0.2), then center and standardize them, and we assume that *Z* explains 1% of the variance of the trait. Among a much larger set of *m* SNPs in the GWAS, we assume that there are 149 associated SNPs, consisting of 49 SNPs that each explain 1% of the variance of the trait and 100 SNPs that each explain 0.5% of the variance of the trait. Conditional on allele frequency *p*_*j*_, the genotype *G*_*j*_ of the *j*th SNP is simulated as *n* i.i.d. draws from Bin(2, *p*_*j*_), where *p*_1_, …, *p*_*m*_ are i.i.d. Uniform(0.2,0.8), and where SNPs are taken to be unlinked. Then each *G*_*j*_ is centered and standardized based on *p*_*j*_ (i.e., 2*p*_*j*_ is subtracted off and the result divided by ^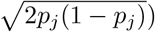^. Then *Y* is simulated as

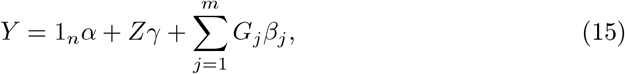

where 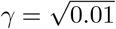, exactly 49 of the *β*_*j*_ ‘s have magnitude |*β*_*j*_ | equal to ^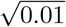^ with a random sign (positive or negative) given to each *β*_*j*_, exactly 100 of the *β*_*j*_ ‘s have magnitude 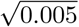with a random sign, and the remaining *β*_*j*_ ‘s are 0. (The value of the overall mean *α* has no impact and can be taken to be 0.) In each replicate, we calculate the diagnostic ratio based on (*Y, Z*), and for each *G*_*j*_, *j* = 1, …, *m*, we test for interaction between *Z* and *G*_*j*_ in the model of Eq 1 using (i) the usual t-test for interaction (ii) a TINGA-adjusted t-test in which we condition on (*Y, Z*) and (iii) the HC3 method [44], which is a heteroscedasticity-corrected t-test.

### Power simulations I

For the first set of power simulations, we simulate 10^6^replicate GWASs each with *n* = 10^3^individuals. In each replicate, we simulate *Z* and *G*_*j*_ *j* = 1, … *m* as above. Then *Y* is simulated as

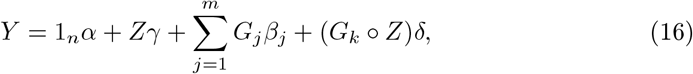

where *α, γ* and *β*_*j*_ are as above, *k* is the index of one of the SNPs for which 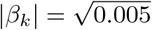, and 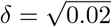. In each replicate, we calculate the diagnostic ratio based on (*Y, Z*), and we test for interaction between *Z* and *G*_*k*_ in the model of Eq 1 using (i) the usual t-test for interaction and (ii) a TINGA-adjusted t-test in which we condition on (*Y, Z*).

### Replicability

To estimate replicability as a function of the diagnostic ratio, we first binned the observed diagnostic ratios from the type 1 error and power simulations. For simulated GWASs with diagnostic ratio in the *i*th bin, we calculated replicability as

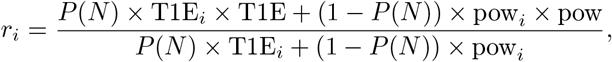

where *P*(*N*) is the probability that a tested interaction is a true null, which we took to be 0.9998, T1E_*i*_is the empirical type 1 error at level 10^−4^ observed for the simulated GWASs with diagnostic ratio in the *i*th bin, pow_*i*_is the empirical power observed for the simulated GWASs with diagnostic ratio in the *i*th bin, T1E is the overall average type 1 error at level 10^−4^, and pow is the overall average power. Thus, *r*_*i*_ represents the probability that an interaction detected for a GWAS with diagnostic ratio in the *i*th bin would be replicated in another randomly chosen GWAS.

### “Feast” effect due to *Z* being associated with a causal variant: example of Hemani et al. 2014 and 2021

Hemani et al. 2014 [45] studied epistasis in expression traits in humans, identifying 501 significant pairwise interactions between common SNPs affecting expression of 238 genes. In the case of some of these significant pairwise interactions, it was later noted that a third SNP associated with the trait could explain all of the pairwise interaction. [37] While this type of effect is well-known [34, 37] in the case when the two putatively interacting SNPs are proximal and in LD with one another, Hemani et al. 2021 [38] found that this effect had occurred in a large number of cases in which one of the two putatively interacting SNPs was cis to the gene whose expression was being influenced, the other putatively interacting SNP was on a different chromosome, and the third SNP whose effect could explain the interaction was cis to the gene and in LD with the first SNP, a type of effect which was unexpected [38]. Motivated by these findings, Hemani et al. 2021 performed simulations in which an untyped SNP *W* is causal for *Y*, and an observed SNP *Z* is in LD with *W* and is conditionally independent of *Y* given *W. Z* is tested for interaction with SNPs *G*_*j*_, *j* = 1, …, *m*, none of which are on the same chromosome with *Z*. In the simulations, they find the effect we have called the “feast” effect, i.e., the genomic control inflation factor and type 1 error of the GWIS varied across replicates and were on average inflated. They report, “Here we show… a previously unrecognized property of the gold-standard statistical test to detect interactions, namely that the presence of imperfectly tagged additive causal variants can lead to phantom epistasis between unlinked markers. Therefore, the false positive rate in studies that use the test may not be sufficiently controlled and, to our knowledge, no current statistical fix exists for this problem.”

We perform simulations similar to those of Hemani et al. 2021. Specifically, we simulate 100 GWAS replicates, each based on *n* = 10^3^individuals. In each replicate, we take the haplotype frequencies of (*W, Z*) to be those of (rs67903230,rs13069559) observed in the data of Hemani et al. 2014, and we take *m* = 10^4^. Following Hemani et al. 2021, we sample the proportion of variance of *Y* explained by *W* from Uniform(0,.5) and test for interaction between *Z* and each *G*_*j*_, *j* = 1, …, *m*. For each replicate, we calculate the diagnostic ratio based on (*Y, Z*), the *m* p-values for interaction based on the ordinary t-test, and the *m* p-values for interaction based on the TINGA-adjusted t-test.

### Type 1 error and power simulations II

In the next set of simulations, allow for a linear mixed model for *Y* instead of a linear model, and we consider the case where both *Z* and *G*_*j*_ are Bernoulli distributed, because that is the situation in the *A. thaliana* dataset. We compare the performance of the t-test and our methods in 3 simulation settings.

### Non-GRM case

In each replicate, we simulate a Bernoulli *Z* and *m* = 4 Bernoulli 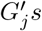 for *n* = 1000 independent individuals, and simulate *Y* under the alternative model 17

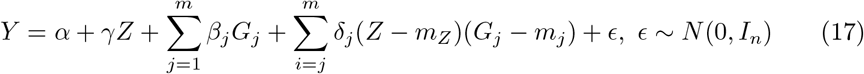

where *Z, G*_2_, *G*_3_, *G*_4_ have marginal effects on *Y* and only *G*_4_ has interactive effect with *Z* on *Y* (setting 3 in S1 Text).

### GRM case 1: unrelated individuals; accounting for additive polygenic effects

In this case, *Z, G*_*j*_ ‘s are simulated in the same way as GRM case 1. *Y* is simulated with the same model 17, except a GRM as an extra variance component 18

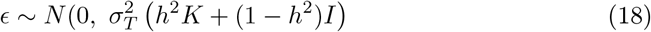

(setting 4 in S1 Text).

### GRM case 2: population structure with 3 sub-populations

We also tried the setting with population structure in which there are 3 sub-populations. See setting 11 in S1 Text. Both *Z* and *G*_*j*_ ‘s are still Bernoulli distributed, and simulated with the 3 sub-populations. *Z, G*_2_, *G*_3_, *G*_4_ and (*Z ° G*_4_) have effects on *Y*. *Y* also has indicators of the sub-populations as covariates.

## Results

### Simulations

#### Diagnostic ratio predicts the “feast or famine” effect

In Fig 3, type 1 error of the ordinary t-test and of the TINGA-corrected t-test are plotted against the diagnostic ratio. The plot clearly demonstrates the “feast or famine” effect, i.e., the type 1 error of the t-test varies systematically and significantly across GWISs, with some GWISs having type 1 error that is too small and others having type 1 error that is too large, where this effect can be accurately predicted in advance by the diagnostic ratio calculated from (*Y, Z*), without using any information on the genotypes *G*_1_, …, *G*_*m*_,

#### “Feast or famine” adversely affects type 1 error, power, and replicability

In Fig 4, power of the t-test and of the TINGA-adjusted t-test are plotted against the diagnostic ratio, while in Fig 5, the replicability of significant interaction results detected using the t-test and detected using the TINGA-adjusted t-test are plotted against the diagnostic ratio. From the results, we can see that for the ordinary t-test for interaction, the “feast or famine” effect results in systematic inflation or deflation of type 1 error ascross SNPs within a GWAS (Fig 3), reduced power in the famine GWASs (Fig 4), and lack of replicability of interaction detected in the feast GWASs (Fig 5), where an excessively low diagnostic ratio predicts deflated type 1 error and reduced power for the GWAS and an excessively high diagnostic ratio predicts inflated type 1 error and reduced replicability of the detected interaction.

**Fig 3.**
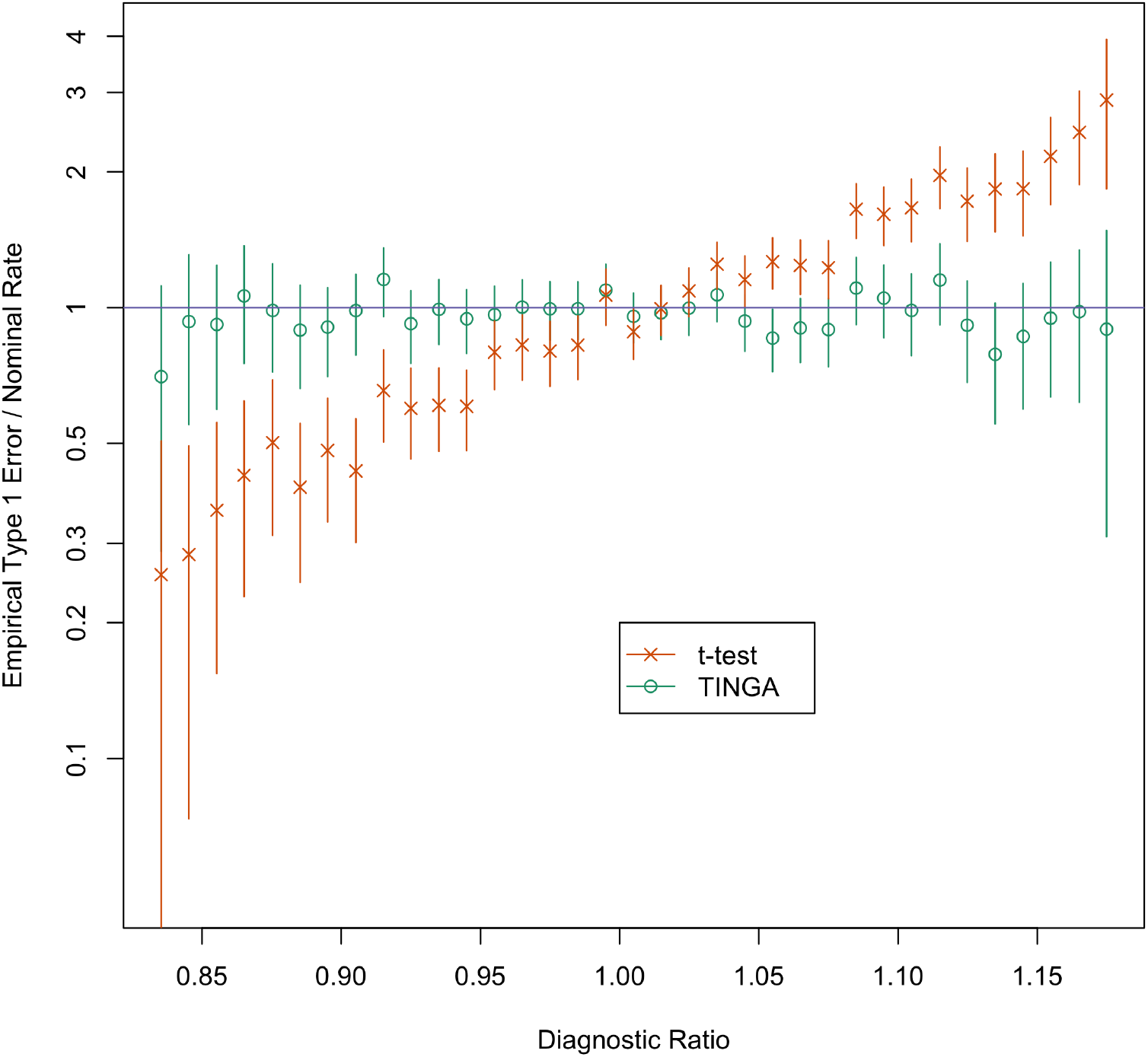
Empirical type 1 error vs. diagnostic ratio for the standard t-test for interaction and the TINGA adjusted t-test for interaction. 10^5^simulated GWAS’s are divided into bins based on their observed diagnostic ratio, and the average type 1 error in the bin is plotted against the average diagnostic ratio in the bin, for the standard t-test and the TINGA adjusted t-test that conditions on (*Z, Y*). Each vertical line segment represents a 95% confidence interval for the type 1 error in the bin, for the given testing method. The y-axis is empirical type 1 error at nominal level 10^−4^ divided by 10^−4^, so, e.g., a y-value of 2 corresponds to an empirical type 1 error of 2e-04. The y-axis is logarithmically scaled.

#### Conditioning on (*Y, Z*) can correct the “feast or famine” effect

In Fig 3, we can see that the type 1 error of the TINGA-adjusted t-test does not vary significantly from the nominal level across GWISs. Similarly, the power (Fig 4) and replicability (Fig 5) appear to stay approximately constant across GWISs when the TINGA-adjusted t-test is used. This shows that using appropriate conditioning (i.e., conditioning on (*Y, Z*) instead of on (*Z, G*_*j*_)) in the analysis effectively eliminates the “feast or famine” effect, as we predicted theoretically.

**Fig 4.**
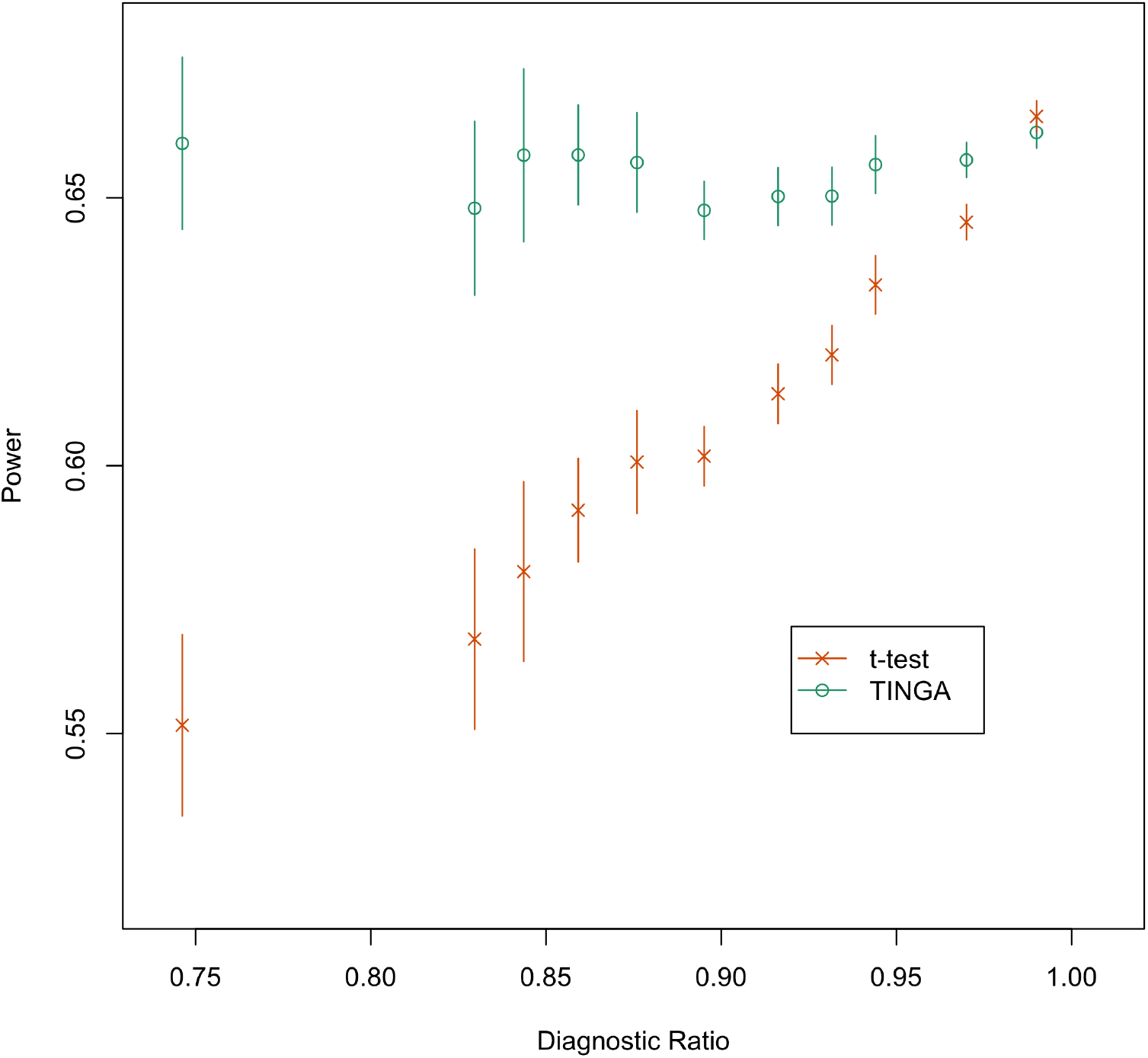
Power vs. diagnostic ratio for the standard t-test for interaction and the TINGA adjusted t-test for interaction. 10^6^simulated GWASs are divided into bins based on their observed diagnostic ratio, and the average observed power at level 1e-04 in the bin is plotted against the average diagnostic ratio in the bin, for the standard t-test and the TINGA adjusted t-test that conditions on (*Z, Y*). Each vertical line represents a 95% confidence interval for the average power of the GWASs with diagnostic ratios in the given bin, based on the given testing method.

#### “Feast” effect due to *Z* being associated with a causal variant: example of Hemani et al. 2014 and 2021

The example of Hemani et al. 2014 and 2021 demonstrates a type of model misfit that can lead to a particularly strong “feast” effect for the standard interaction t-test. In Fig 6, we can see that the smallest p-values based on the t-test can be more than 6 orders of magnitude too small. In Figs 7 and 8, we can see that the genomic control inflation factor based on the t-test can range all the way up to 3.5 in our simulations. In Fig 9, we can see that the type 1 error (at nominal level 10^−3^) for the t-test in a given GWIS can range as high as 0.07 or higher across the simulated GWISs. These results are in close agreement with previous work [38].

**Fig 5.**
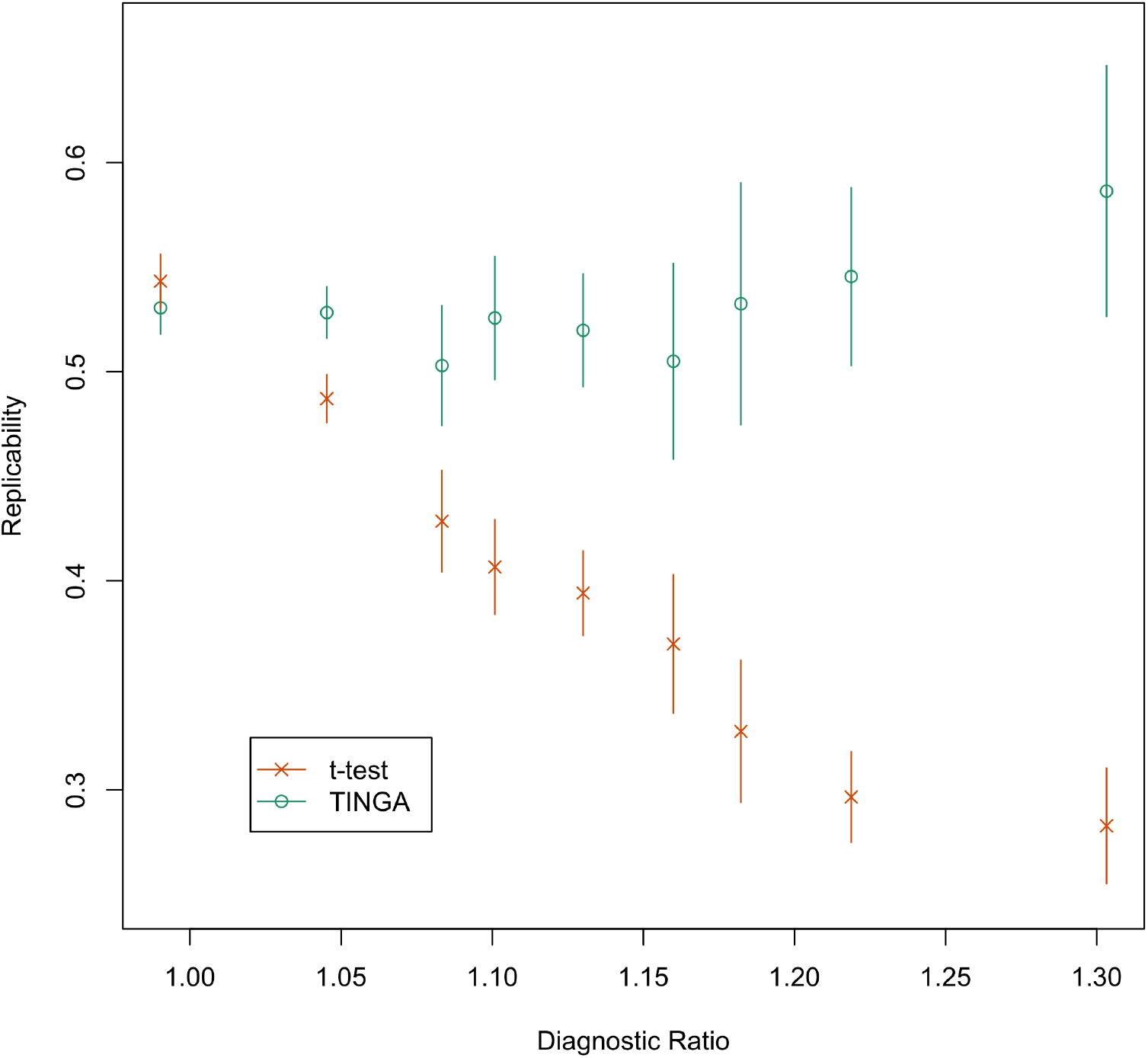
Replicability vs. diagnostic ratio for the standard t-test for interaction and the TINGA adjusted t-test for interaction. 10^6^simulated GWASs are divided into bins based on their observed diagnostic ratio, and the average replicability at level 10^−4^in the bin is plotted against the average diagnostic ratio in the bin, for the standard t-test and the TINGA adjusted t-test that conditions on (*Z, Y*). Replicability for a given testing method in a given GWAS is defined as the probability that a significant result at level 10^−4^in the given GWAS would be replicated at level 10^−4^in an independent GWAS. Each vertical line represents a 95% confidence interval for the average replicability of GWASs with diagnostic ratios in the given bin, based on the given testing method.

Furthermore, we can see that the magnitude of the “feast” effect is very accurately predicted by the diagnostic ratio, with the diagnostic ratio itself appearing to be an unbiased estimated of the genomic control inflation factor of the t-test (Fig 8), while the type 1 error of the t-test also appears to be a monotonic function of the diagnostic ratio (Fig 9).

Finally, the results show clearly that conditioning on (*Y, Z*) corrects the feast effect. In Fig 6, the TINGA-adjusted t-statistic can be seen to have correctly calibrated p-values under the null hypothesis. In Fig 7, the genomic control inflation factor of the TINGA-adjusted t-statistic appears to have a symmetric distribution closely centered on 1, which is the expected behavior under the null hypothesis. In Fig 8, the genomic control inflation factor of the TINGA-adjusted t-tstatistic appears to be close to 1 for all observed values of the diagnostic ratio. In Fig 9, the type 1 error of the TINGA adjusted t-statistic is seen to be not significantly different from the nominal, across all observed values of the diagnostic ratio.

**Fig 6.**
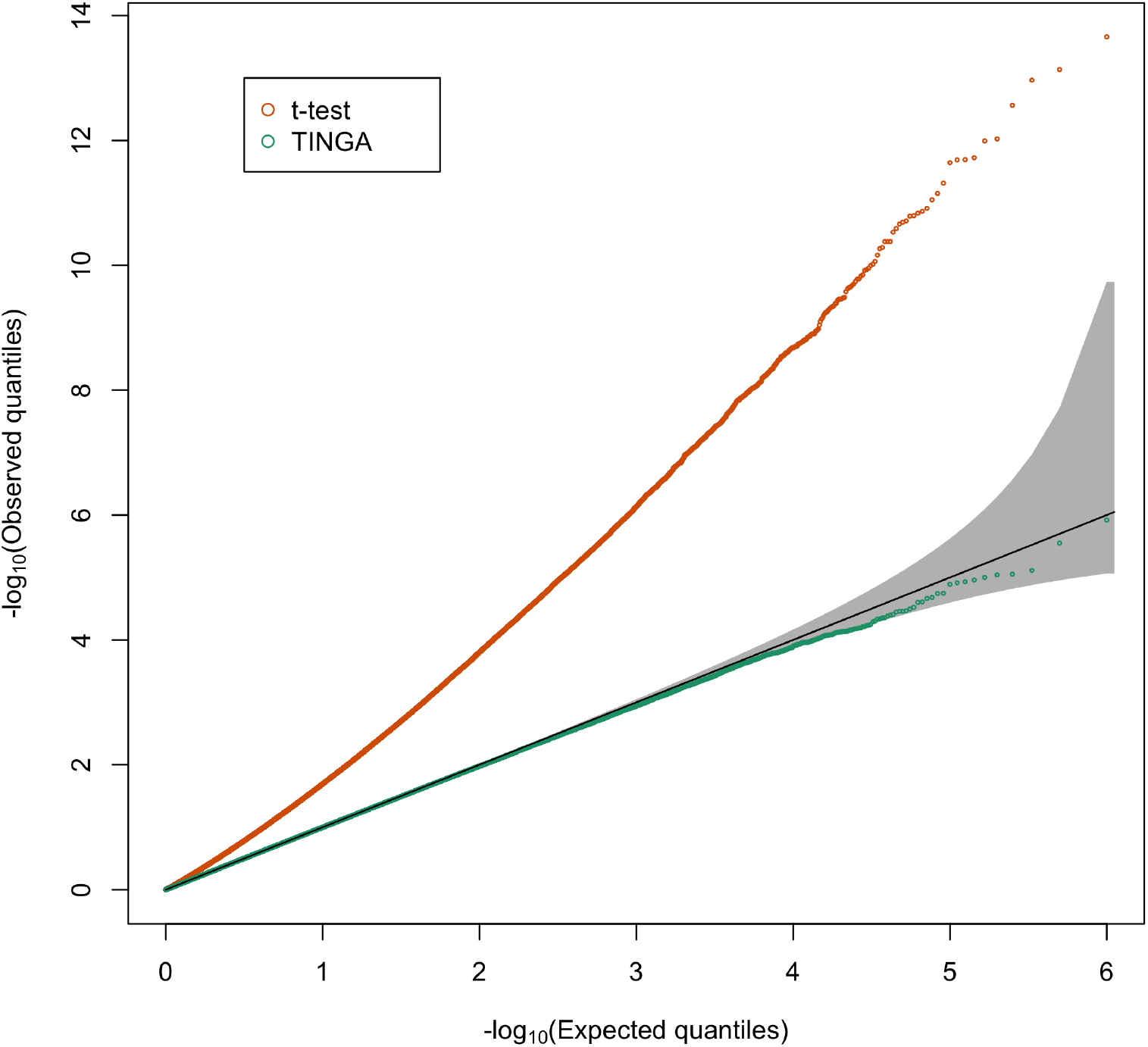
QQ-plot of p-values for the standard t-test for interaction and the TINGA adjusted t-test for interaction when *Z* is associated with a causal variant. Each QQ-plot is based on 10^6^interaction p-values from 100 GWISs simulated under the null hypothesis of no interaction. The shaded region represents the 95% acceptance region based on equal local levels [42] for a test of the null hypothesis that the p-values are i.i.d. Unif(0, 1) under the null hypothesis.

#### Type 1 error and power simulations II

We run the each of the 3 simulation settings (Non-GRM, GRM case 1 and GRM case 2) 5000 times independently to mimic 5000 independent GWASs. For *G*_1_, *G*_2_, *G*_3_, we test at level 0.05. For 5000 replicates, the 95% confidence interval is (0.0440, 0.0560). The results are in Table 2. Since the type I error rates are obtained across multiple GWASs, both uncorrected and corrected have reasonable type I error.

**Fig 7.**
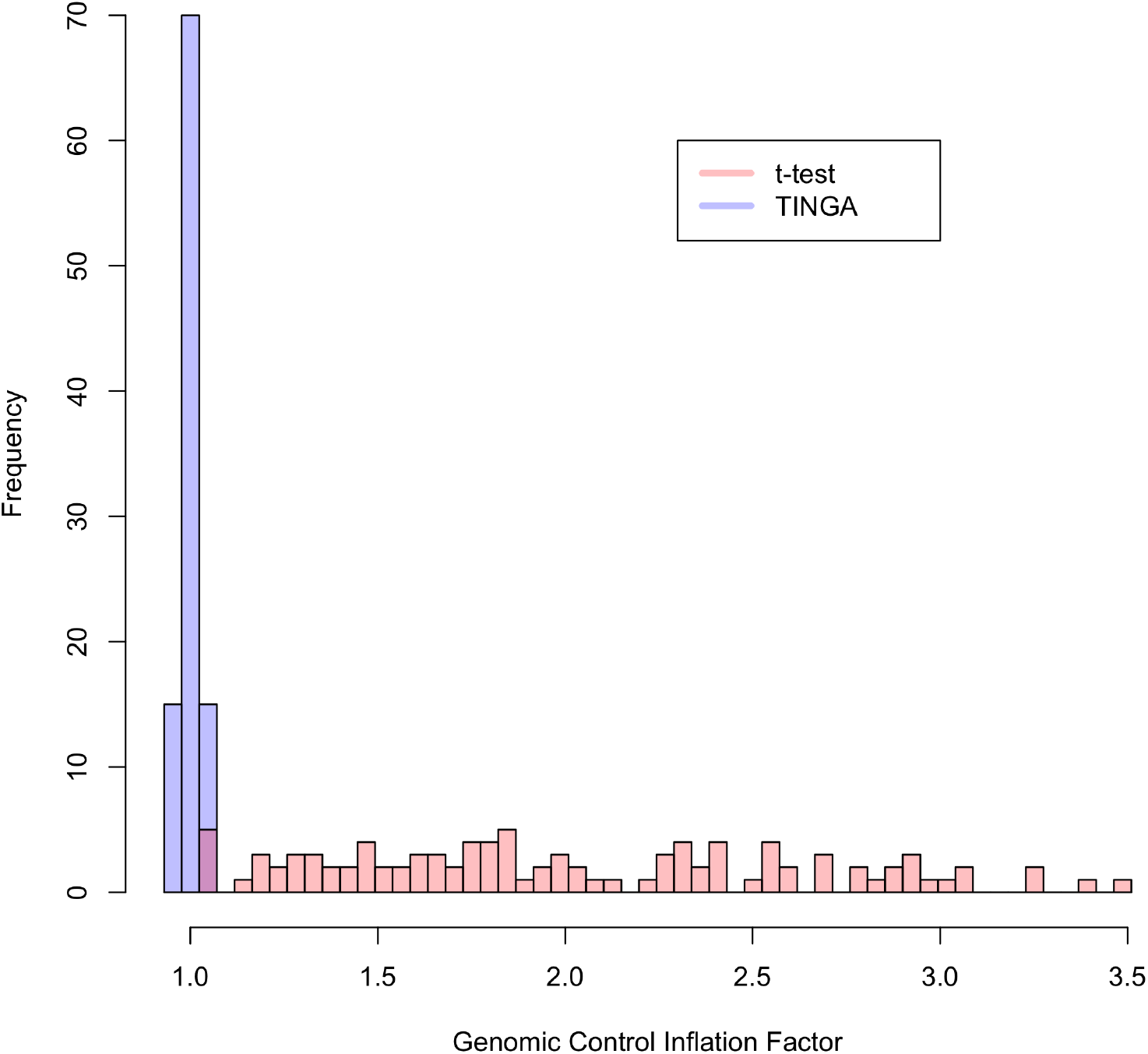
Histogram of genomic control inflation factors for the standard t-test for interaction and the TINGA adjusted t-test for interaction when *Z* is associated with a causal variant. Each histogram is based on 100 GWISs simulated under the null hypothesis of no interaction.

Table 3 compares the power of uncorrected and corrected methods for detecting interaction between *G*_4_ and *Z*. Fig 10 are plots of the power curves for the first two simulations. We can see that TINGA consistently has higher power than the unadjusted approach. For the results when *Z* and *G*_*j*_ have other distributions see S1 Text.

#### Simulation under null: check p-values within a GWAS

In this part, we consider the distribution of the GCIF within a GWAS. For each replicate GWAS, the sample size is 1000, and we simulate *Z* and *m* = 5000 *G*_*j*_ ‘s independently. We consider the following cases: (1) Both *Z* and *G*_*j*_ ‘s are Bernoulli, and *Y* is simulated under a linear model (setting 1 in S1 Text). (2) Both *Z* and *G*_*j*_ ‘s are Binomial(2), and *Y* is simulated under a linear model (setting 1 in S1 Text). (3) Both *Z* and the *G*_*j*_ ‘s are normal, and *Y* is simulated under a linear model. (4) Both *Z* and *G*_*j*_ ‘s are Bernoulli, and *Y* is simulated under a LMM (setting 2 in S1 Text). Then for every GWAS, we calculate a GCIF based on the p-values for interaction. Fig 11 gives the histograms of the resulting GCIFs. From this we can see that our methods make the GCIF much more concentrated around 1.

**Fig 8.**
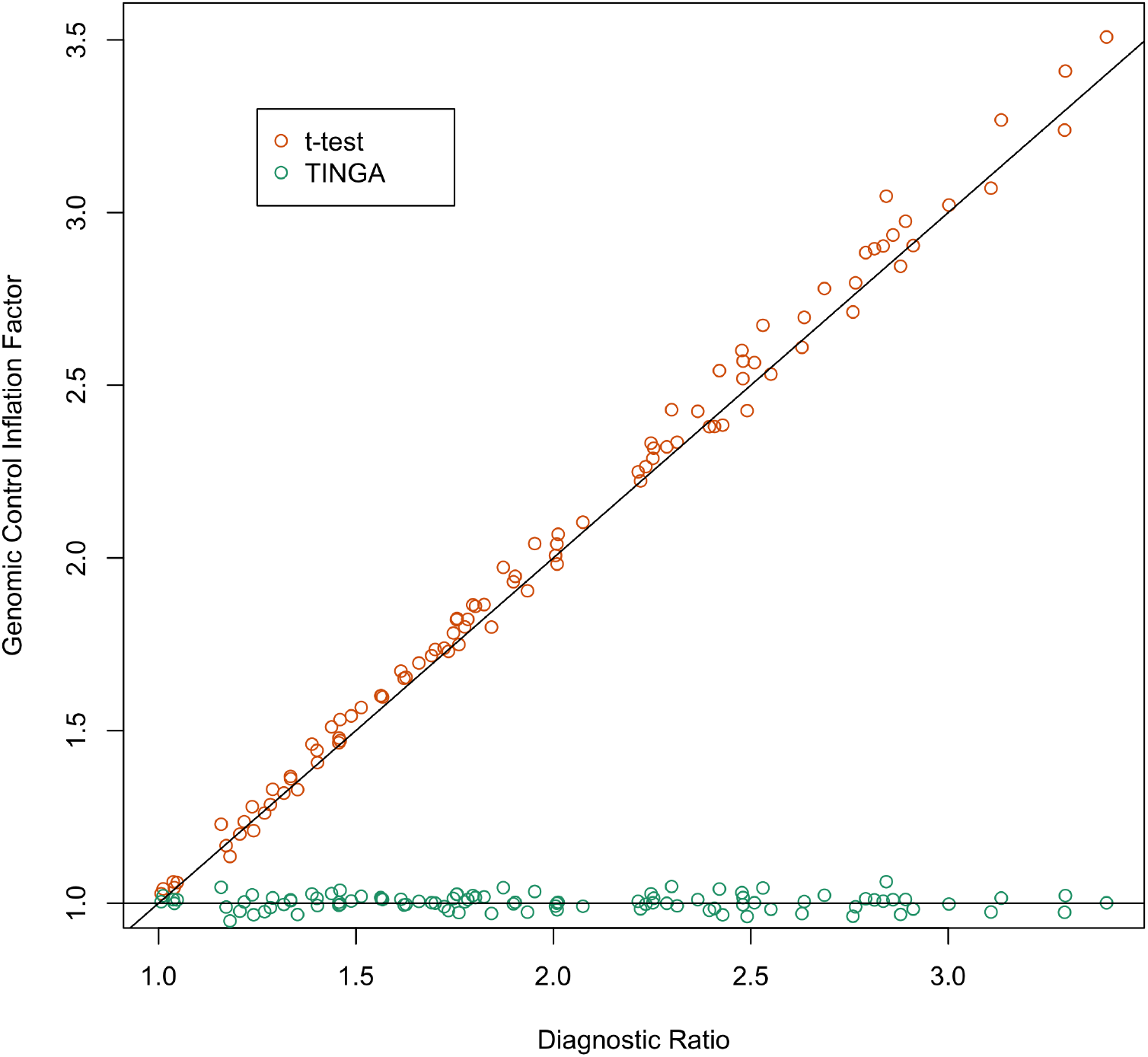
Genomic control inflation factor vs. diagnostic ratio for the standard t-test for interaction and the TINGA adjusted t-test for interaction when *Z* is associated with a causal variant. Results from 100 simulated GWISs are plotted under the null hypothesis of no interaction. The lines y=x and y=1 are shown in black.

#### Switch role of *Z* and *G*_*j*_

As described above, for the case of detecting epistasis between a pair of genetic variants, there could be two possible ways to apply TINGA. We have proposed the strategy of conditioning on the less polymorphic of the two variants (i.e., the one with the smaller minor allele frequency), because we expect that it should result in more information available for the statistical test, leading to a more powerful test. We design a simulation to test this idea. In each of 1,000 replicates, we simulate one variant with MAF .07 and another with MAF .25, independently, and we simulate *Y* according to simulation setting 5 in S1 Text. We then test interaction between the two variants using TINGA with (1) *Z* being the variant with smaller MAF and *G*_*j*_ being the variant with larger

MAF and (2) the reverse (*Z* being the variant with larger MAF and *G*_*j*_ being the variant with smaller MAF). Fig 12 is a scatterplot of the resulting p-values on the -log10 scale. This verifies our intuition that it is a more powerful strategy to condition on the variant with the smaller MAF, so we employ this strategy in the data analysis.

**Fig 9.**
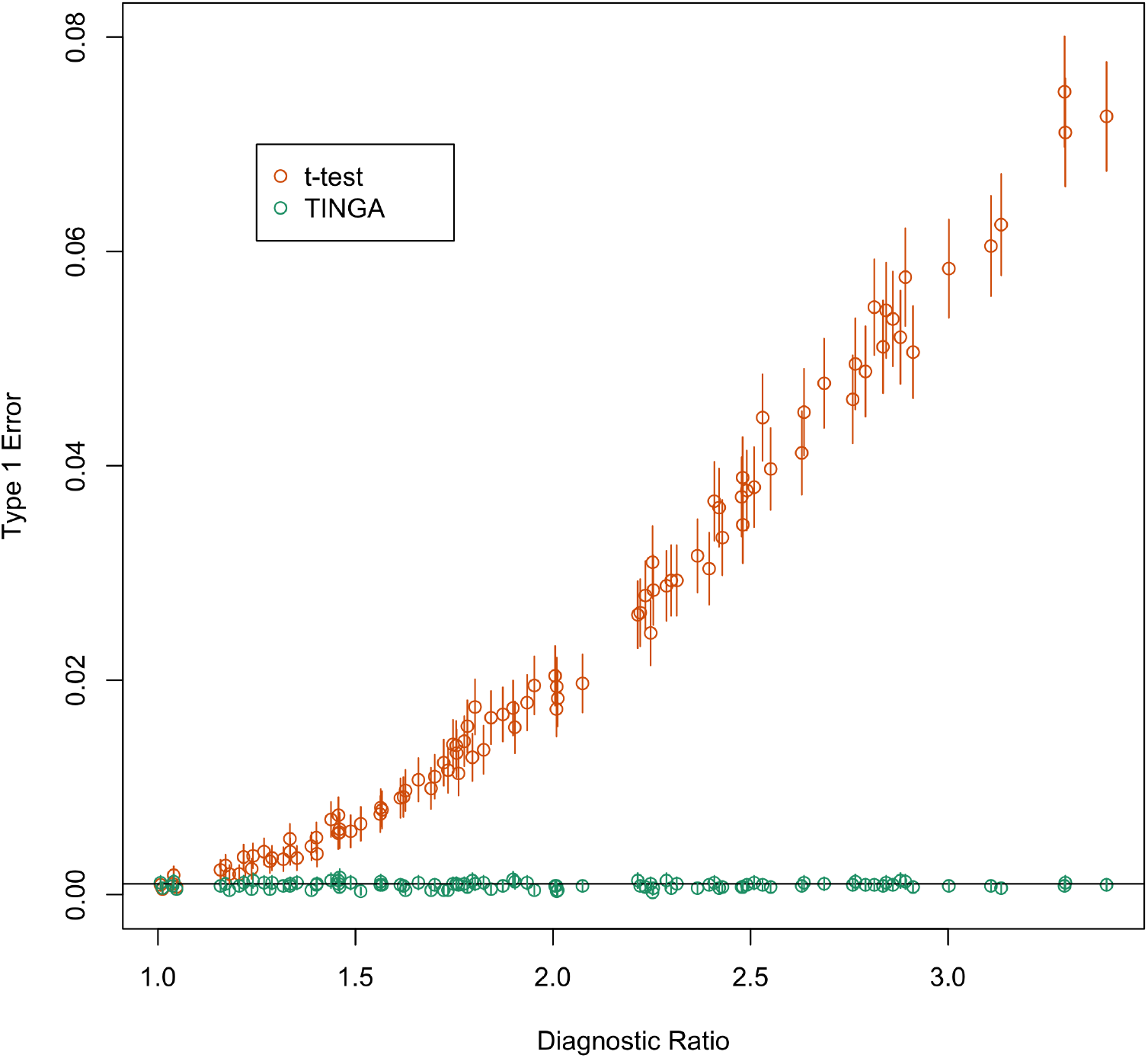
Type 1 error vs. diagnostic ratio for the standard t-test for interaction and the TINGA adjusted t-test for interaction when *Z* is associated with a causal variant. Results from 100 simulated GWISs are plotted under the null hypothesis of no interaction. Type 1 error for each GWIS is estimated based on 10^4^ unassociated SNPs. In each case, the vertical line represents the 95% confidence interval for type 1 error. The nominal type 1 error of 10^−3^, is represented by a horizontal black line.

## Analysis of flowering time in *A. thaliana*

### Data Description

We apply our methods to a data set on flowering time in *Arabidopsis thaliana* that has been previously analyzed [46]. We use the number of days between germination and flowering at 10^*°*^C as the phenotype, and we include samples from 931 selected accessions from different regions. The SNPs were filtered based on minor allele frequency (MAF) ≥ 0.03 [47]. LD pruning was done to remove variants with pairwise LD of *r*^2^ *>* 0.99 [47]. After filtering, there are 865,350 SNPs remaining. We use a LMM for the phenotype, where the GRM is computed based on all available SNPs with allele frequency ≥ 0.05.

**Table 2.**
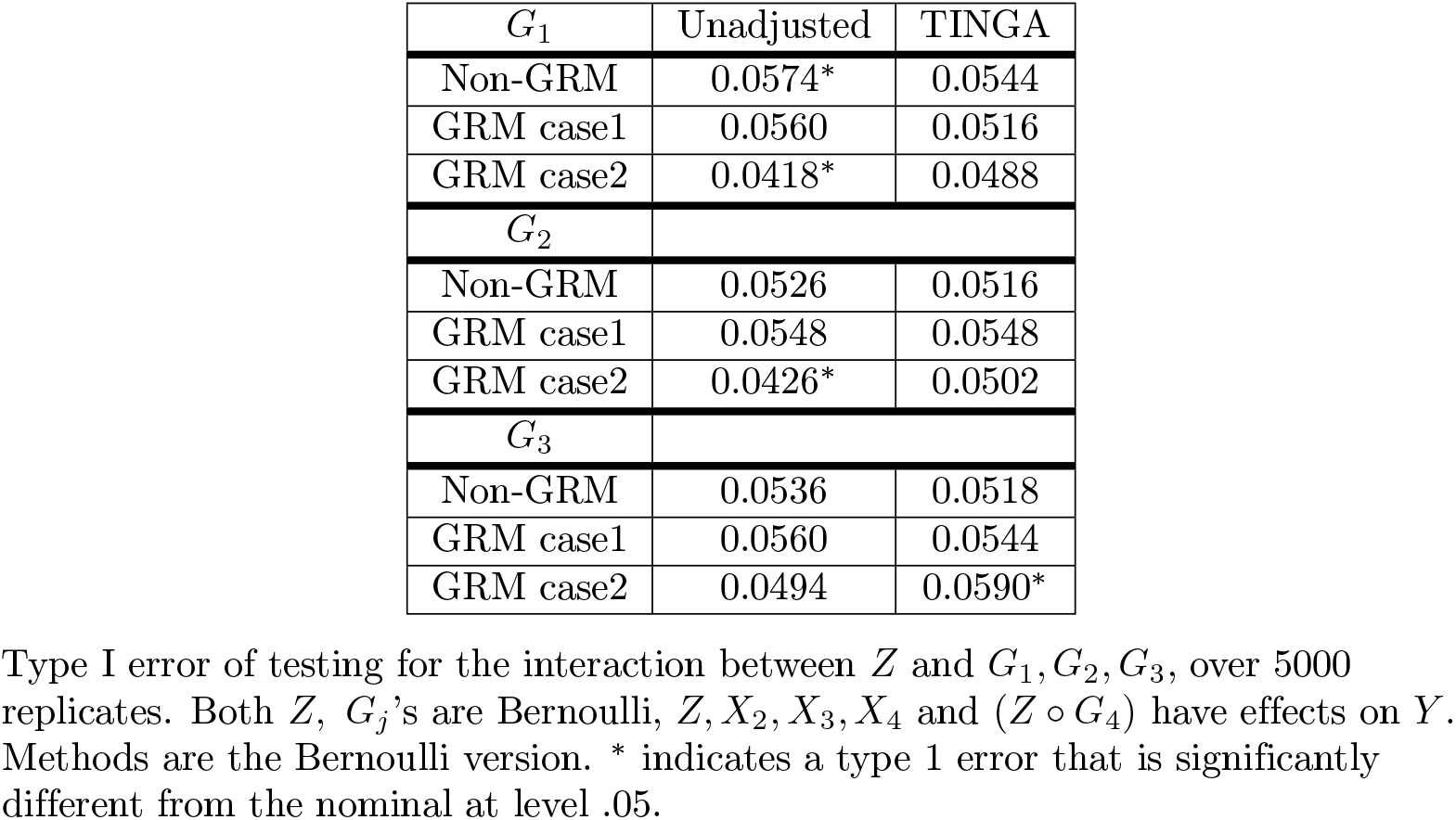
Type I error at level 0.05.

**Table 3.** Power at different p-value cutoffs.

| p-value cutoff $10^{-5}$ | Unadjusted | TINGA |
| --- | --- | --- |
| Non-GRM | 0.7046 | 0.7346 |
| GRM case1 | 0.7064 | 0.7322 |
| GRM case2 | 0.7222 | 0.8168 |
| p-value cutoff $10^{-6}$ | | |
| Non-GRM | 0.5216 | 0.5748 |
| GRM case1 | 0.5308 | 0.5810 |
| GRM case2 | 0.5566 | 0.6526 |

Power of testing for the interaction between $Z$ and $G_4$ , over 5000 replicates. Both $Z, G_j$ 's are Bernoulli, $Z, G_2, G_3, G_4$ and $(Z \circ G_4)$ have effects on $Y$ . Methods are the Bernoulli version.

### Strategy for detecting epistasis

#### Step1: Select 865 variants with smallest marginal p-values

Due to the large number of SNPs (865,350 after filtering) and the fact that we use a LMM for *Y*, it is computationally impractical to do a pairwise search over all possible pairs of SNPs for epistasis. Therefore, we start by identifying the .1% of SNPs with the smallest p-values from the ordinary GWAS based on the LMM for *Y*, which results in 865 SNPs selected. For each of these 865 SNPs, we test for interaction with with each of the 865,350 other SNPs in the genome (subject to constraints on informativeness and the constraint that the SNPs have *r*^2^ *<* .01, as described in the Methods section).

#### Step2: Perform fast, approximate, Wald tests in an LMM for testing interaction between each of the 865 selected SNPs and each of the 865,350 other SNPs in the genome

**Fig 10.**
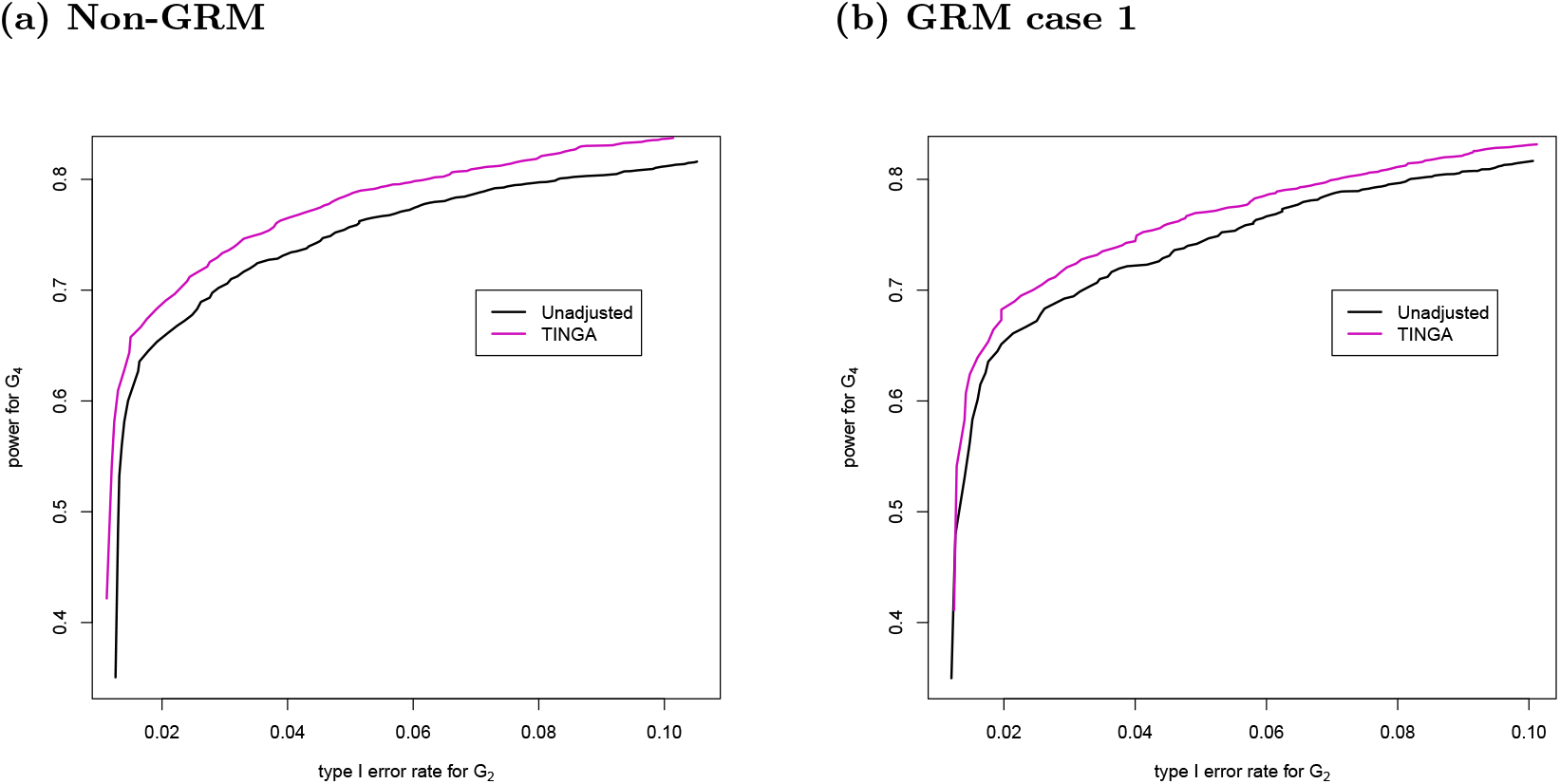
Power curves. x-axis is the type I error rates for testing *Z ° G*_2_, y-axis is the power for testing *Z ° G*_4_. Both *Z* and *G*_*j*_ are independent Bernoulli. *Z, G*_2_, *G*_3_, *G*_4_ and (*Z ° G*_4_) have effects on *Y*. (a) non-GRM case (b) GRM case 1

Even with the number of tests reduced by a factor of more than 500, we still need a fast computation strategy because we are performing interaction tests based on an LMM. We take a two-stage approach, where we first apply a fast, approximate Wald test. Then we only perform more time-consuming and accurate calculations for p-values that are small based on the fast, approximate Wald test, and we content ourselves with the coarser approximation for the p-values that are large. The key idea of the fast approximate Wald test is to regress out all variables aside from the interaction term step by step using matrix operations, so that we can avoid looping over the SNPs. We have adapted this method to LMM. (See S1 Text for details.)

#### Step3: Perform more accurate p-value calculation only for those pairs with fast approximate Wald p-value

*<* 10^−4^ Both the p-value for interaction in a LMM and the TINGA method will be applied only to those pairs with fast, approximate Wald p-value *<* 10^−4^. Furthermore, for some pairs, interaction was not tested at all because informativeness constraints were not met (we required MCC ≥ 5) or our constraint on correlation was not met (we required *r*^2^ *<* .01). After these filtering steps (based on MCC, *r*^2^ and fast approximate Wald p-value), there are 71,863 pairs of SNPs remaining, with 762 of the originally chosen SNPs having at least one pair, and these 71,863 are the pairs for which we calculate the interaction t-statistic and TINGA statistic.

#### Step4: Significance under Bonferroni correction

When applying the Bonferroni correction, we arguably only need to correct for the number of pairs that have at least one of the two SNPs in the selected set of 865 associated variants and that satisfy MCC ≥ 5 and *r*^2^ *<* .01. However, if we are being very conservative, we could consider that we are potentially searching over all distinct pairs with MCC ≥ 5 and *r*^2^ *<* .01, of which there are 2.7 *×* 10^11^. Taking in to account that two tests were performed, the Bonferroni correction level could be very conservatively taken to be 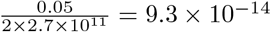.

**Fig 11.**
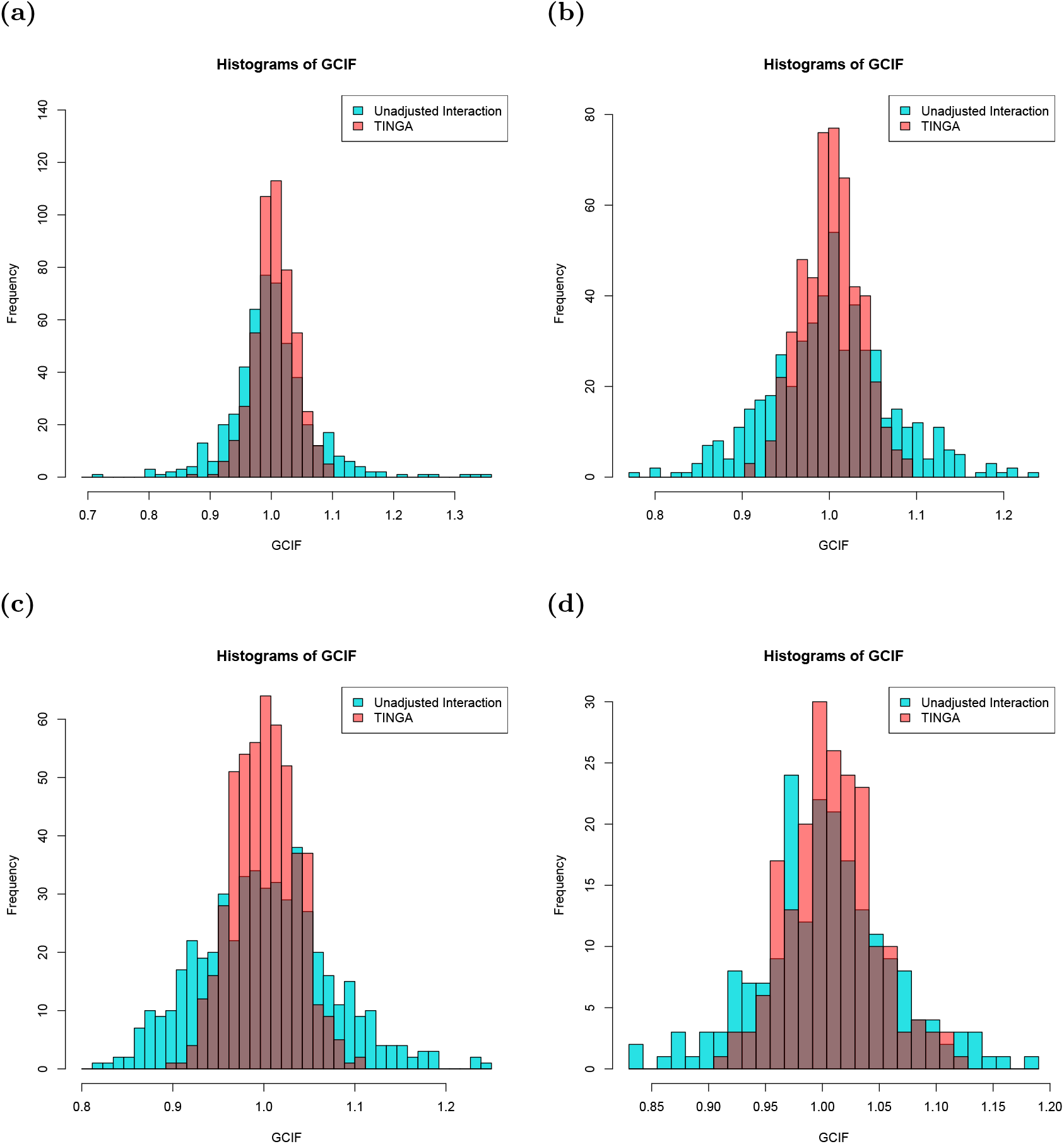
Uncorrected vs. corrected GCIF under the null. Genomic control inflation factors of interaction tests between *Z* and each of *m* = 5000 *G*_*j*_ ‘s where *Y* is the outcome; *Z* and the *G*_*j*_ ‘s are independent; (a) Both *Z* and *G*_*j*_ ‘s are Bernoulli distributed; linear model for *Y* ; 500 replicates (b) Both *Z* and *G*_*j*_ are binomial; linear model for *Y* ; 500 replicates; (c) Both *Z* and *G*_*j*_ are normal; linear model for *Y* ; 500 replicates (d) Both *Z* and *G*_*j*_ ‘s are Bernoulli; LMM for *Y* ; 200 replicates

**Fig 12.**
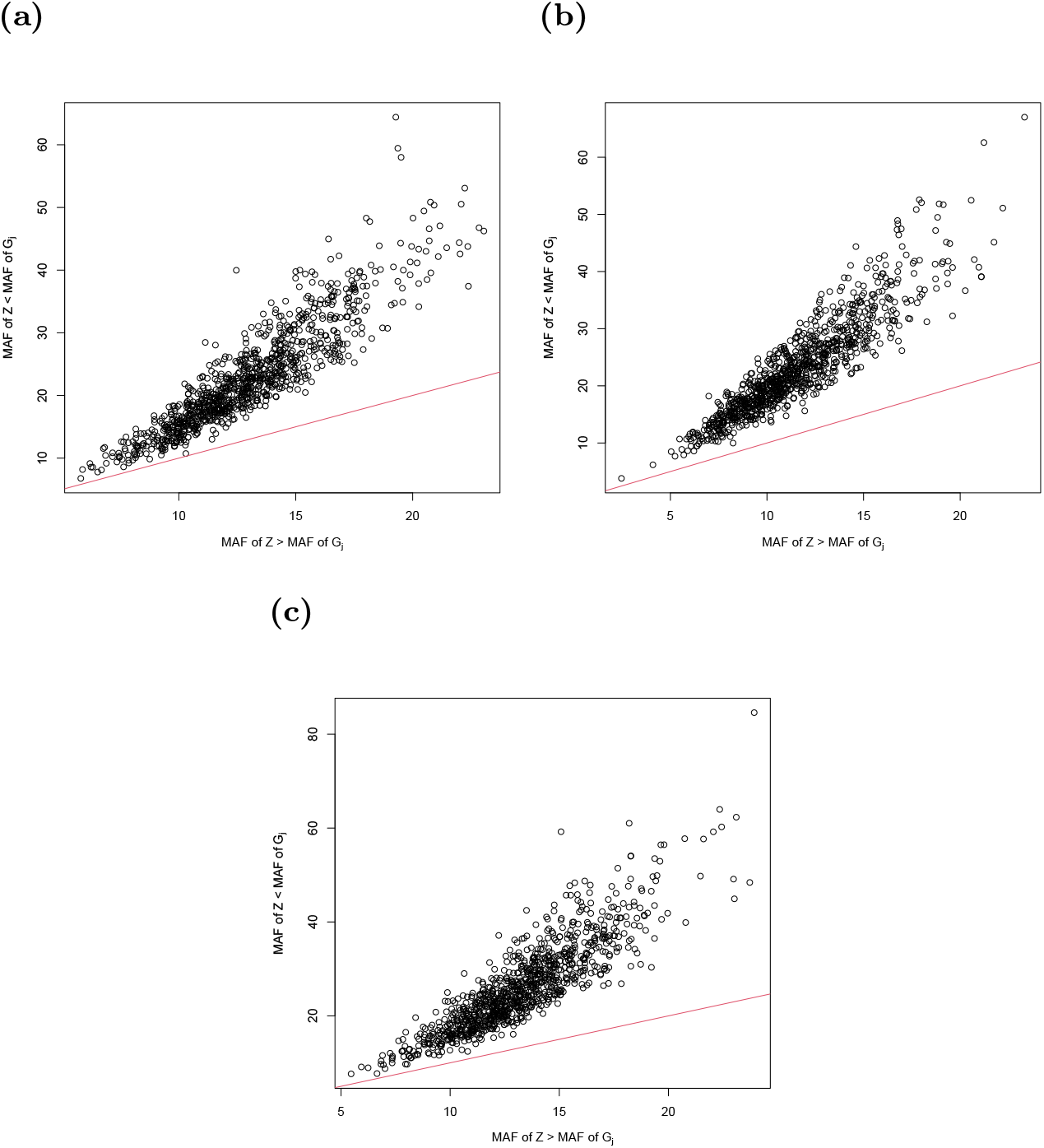
scatter plot of −*log*10 scaled p-values for the two possible TINGA analyses of interaction between a pair of genetic variants, where the *x*-axis is p-value for the case when *Z* is taken to be the variant with the larger MAF, and the *y*-axis is for the same pair where *Z* is taken to be the variant with the smaller MAF. Both *Z, G*_*j*_ are Bernoulli distributed. The two MAFs used to generate the data are 0.07 and 0.25.

## Findings

Table 4 contains information on the pairs that are significant after Bonferroni correction. Table 5 lists the corresponding genes. Among the identified SNPs, Chr5:18607017 is also detected in the study of its association with plant dry weight [13], average growth rate [13] and flowering time [46], [48]; Chr5:20430580 is also detected in the study of its association with leaf margin serrated [49]. Other SNPs are not directly found in other studies, but the genes in which they are located such as AT5G10140 [46] [50] [49] are found to be related to flowering time in many other studies.

## Example QQ-Plot for a given choice of *Z*

Of course in the data, we do not know the truth. However, it can be interesting to consider how the QQ-plot is affected by the TINGA correction for a given SNP that does not appear to show evidence of interaction. We consider SNP Chr5_18593622 (MAF 0.28) which has a relatively small p-value for SNP-trait association, but shows little evidence of interaction. For this particular SNP, in addition to performing the 2-stage process described above, we calculate both its Wald t-test and TINGA interaction p-values in a LMM for each of the 696,396 SNPs in the genome with which it has *r*^2^ *<* 0.01 and MCC ≥ 20 (skipping the step of filtering by the fast, approximate Wald test). Fig 13 displays the (differenced) QQ-plots of the p-values from these methods, with simultaneous 95% acceptance regions for i.i.d. uniform p-values outlined in red, where these use the method of [42]. (In a differenced QQ-plot, the y-axis depicts the difference between observed and expected p-values, which is particularly helpful for creating a useful visualization when the plot contains a large number of points.) We can see that for this particular SNP, the distribution of p-values is much closer to uniform after TINGA adjustment.

**Table 4.** Significant pairs.

| SNP $G_j$ | MAF | Mar p | SNP $Z$ | MAF | Mar p | MCC | Wald | TINGA |
| --- | --- | --- | --- | --- | --- | --- | --- | --- |
| Chr5:3176549 | 0.41 | 2.0e-4 | Chr1:21470240 | 0.061 | 0.070 | 26 | 3.6e-6 | 1.3e-14 |
| Chr5:3198884 | 0.44 | 2.3e-4 | Chr5:1921009 | 0.064 | 0.072 | 29 | 1.6e-7 | 5.6e-15 |
| Chr5:18607017 | 0.27 | 1.8e-5 | Chr4:4835999 | 0.052 | 0.47 | 20 | 8.7e-7 | 3.2e-16 |
| Chr5:20430580 | 0.31 | 2.4e-4 | Chr5:25047282 | 0.084 | 0.14 | 33 | 1.0e-7 | 4.6e-15 |
| Chr5:12406770 | 0.22 | 0.0076 | Chr5:25333255 | 0.083 | 1.5e-4 | 22 | 9.1e-8 | 8.7e-15 |

“Mar p”: Marginal p-value of corresponding SNP in the gene-phenotype association test.

**Table 5.** Significant pairs. The genes that the SNPs are in (black) or near (red).

| SNP $G_j$ | Gene | SNP $Z$ | Gene |
| --- | --- | --- | --- |
| Chr5:3176549 | AT5G10140 | Chr1:21470240 | AT1G58030 |
| Chr5:3198884 | AT5G10190 | Chr5:1921009 | AT5G06290 |
| Chr5:18607017 | AT5G45870 | Chr4:4835999 | AT4G08025 |
| Chr5:20430580 | AT5G50180 | Chr5:25047282 | AT5G62370 |
| Chr5:12406770 | AT5G05055 | Chr5:25333255 | AT5G63160 |

## Discussion

Identifying interaction, either *G × G* or *G × E*, can give insight into both genetic effects on a complex trait and underlying biological mechanisms, and it can also help to clarify the role of environment in the case of *G × E* testing. For testing interaction in a genomewide context, we have identified and described the “feast or famine” effect, in which different GWISs have fundamentally different null distributions. For example, if we consider GWISs in which the assumed null model holds and there is no interaction under the null hypothesis, then on average over different GWISs standard methods have correct type 1 error overall, but false positives are overly concentrated in certain GWISs (“feast” GWISs) and false negatives are overly concentrated in certain other GWISs (“famine” GWISs). If the environmental variable does interact with some predictors (either genetic variants or non-observed covariates), then the type 1 error disparity for non-interacting variants is even more extreme. In the example of Hemani et al. 2014 and 2021 [38, 45], in which the *Z* variable is associated with a causal SNP and there is model misspecification in the standard null model, the feast effect can be quite extreme, and the overall type 1 error very inflated. We show that whether a given GWIS will be a “feast” or “famine” GWIS is a reproducible property that can be predicted as a function of the observed trait and environmental values. We show that the “feast or famine” effect applies for different types of variables, including normal, binomial or binary. We show that the feast or famine effect occurs across a wide range of GxE analysis methods, including but not limited to (1) testing interaction in a linear or linear mixed model (LMM) using standard approaches such as t-tests/Wald tests, likelihood ratio tests, or score tests; (2) doing a combined interaction-association test in a linear model or LMM using standard approaches; (3) testing interaction with multiple environments or multiple SNPs, where these are modeled as random effects in a LMM using standard approaches; (4) performing tests of interaction in a GWIS where significance is assessed using permutation of the trait residuals. We show that the “feast or famine effect” affects only interaction GWAS, not ordinary association GWAS. The “feast or famine effect” can lead to excess type 1 error, reduced power, inconsistent results across studies, and failure to replicate true signal. Furthermore, we show that whether a given GWAS will be a “feast” or “famine” GWAS is a reproducible property, and that it can be corrected for.

**Fig 13.**
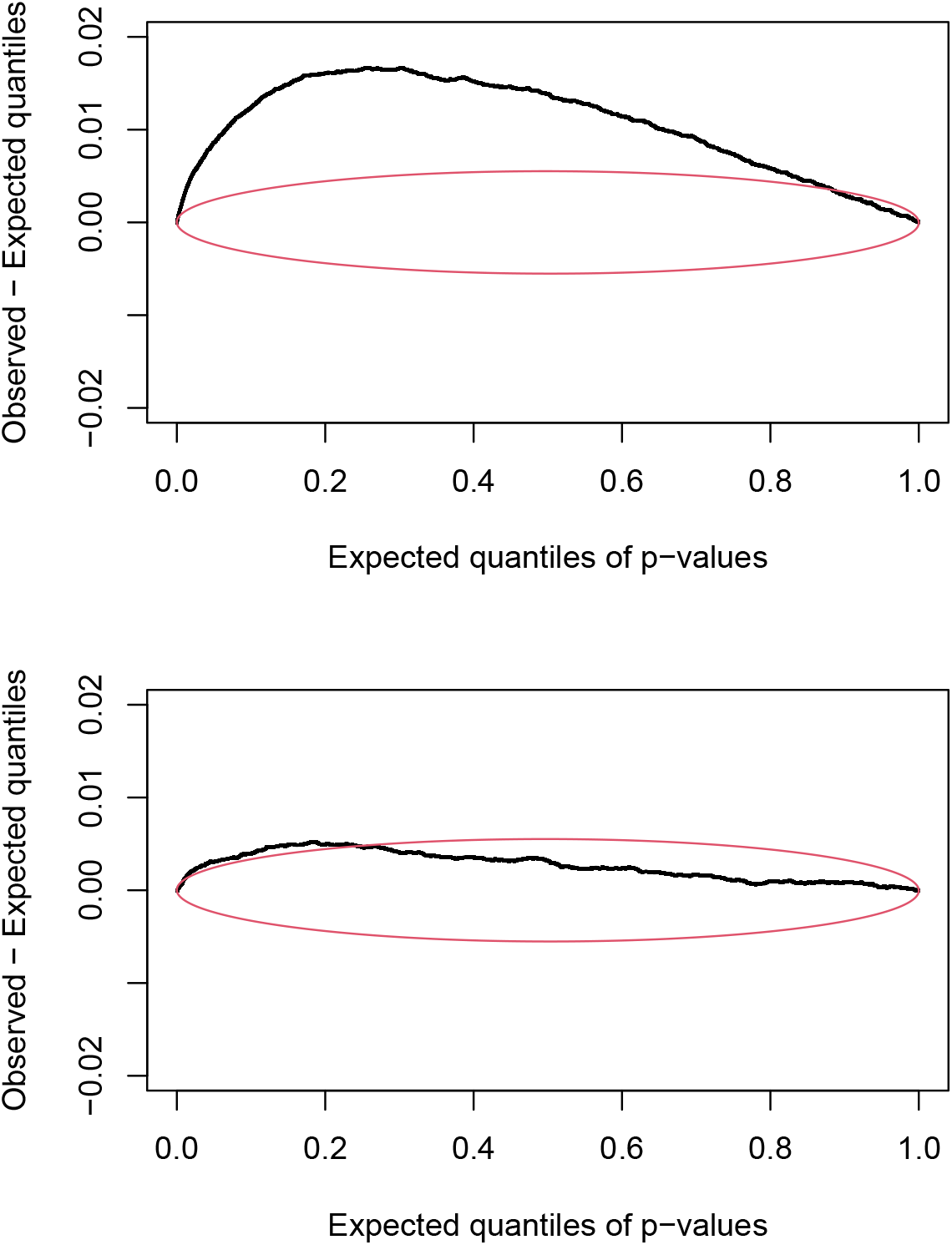
Differenced QQ-plots of p-values for interaction of SNP Chr5_18593622 with 696,396 genomewide SNPs. using (A) the t-test for interaction in an LMM and (B) TINGA. The expected quantile is plotted on the x-axis, and the difference between the observed and expected quantiles is plotted on the y-axis. The red lines give the boundaries of the 95% simultaneous acceptance region for i.i.d. uniform p-values.

We develop the TINGA method which corrects the test statistic for interaction by March 4, 2025 30/35 choosing different conditioning variables that are more appropriate for a GWAS than the standard choice. TINGA also allows for covariates and population structure through a LMM, and it accounts for heteroscedasticity. In simulations we show that TINGA can greatly reduce or eliminate the “feast or famine” effect while preserving the overall type 1 error, which we show can result in higher power.

Furthermore, we have developed a diagnostic ratio, based only on the observed (*Z, Y*), which summarizes the degree of “feast” or “famine” that would be expected for that interaction GWAS. When the diagnostic indicates only a weak effect, the researcher could more reasonably proceed with a standard analysis, while when the diagnostic indicates a strong effect, then it will be clear that use of a more sophisticated statistical method would be critical to both improve power and guard against excess false positive results. The corresponding diagnostic result could also be reported alongside any detected interaction signal, as a way of addressing concerns about statistical validity. In the context of epistasis GWAS, having a diagnostic can in addition be important for computational reasons, with sophisticated statistical methods such as TINGA reserved for choices of variants for which a strong “feast or famine” effect is predicted.

We apply TINGA to a GWAS for flowering time in *A. thaliana*. Using TINGA we detect 5 significant interactions after Bonferroni correction, where all the detected interactions involve loci identified in previous studies as associated with flowering time. This demonstrates the potential of the TINGA method for detecting interaction in a GWAS.

For epistasis detection in a GWAS, there is a computational challenge in testing epistasis for all possible pairs of variants. When the model for *Y* is a LMM, as in our data analysis, this computational challenge is made much greater, even for the usual LMM-based t-test for interaction without any correction. We have developed a fast approximate version of the LMM-based t-test for interaction, and we use it as part of an adaptive approach to genomewide testing, where more accurate but time-consuming methods are applied only if the approximate p-value is sufficiently small. In other words, our strategy is to spend more computational time on small p-values and to be content with coarse approximations to large p-values. In future work, there could be further scope for making faster algorithms for all aspects of interaction testing with a LMM in a GWAS context.

## Acknowledgments

This study was supported by NIH grant R01 HG001645 (to M.S.M.). We thank Peter Laurin for suggesting the data set and for help with preliminary data analysis.

