## Supplemental Text, Tables and Figures for "Overcoming the “feast or famine” effect: improved interaction testing in genome-wide association studies"

### S1 Text

#### R script to calculate p-values for the two-sided equal local levels test for i.i.d. uniformity

A two-sided equal local levels (ELL) test for i.i.d. uniformity a set of variables $X_1, \ldots, X_n$ is described in [1], who created the R package qqconf, which is available on CRAN. The primary purpose of qqconf is to generate appropriate simultaneous testing bands for a QQ-plot, but in addition, the functions available in qqconf can be used to generate p-values for the two-sided ELL test for i.i.d. uniformity.

In the R code below, suppose $x \in (0,1)^n$. The code obtains a p-value for the deviation of $x$ from i.i.d. uniform(0,1). The test is a QQ-plot based ELL test. It answers the question: what is the largest level $\alpha$ for the acceptance region for the QQ-plot that would result in non-rejection of $x$, where the acceptance region is based on 2-sided ELL.

```
library(qqconf)
qqpvu <- function(x){
n = length(x)
tmp1 = sort(x)
tmp2 = pbeta(tmp1,c(1:n),c(n:1))
tmp3 = min(min(tmp2),1-max(tmp2))*2
lb = qbeta(tmp3/2,c(1:n),c(n:1))
ub = qbeta(1-tmp3/2,c(1:n),c(n:1))
get_level_from_bounds_two_sided(lb,ub)
}
```

#### Supplemental Fig. 1

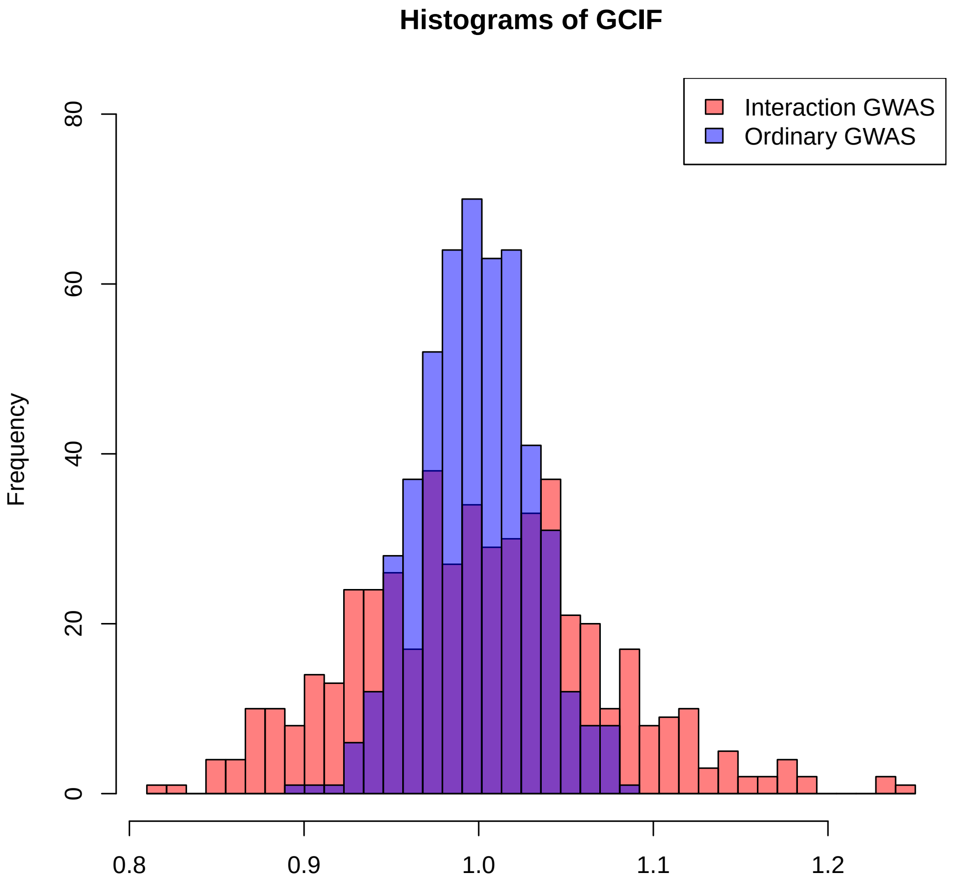


**Fig S1. Histograms of GCIFs for interaction GWAS and for ordinary, non-interaction GWAS: Case of normal $Z$ and $G_j$** Each histogram is based on $r = 5{,}000$ simulated null GWASs in which $Y$, $Z$ and $G$ are simulated as vectors of i.i.d. normal random variables. For each GWAS, two different GCIFs are calculated, each based on $m = 5{,}000$ test statistics. The GCIF for ordinary (non-interaction) GWAS uses the $m$ genetic association tests between $Y$ and the $G_j$s, and the GCIF for interaction GWAS uses the $m$ interaction tests based on Model 1 in the main text. The blue histogram represents the $r$ resulting GCIFs for ordinary (non-interaction) association testing, and the red histogram represents the $r$ resulting GCIFs for interaction testing.

#### Simulation settings used in the paper

##### Setting 1: Null, non-GRM case

- $n = 1000,\ m = 5000$
- $Z \sim Bernoulli(m_Z),\ m_Z \sim Unif(0.1, 0.9)$

- $G_j \overset{\text{indep.}}{\sim} Bernoulli(m_j),\ m_j \overset{\text{iid}}{\sim} Unif(0.1, 0.9), cor(Z, G_j) = 0,\ j = 1, 2, ..., m$
- $Y = \alpha + \epsilon,\ \alpha \sim Unif(-10, 10),\ \epsilon \sim N(0, 1)$

When simulate each $G_j$, we check the following 2 conditions and keep re-generating $G_j$ until both of the 2 conditions are satisfied:

1. $|cor(G_j, Z)| \leq 0.1$
2. $MCC(G_j, Z) \geq 5$

**Setting 2: Null, GRM case**

- $n = 1000,\ m = 5000$
- $Z \sim Bernoulli(m_Z),\ m_Z \sim Unif(0.1, 0.9)$
- $G_j \overset{\text{indep.}}{\sim} Bernoulli(m_j),\ m_j \overset{\text{iid}}{\sim} Unif(0.1, 0.9), cor(Z, G_j) = 0,\ j = 1, 2, ..., m$
- GRM $K$ is calculated from a simulated genotype matrix $G$, which is independent of $Z$ and $G_j$'s:

$$G = (g_1, ..., g_{10000}),\ g_i \overset{indep.}{\sim} Ber(f_i),\ f_i \overset{iid}{\sim} Unif(0.1, 0.9)$$
$$\tilde{G} = (\tilde{g_1}, ...\tilde{g}_{10000}),\ \tilde{g_i} = \frac{g_i - \bar{g_i}}{\sqrt{\bar{g_i}(1 - \bar{g_i})}}$$
$$K = \frac{1}{10000} \tilde{G}\tilde{G}^T$$

- $\alpha \sim Unif(-10, 10),\ h^2 = 0.3,\ \sigma_T^2 = 1$
- $Y = \alpha + \epsilon,\ \epsilon \sim N(0,\ \sigma_T^2 \left(h^2 K + (1 - h^2)I\right)$

When simulate each $G_j$, we check the following 2 conditions and keep re-generating $G_j$ until both of the 2 conditions are satisfied:

1. $|cor(G_j, Z)| \leq 0.1$
2. $MCC(G_j, Z) \geq 5$

**Setting 3: Alternative, non-GRM case**
Simulation setting:

- $n = 1000,\ m = 4$
- $Z \sim Ber(m_Z),\ m_Z \sim Unif(0.1, 0.9)$
- $G_j \overset{\text{indep.}}{\sim} Ber(m_j),\ m_j \overset{\text{iid}}{\sim} Unif(0.1, 0.9),\ cor(G_j, Z) = 0,\ j = 1, 2, 3, 4$
- $Y = \alpha + \gamma Z + \sum_{j=1}^m \beta_j G_j + \sum_{i=j}^m \delta_j (Z - m_Z)(G_j - m_j) + \epsilon$
- $(\beta_1, \beta_2, \beta_3, \beta_4) = \frac{1}{2} \times \left(0, \sqrt{\frac{0.025}{\sigma_{G2}^2}}, -\sqrt{\frac{0.025}{\sigma_{G3}^2}}, \sqrt{\frac{0.05}{\sigma_{G4}^2}}\right)$
- $\gamma = \sqrt{\frac{0.025}{\sigma_z^2}}$
- $(\delta_1, \delta_2, \delta_3, \delta_4) = (0, 0, 0, \sqrt{\frac{0.025}{\sigma_{(G_4 \circ Z)}^2}})$

- $\epsilon \sim N(0,1)$

When simulate each $G_j$, we check the following 2 conditions and keep re-generating $G_j$ until both of the 2 conditions are satisfied:

1. $|cor(G_j, Z)| \leq 0.1$
2. $MCC(G_j, Z) \geq 5$

**Setting 4: Alternative, GRM case**

All the simulation settings are the same as those for the non-GRM case except that there is a GRM in the covariance matrix of $Y$:

GRM $K$ is calculated from a simulated genotype matrix $G$, which is independent of $Z$ and $G_j$'S:

- First generate $M = 10,000$ independent SNPs $g_i$:
- $g_i \overset{indep.}{\sim} Ber(f_i),\ f_i \overset{iid}{\sim} Unif(0.1, 0.9)$
- Only keep those $g_i$'s with MAF $> 0.05$
- Let $\tilde{g_i} = \frac{g_i - \bar{g_i}}{\sqrt{\bar{g_i}(1-\bar{g_i})}}$
- $K = \frac{1}{M}\sum_i g_i g_i^T$

When generation $Y$, we have

$$\epsilon \sim N(0,\ \sigma_T^2 \left(h^2 K + (1-h^2)I\right)$$

where $\sigma_T^2 = 1,\ h^2 = 0.3$

For the experiments where we need a large number of replicates, it is not necessary to have as many different GRMs as the number of replicates. For 5000 replicates, we only simulate 10 different GRMs: for every 500 replicates, we simulate a new GRM.

**Setting 5: Different frequency levels for $Z$ and $G_j$**

- $n = 3000$
- $G_j \sim Ber(0.07),\ Z \sim Ber(0.25),\ G_j, Z$ independent
- $T = \alpha + bG_j + rZ + \delta(G_j - \mu_j)(Z - \mu_Z) + \epsilon,\ \epsilon \sim\ N(0,\ 1)$
- $\alpha = 1,\ \delta = \sqrt{\frac{0.025}{\sigma_x^2 \sigma_z^2}} = 1.43$
- $(Z, G_j)$ are filtered by criterion of minimum cell count $\geq 20$

**Setting 6: Null, non-GRM case where $Z$, $G_j$ both normal**

- $n = 1000,\ m = 5000$
- $Z \sim Normal(m_Z, 1),\ m_Z \sim Unif(-10, 10)$
- $G_j \overset{\text{indep.}}{\sim} Normal(m_j, 1),\ m_j \overset{\text{iid}}{\sim} Unif(-10, 10), cor(Z, G_j) = 0,\ j = 1, 2, ..., m$
- $Y = \alpha + \epsilon,\ \alpha \sim Unif(-10, 10),\ \epsilon \sim N(0, 1)$

**Setting 7: Null, non-GRM case where $Z$, $G_j$ both Binomial**

- $n = 1000,\ m = 5000$
- $Z \sim Binom(2, m_Z),\ m_Z \sim Unif(0.1, 0.9)$
- $G_j \overset{\text{indep.}}{\sim} Binom(2, m_j),\ m_j \overset{\text{iid}}{\sim} Unif(0.1, 0.9), cor(Z, G_j) = 0,\ j = 1, 2, ..., m$
- $Y = \alpha + \epsilon,\ \alpha \sim Unif(-10, 10),\ \epsilon \sim N(0, 1)$

**Setting 8: Alternative, non-GRM case, $Z$ normal, $G_j$ Binomial**

- $n = 1000,\ m = 4$
- $Z \sim N(m_Z, 1),\ m_Z \sim Unif(-10, 10)$
- $G_j \overset{\text{indep.}}{\sim} Binom(2, m_j),\ m_j \overset{\text{iid}}{\sim} Unif(0.1, 0.9),\ cor(G_j, Z) = 0,\ j = 1, 2, 3, 4$
- $Y = \alpha + \gamma Z + \sum_{j=1}^{m} \beta_j G_j + \sum_{i=j}^{m} \delta_j (Z - m_Z)(G_j - m_j) + \epsilon$
- $(\beta_1, \beta_2, \beta_3, \beta_4) = \frac{1}{2} \times \left(0, \sqrt{\frac{0.025}{\sigma_{x_2^2}}}, -\sqrt{\frac{0.025}{\sigma_{x_3^2}}}, \sqrt{\frac{0.05}{\sigma_{x_4^2}}}\right)$
- $\gamma = \sqrt{\frac{0.025}{\sigma_z^2}}$
- $(\delta_1, \delta_2, \delta_3, \delta_4) = (0, 0, 0, \sqrt{\frac{0.025}{\sigma^2_{(x_4 \circ z)}}})$
- $\epsilon \sim N(0, 1)$

**Setting 9: Alternative, non-GRM case, $Z$ normal, $G_j$ Bernoulli**

- $n = 1000,\ m = 4$
- $Z \sim N(m_Z, 1),\ m_Z \sim Unif(-10, 10)$
- $G_j \overset{\text{indep.}}{\sim} Ber(m_j),\ m_j \overset{\text{iid}}{\sim} Unif(0.1, 0.9),\ cor(G_j, Z) = 0,\ j = 1, 2, 3, 4$
- $Y = \alpha + \gamma Z + \sum_{j=1}^{m} \beta_j G_j + \sum_{i=j}^{m} \delta_j (Z - m_Z)(G_j - m_j) + \epsilon$
- $(\beta_1, \beta_2, \beta_3, \beta_4) = \frac{1}{2} \times \left(0, \sqrt{\frac{0.025}{\sigma_{x_2^2}}}, -\sqrt{\frac{0.025}{\sigma_{x_3^2}}}, \sqrt{\frac{0.05}{\sigma_{x_4^2}}}\right)$
- $\gamma = \sqrt{\frac{0.025}{\sigma_z^2}}$
- $(\delta_1, \delta_2, \delta_3, \delta_4) = (0, 0, 0, \sqrt{\frac{0.025}{\sigma^2_{(x_4 \circ z)}}})$
- $\epsilon \sim N(0, 1)$

**Setting 10: Alternative, non-GRM case, $Z$ Binomial, $G_j$ Binomial**

- $n = 1000,\ m = 4$
- $Z \sim Binom(2, m_Z),\ m_Z \sim Unif(0.1, 0.9)$
- $G_j \overset{\text{indep.}}{\sim} Binom(2, m_j),\ m_j \overset{\text{iid}}{\sim} Unif(0.1, 0.9),\ cor(G_j, Z) = 0,\ j = 1, 2, 3, 4$
- $Y = \alpha + \gamma Z + \sum_{j=1}^{m} \beta_j G_j + \sum_{i=j}^{m} \delta_j (Z - m_Z)(G_j - m_j) + \epsilon$
- $(\beta_1, \beta_2, \beta_3, \beta_4) = \frac{1}{2} \times \left(0, \sqrt{\frac{0.025}{\sigma_{x_2^2}}}, -\sqrt{\frac{0.025}{\sigma_{x_3^2}}}, \sqrt{\frac{0.05}{\sigma_{x_4^2}}}\right)$
- $\gamma = \sqrt{\frac{0.025}{\sigma_z^2}}$
- $(\delta_1, \delta_2, \delta_3, \delta_4) = (0, 0, 0, \sqrt{\frac{0.025}{\sigma^2_{(x_4 \circ z)}}})$
- $\epsilon \sim N(0, 1)$

**Setting 11: Alternative, 3 sub-populations, $Z$ Bernoulli, $G_j$ Bernoulli**

For this setting we simulate a population structure with 3 sub-populations using a Balding-Nichols model.

We assign 1/3 of the total population to each sub-population. In our case, $n = 1000$, so the sample sizes for sub-population 1, 2, 3 are $333, 333, 334$ respectively. Let the fixation index $F = 0.01$, which is representative of the population structure seen in humans within continents.

For each SNP $s$, the ancestral allele frequency $p_s$ is drawn $\overset{\text{iid}}{\sim} Unif(0.2, 0.8)$ (iid across SNPs). For each sub-population $k = 1, 2, 3$, the allele frequency $p_k$ is drawn independently from $Beta(\frac{p_s(1-F)}{F}, \frac{(1-p_s)(1-F)}{F})$ . Then for an individual assigned to sub-population $k$, the genotype is drawn iid from $Ber(p_k)$. We only keep the SNPs with MAF $\geq 0.05$.

We use the above strategy to simulate one $Z$ and $m = 4$ $G_j$'s, independently. When simulate each $G_j$, we check the following 2 conditions and keep re-generating $G_j$ until both of the 2 conditions are satisfied:

1. $|cor(G_j, Z)| \leq 0.1$
2. $MCC(G_j, Z) \geq 5$

For the GRM, we simulate $10^5$ independent SNPs, these SNPs are also independent from $Z$ and $G_j$'s.

We simulate $Y$ by the following model:

$$Y = \alpha + \gamma Z + \sum_{j=1}^{m} \beta_j G_j + \sum_{i=j}^{m} \delta_j (Z - m_Z)(G_j - m_j) + \zeta_1 \mathbb{I}_1 + \zeta_2 \mathbb{I}_2 + \epsilon$$

where

- $(\beta_1, \beta_2, \beta_3, \beta_4) = 0.2 \times \left(0, \sqrt{\frac{0.025}{\sigma_{x_2^2}}}, -\sqrt{\frac{0.025}{\sigma_{x_3^2}}}, \sqrt{\frac{0.05}{\sigma_{x_4^2}}}\right)$
- $\gamma = \sqrt{\frac{0.025}{\sigma_z^2}}$
- $(\delta_1, \delta_2, \delta_3, \delta_4) = (0, 0, 0, \sqrt{\frac{0.02}{\sigma^2_{(x_4 \circ z)}}})$

- $\zeta_1 = \zeta_2 = 3\sqrt{0.15}$, $\mathbb{I}_1, \mathbb{I}_2$ are indicators of membership in sub-population 1, 2. In this way, the variance explained by $\zeta_1\mathbb{I}_1 + \zeta_2\mathbb{I}_2$ will be 0.3, which is the same as the heritability $h^2$ in previous settings of GRM such as setting 2.
- $\epsilon \sim N(0, 0.7I_n)$, which makes the total variance equal 1, the same as previous settings.

Note although we simulate $Y$ with those indicators of sub-populations, when we do the analysis, we do NOT use the indicators as covariates. We use the same LMM approach as before.

#### Computation of $T,\ E(T|y,z),\ Var(T|y,z)$

When testing for interaction between $z$ and each of the $x$'s, we test $H_0 : \delta = 0\ vs.\ H_1 : \delta \neq 0$ in the model

$$y \sim N(\alpha + \beta_1 x + \gamma z + \delta(x \circ z),\ \sigma_T^2 \Sigma)$$

Where $\Sigma = \frac{\sigma_g^2}{\sigma_T^2} K + \frac{\sigma_e^2}{\sigma_T^2} I$ is assumed known, $(x \circ z)_j = (x_j - \bar{x})(z_j - \bar{z})$.

The Wald statistic is

$$T = \frac{\sqrt{n-4}\,(x \circ z)^T P_u y}{\sqrt{(x \circ z)^T P_u (x \circ z)\ \cdot y^T P_u y - ((x \circ z)^T P_u y)^2}}$$

Where $P_u = \hat{\Sigma}^{-1} - \hat{\Sigma}^{-1} U \left(U^T \hat{\Sigma}^{-1} U\right)^{-1} U^T \hat{\Sigma}^{-1},\ U = (1,\ x,\ z)$. We are mainly interested in the numerator part $(x \circ z)^T P_u y$.

Note that $P_u$ is the matrix that regresses out $U$ in above linear mixed model. We could compute it explicitly using an iterative method:

In simple linear model $y \sim N(U\beta,\ \sigma^2 I)$ where $U$ contains intercept term 1, variable $x$ and variable $z$. Then $H = P_0 = I - \frac{1}{n}\mathbf{1}\mathbf{1}^T$ will be the projection matrix that project out $\mathbf{1}, P_1 = P_0 - \frac{P_0 x x^T P_0}{x^T P_0 x}$ will be the matrix that further regresses out $x$ (therefore regressing out $(1,\ x)), P_2 = P_1 - \frac{P_1 z z^T P_1}{z^T P_1 z}$ will be the matrix that regresses out $(1,\ \ x,\ \ z)$.

To apply above formula to the LMM case, we let $C^T C = \hat{\Sigma}^{-1},\ CU = \widetilde{U} = \left(\widetilde{1},\ \widetilde{x},\ \widetilde{z}\right)$. Then $Cy = \widetilde{y} \sim \left(\widetilde{U}\beta + \gamma\widetilde{(x \circ z)},\ \sigma_T^2 I\right)$, here we denote $\widetilde{w}$ denotes $Cw$ for any vector $w$.

$$P_u = \hat{\Sigma}^{-1} - \hat{\Sigma}^{-1} U \left(U^T \hat{\Sigma}^{-1} U\right)^{-1} U^T \hat{\Sigma}^{-1} = C^T C - C^T \widetilde{U} \left(\widetilde{U}^T \widetilde{U}\right)^{-1} \widetilde{U} C$$

$$(x \circ z)^T P_u y = \widetilde{(x \circ z)}^T \left(I - \widetilde{U}\left(\widetilde{U}^T \widetilde{U}\right)^{-1} \widetilde{U}\right) \widetilde{y} = \widetilde{(x \circ z)}^T P_{\widetilde{u}} \widetilde{y}$$

Where $P_{\widetilde{u}} = I - \widetilde{U}\left(\widetilde{U}^T \widetilde{U}\right)^{-1} \widetilde{U}$.

Note $P_{\widetilde{u}}$ is the matrix that regresses out $\widetilde{U}$ in the simple linear model, so we can compute it using above iterative method:

$$\widetilde{H} = P_0\prime = I - \widetilde{1}\left(\widetilde{1}^T \widetilde{1}\right)^{-1} \widetilde{1}^T$$

$$P_1' = P_0' - \frac{P_0\prime \widetilde{x}\widetilde{x}^T P_0\prime}{\widetilde{x}^T P_0' \widetilde{x}}$$

$$P_{\widetilde{u}} = P_1' - \frac{P_1'\widetilde{z}\widetilde{z}^T P_1\prime}{\widetilde{z}^T P_1\prime\widetilde{z}}$$

Therefore, we can iteratively compute an explicit expression for $P_{\widetilde{u}}$:

$$P_1' = \widetilde{H} - \frac{\left(\widetilde{H}\widetilde{x}\right)\left(\widetilde{H}\widetilde{x}\right)^T}{\widetilde{x}^T\widetilde{H}\widetilde{x}}$$

$$P_{\widetilde{u}} = \widetilde{H} - \frac{\left(\widetilde{H}\widetilde{x}\right)\left(\widetilde{H}\widetilde{x}\right)^T}{\widetilde{x}^T\widetilde{H}\widetilde{x}} - \frac{(\widetilde{H}\widetilde{z} - \frac{\left(\widetilde{H}\widetilde{x}\right)\left(\widetilde{H}\widetilde{x}\right)^T\left(\widetilde{H}\widetilde{z}\right)}{\widetilde{x}^T\widetilde{H}\widetilde{x}})(\widetilde{z}^T\widetilde{H} - \frac{(\widetilde{z}^T\widetilde{H})\left(\widetilde{H}\widetilde{x}\right)\left(\widetilde{H}\widetilde{x}\right)^T}{\widetilde{x}^T\widetilde{H}\widetilde{x}})}{\widetilde{z}^T\widetilde{H}\widetilde{z} - \frac{(\left(\widetilde{H}\widetilde{z}\right)^T\left(\widetilde{H}\widetilde{x}\right))^2}{\widetilde{x}^T\widetilde{H}\widetilde{x}}}$$

Let $S_{ab} = \left(\widetilde{H}a\right)^T\left(\widetilde{H}b\right)$ for any vectors $a,\ b$, then

$$\widetilde{(x\circ z)}^T P_{\widetilde{u}}\widetilde{y}$$
$$= S_{\widetilde{(x\circ z)}\widetilde{y}} - \frac{S_{\widetilde{(x\circ z)}\widetilde{x}}\, S_{\widetilde{z}\widetilde{y}}\, S_{\widetilde{z}\widetilde{z}} + S_{\widetilde{(x\circ z)}\widetilde{z}}\, S_{\widetilde{z}\widetilde{y}}\, S_{\widetilde{x}\widetilde{x}} - S_{\widetilde{(x\circ z)}\widetilde{z}}\, S_{\widetilde{x}\widetilde{z}}\, S_{\widetilde{x}\widetilde{y}} - S_{\widetilde{(x\circ z)}\widetilde{x}}\, S_{\widetilde{x}\widetilde{z}}\, S_{\widetilde{y}\widetilde{z}}}{S_{\widetilde{x}\widetilde{x}}\, S_{\widetilde{z}\widetilde{z}} - S^2_{\widetilde{x}\widetilde{z}}}$$

Note this formula is the same as the formula we get previously for non-GRM case, except that $S_{ab}$ is the simple inner product in that case. We can then transform these variables back and get an expression in terms of the original variables:

$$S_{\widetilde{x}\widetilde{y}} = \widetilde{x}^T\widetilde{H}\widetilde{y} = x^T C^T\left(I - \widetilde{1}\left(\widetilde{1}^T\widetilde{1}\right)^{-1}\widetilde{1}^T\right)Cy$$
$$= x^T\left(\hat{\Sigma}^{-1} - \hat{\Sigma}^{-1}1\left(1^T\hat{\Sigma}^{-1}1\right)^{-1}1\hat{\Sigma}^{-1}\right)y$$
$$= x^T\hat{H}y$$

Where $\hat{H} = \hat{\Sigma}^{-1} - \hat{\Sigma}^{-1}1\left(1^T\hat{\Sigma}^{-1}1\right)^{-1}1\hat{\Sigma}^{-1}$. Similarly for other $S_{ab}$ terms. Then we reorganize these terms and get

$$\widetilde{(x\circ z)}^T P_{\widetilde{u}}\widetilde{y} = (x\circ z)^T\hat{H}\left(y - \frac{S_{\widetilde{z}\widetilde{y}}\, S_{\widetilde{x}\widetilde{x}} - S_{\widetilde{x}\widetilde{z}}\, S_{\widetilde{x}\widetilde{y}}}{S_{\widetilde{x}\widetilde{x}}\, S_{\widetilde{z}\widetilde{z}} - S^2_{\widetilde{x}\widetilde{z}}}z\right) - \frac{S_{\widetilde{x}\widetilde{y}}\, S_{\widetilde{z}\widetilde{z}} - S_{\widetilde{x}\widetilde{z}}\, S_{\widetilde{y}\widetilde{z}}}{S_{\widetilde{x}\widetilde{x}}\, S_{\widetilde{z}\widetilde{z}} - S^2_{\widetilde{x}\widetilde{z}}}\ (x\circ z)^T\hat{H}x$$

We want the conditional mean and variance of it using the conditional distribution of $x$ on $(y,\ z)$.

We simplify terms $\frac{S_{\widetilde{z}\widetilde{y}}\, S_{\widetilde{x}\widetilde{x}} - S_{\widetilde{x}\widetilde{z}}\, S_{\widetilde{x}\widetilde{y}}}{S_{\widetilde{x}\widetilde{x}}\, S_{\widetilde{z}\widetilde{z}} - S^2_{\widetilde{x}\widetilde{z}}}$ and $\frac{S_{\widetilde{x}\widetilde{y}}\, S_{\widetilde{z}\widetilde{z}} - S_{\widetilde{x}\widetilde{z}}\, S_{\widetilde{y}\widetilde{z}}}{S_{\widetilde{x}\widetilde{x}}\, S_{\widetilde{z}\widetilde{z}} - S^2_{\widetilde{x}\widetilde{z}}}$ by dividing both numerator and denominator by $n^2$ and approximating $\frac{S_{ab}}{n}$ terms by their asymptotic means under the null conditional distribution given $(y, z)$. .

Assume $x|y,\ z \overset{\text{approx.}}{\sim} N\left(\mu_2,\ V_2\right)$, all following calculations are conditioning on $(y,\ z)$:

$$\frac{S_{\widetilde{x}\widetilde{x}}}{n} = \frac{x^T\hat{H}x}{n} \sim \frac{1}{n}\left(\mu_2\hat{H}\mu_2\right) + \frac{1}{n}tr\left(\hat{H}V_2\right)$$
$$\frac{S_{\widetilde{x}\widetilde{z}}}{n} = \frac{x^T\hat{H}z}{n} \sim \frac{1}{n}\left(\mu_2\hat{H}z\right)$$
$$\frac{S_{\widetilde{x}\widetilde{y}}}{n} = \frac{x^T\hat{H}y}{n} \sim \frac{1}{n}\left(\mu_2\hat{H}y\right)$$

Denote the simplified terms of $\frac{S_{\widetilde{zy}}S_{\widetilde{xx}}-S_{\widetilde{xz}}S_{\widetilde{xy}}}{S_{\widetilde{xx}}S_{\widetilde{zz}}-S_{\widetilde{xz}}^2}$ and $\frac{S_{\widetilde{xy}}S_{\widetilde{zz}}-S_{\widetilde{xz}}S_{\widetilde{yz}}}{S_{\widetilde{xx}}S_{\widetilde{zz}}-S_{\widetilde{xz}}^2}$ by $\alpha_1,\ \alpha_2$, respectively.

Note $x \circ z = D_{Hz}Hx$, where $D_{Hz}$ is the diagonal matrix whose diagonal entries are $Hz$. Then we get the approximated numerator of Wald statistic:

$$T \approx x^T H D_{Hz} \hat{H}\left(y - \alpha_1 z\right) - \alpha_2 x^T H D_{Hz}\hat{H}x = x^T Bx + b^T x$$

Where $B =\ -\alpha_2 H D_{Hz}\hat{H},\ \ b = HD_{Hz}\hat{H}(y - \alpha_1 z)$ are constants conditioning on $(y,\ z)$.

Note that $B$ is not necessarily symmetric (except for the non-GRM case, where $\Sigma = I,\ \hat{H} = \hat{\sigma}_T^{-2}H$). We write $x^T Bx = x^T B_s x$, where

$$B_s = \frac{B + B^T}{2}$$

is symmetric. Then

$$\begin{aligned} E(x^T B_s x + b^T x|y,\ z) &= \mu_2^T B_s \mu_2 + tr\left(B_s V_2\right) + b^T\mu_2 \\ &= \mu_2^T B\mu_2 + tr\left(BV_2\right) + b^T\mu_2 \end{aligned}$$

$$\begin{aligned} var\left(x^T B_s x + b^T x\middle|y,\ z\right) =& 2tr\left(B_s V_2 B_s V_2\right) + 4\mu_2^T B_s V_2 B_s \mu_2 + b^T V_2 b \\ &+ 2cov(x^T B_s x,\ b^T x|y,\ z) \end{aligned}$$

Note that when $w \sim N\left(0,\ I\right),\ cov\left(w^T B_s w,\ b^T w\right) = 0$.

Write $x = \mu_2 + Vw$, where $w|y,\ z \sim N\left(0,\ I\right),\ VV^T = V_2$,

$$cov\left(x^T B_s x,\ b^T x\middle|y,\ z\right) = 2\mu_2^T B_s VV^T b = 2\mu_2^T B_s V_2 b$$

Therefore,

$$\begin{aligned} var\left(x^T B_s x + b^T x\middle|y,\ z\right) =& 2tr\left(B_s V_2 B_s V_2\right) + 4\mu_2^T B_s V_2 B_s \mu_2 \\ &+ b^T V_2 b + 4\mu_2^T B_s V_2 b \end{aligned}$$

We then standardize the numerator of Wald statistic using above conditional mean and variance and use it as our corrected test statistic.

**Case with population structure and covariates**

Next, we consider a more generic scenario, where there are some covariates in the model. Suppose $y \sim N(A\alpha + \beta_1 x + \gamma z + \delta\left(x \circ z\right),\ \sigma_T^2\Sigma)$, where $A \in \mathbb{R}^{n\times k}$ is the covariate matrix, k<n. Here we assume $A$ contains the intercept term.

We want to first eliminate the covariate terms. Let $A' \in \mathbb{R}^{(n-k)\times n}$. Rows of $A\prime$ are linearly independent vectors in the orthogonal complement of the column space of A. Therefore, $A'A = 0_{(n-k)\times k}$.

In practice, we can get $A'$ by $svd: A = U\Lambda V,\ U \in \mathbb{R}^{n\times n},\ \Lambda \in \mathbb{R}^{n\times k},\ V \in \mathbb{R}^{k\times k}$. Then $U\left[,\ (k+1):n\right]^T$ can be our $A\prime$ because $U$ is an orthogonal matrix. After getting $A'$, we multiply $y$ by it:

$$A'y := y' \sim N(\beta_1 x' + \gamma z' + \delta(x \circ z)', \sigma_T^2 A'\Sigma A'^T)$$

Where $w' = A\prime w$ for any vector $w$.

Note that $A\prime$ has full row rank, so $A'\Sigma A'^T$ is positive definite and a valid variance matrix.

Now the model is very similar to the previous part with variables $y',\ x',\ z',\ (x\circ z)'$ and variance matrix being $A'\Sigma A'^T$, except that there is no more intercept term. Therefore, in the first step of forming $P_{\widetilde{u}}$, we do not need to regress out the 1 term and hence $\widetilde{H}=I$.

Then

$$S_{\widetilde{x'}\widetilde{y'}}=\left(\widetilde{H}\widetilde{x\prime}\right)^T\left(\widetilde{H}\widetilde{y\prime}\right)=\widetilde{x\prime}^T\widetilde{y}'=x'^TC^TCy'=x'^T\left(\widehat{A'\Sigma A}\right)^{-1}y'=x^TA'^T\left(\widehat{A'\Sigma A'^T}\right)^{-1}A'y:=x^T\hat{H}y$$

Where $\hat{H}=A'^T\left(\widehat{A'\Sigma A'^T}\right)^{-1}A'$.

After getting the new $\hat{H}$, the calculation of conditional mean and variance follows the same procedure.

#### Methods 1-3 when $x$ is Bernoulli

Here we design the methods that make use of the Bernoulli feature of $x$.

##### Non-GRM case

**Method 1: Estimate under the null & no heteroscedasticity correction.**

Estimated the conditional distribution $x|y,z$ by fitting a logistic model

$$x|y,z\sim Ber\left(p\right),\ p=logit(\mu+\alpha y+\beta z)$$

and use the fitted distribution to compute the conditional mean and variance of $x$ : $x|y,z\sim(\mu_2,\ V_2)$, where $\mu_2=\hat{p},\ V_2=diag(\hat{p}\left(1-\hat{p}\right))$

**Method 2: Estimate under the null & with heteroscedasticity correction**

Compute the conditional distribution of $x|y,z$ by Bayes rule:

$$p_{x|y,z}:=p\left(x=1|y,z\right)=\frac{p\left(y|x=1,\ z\right)p(x=1|z)}{p(y|z)} \quad (1)$$

$$=\frac{\left(2\pi\sigma_z^2\right)^{-\frac{1}{2}}e^{\frac{1}{2\sigma_z^2}\left(y-\alpha-\beta-\gamma z-\delta(1-m_x)(z-m_z)\right)^2}p(x=1|z)}{p\left(y|x=1,\ z\right)p\left(x=1|z\right)+p\left(y|x=0,\ z\right)p(x=0|z)} \quad (2)$$

For this method our estimations are all under the null, so $\delta=0$.

When $z$ is also Bernoulli, we test for heteroscedasticity using F-test:

Using the fact that $z$ only takes 2 values, assume

$$y|z=0\sim N\left(\mu,\ \omega_0^2\right):=N(\mu_0,\sigma_0^2)$$

$$y|z=1\sim N\left(\mu+\beta,\ \omega_0^2+\omega_1\right):=N(\mu_1,\sigma_1^2)$$

Testing for heteroscedasticity is equivalent to testing whether $\sigma_0^2=\sigma_1^2$. We can estimate them by

$$\hat{\mu}_0=\sum_{j:z_j=0}\frac{y_j}{n_0},\quad \hat{\sigma}_0^2=\sum_{j:z_j=0}\frac{\left(y_j-\hat{\mu}_0\right)^2}{n_0-1}$$

$$\hat{\mu}_1=\sum_{j:z_j=1}\frac{y_j}{n_1},\quad \hat{\sigma}_1^2=\sum_{j:z_j=1}\frac{\left(y_j-\hat{\mu}_1\right)^2}{n_1-1}$$

Where $n_k$ counts how many $z_j=k,\ k=0,\ 1$.

Under the null $H_0 : \sigma_0^2 = \sigma_1^2$, we have

$$\frac{\hat{\sigma}_0^2}{\hat{\sigma}_1^2} \sim \frac{\chi^2_{n_0-1}/(n_0-1)}{\chi^2_{n_1-1}/(n_1-1)} \frac{\sigma_0^2}{\sigma_1^2} = \frac{\chi^2_{n_0-1}/(n_0-1)}{\chi^2_{n_1-1}/(n_1-1)} \sim F_{n_0-1,\ n_1-1}$$

And we can do a two-sided F-test.

If we do not reject the null, we fit the ordinary linear model $y|z,\ x \sim N(\alpha + \beta x + \gamma z,\ \sigma^2)$ and use the estimated parameters to compute $\hat{p}_{x|y,z}$ using above Bayes formula.

If we reject the null, say, $var\,(y|z=0) = v_0^2,\ var\,(y|z=1) = v_1^2$, we hope to estimate the parameters in model $y|z,x \sim N\left(\alpha + \beta x + \gamma z,\ \sigma_0^2 \mathbb{I}_{z=0} + \sigma_1^2 \mathbb{I}_{z=1}\right)$. To do so, we first fit the weighted linear model $y|z,x \sim N\left(\alpha + \beta x + \gamma z,\ \sigma^2 W\right)$,$W$ is a diagonal matrix with i-th diagonal entry being $\hat{v}_0^2 \mathbb{I}_{z_i=0} + \hat{v}_1^2 \mathbb{I}_{z_i=1}$ and get estimated $\hat{\alpha}, \hat{\beta}, \hat{\gamma}, \hat{\sigma}^2$. Then we estimate $var(y|x,\ z)$ by $\hat{\sigma}_0^2 = \frac{1}{n_0-1.5}\Sigma_{i:z_i=0}\left(y_i - \hat{\alpha} - \hat{\beta}x_i\right)^2,\ \hat{\sigma}_1^2 = \frac{1}{n_1-1.5}\Sigma_{i:z_i=1}\left(y_i - \hat{\alpha} - \hat{\beta}x_i - \hat{\gamma}\right)^2$, where $n_k$ is the number of $z$ equaling $k$. Then we plug the estimated parameters in above Bayes formula to get $\hat{p}_{x|y,z}$. We estimate the conditional mean and variance of $x$ by $Ex|y,z \approx \hat{p}_{x|y,z},\ \ var\,(x|y,z) \approx \hat{p}_{x|y,z}(1 - \ \hat{p}_{x|y,z})$ and use this to compute $E(T|y,z)$ and $var(T|y,z)$.

The rest steps are the same as method 1.

When $z$ is not Bernoulli distributed, we check for heteroscedasticity in the same way as the normal approximation methods.

**Method 3: compute $var(T|y,z)$ under alternative model**

Also use Bayes rule, but compute var$(T|y,z)$ under alternative model: for $E(T|y,z)$, we use the conditional mean and variance of $x$ under the null; for $var(T|y,z)$, we use the conditional mean of $x$ under the null and conditional variance of $x$ under alternative.

The test for heteroscedasticity is the same as method 2.

For homoscedastic y:

For $E(T|y,z)$ :

$p(y_i|x_i,z_i)$ in $E(x|y,z)$ and $var(x|y,z)$ are given by fitting the null model

$$y|z,\ x \sim N(\alpha + \beta x + \gamma z,\ \sigma^2 I)$$

For $var(T|y,z)$:

$p(y_i|x_i,z_i)$ in $E(x|y,z)$ is still given by fitting the null model

$$y|x,z \sim N(\alpha + \beta x + \gamma z,\ \sigma^2 I)$$

$p(y_i|x_i,z_i)$ in $var(x|y,z)$ is given by fitting the interaction model

$$y|x,z \sim N(\alpha + \beta x + \gamma z + \delta(x \circ z), \sigma^2 I)$$

For heteroscedastic y,

For $E(T|y,z)$:

$p(y_i|x_i,z_i)$ in $E(x|y,z)$ and $var(x|y,z)$ are given by fitting the null heteroscedastic model

$$y|z,\ x \sim N(\alpha + \beta x + \gamma z,\ \sigma_0^2 \mathbb{I}_{z=0} + \sigma_1^2 \mathbb{I}_{z=1})$$

as in method 2.

For $var(T|y,z)$:

$p(y_i|x_i,z_i)$ in $E(x|y,z)$ is still given by fitting the model

$$y|x,z \sim N(\alpha + \beta x + \gamma z,\ \sigma_0^2 \mathbb{I}_{z=0} + \sigma_1^2 \mathbb{I}_{z=1})$$

$p(y_i|x_i, z_i)$ in $var(x|y,z)$ is given by fitting the interaction model

$$y|x,z \sim N(\alpha + \beta x + \gamma z + \delta(x \circ z), \sigma_0^2 \mathbb{I}_{z=0} + \sigma_1^2 \mathbb{I}_{z=1})$$

When $z$ is not Bernoulli, we estimate $var(y|x,z)$ by $\sigma^2 W$, where $W = v\hat{a}r(y|z)$.

The rest steps are the same as method 1 & 2.

**Extend the methods to GRM case**

**Method 1:**

Let $K$ be the GRM. Compute the conditional mean of $x: \ Ex_i|y,z = p(x_i = 1|y,z)$ by Bayes rule:

$$p\left(x_i = 1|y,z\right) \approx p\left(x_i = 1|y_i, z_i\right) = \frac{p\left(y_i|x_i = 1, \ z_i\right) p(x_i = 1|z_i)}{p(y_i|z_i)} \tag{3}$$

$$= \frac{p\left(y_i|x_i = 1, \ z_i\right) p(x_i = 1|z_i)}{p\left(y_i|x_i = 1, \ z_i\right) p\left(x_i = 1|z_i\right) + p\left(y_i|x_i = 0, \ z_i\right) p(x_i = 0|z_i)} \tag{4}$$

where

$$y|x,z \sim N(\alpha + \beta x + \gamma z, \ \sigma_g^2 K + \sigma_e^2 I)$$

$$y_i|x_i, \ z_i \sim N\left(\alpha + \beta x_i + \gamma z_i, \ \sigma_g^2 K + \sigma_e^2 I\right) \approx N(\hat{\alpha} + \hat{\beta} x_i + \hat{\gamma} z_i, \ \hat{\sigma}_g^2 K_{ii} + \hat{\sigma}_e^2)$$

Once get estimated $p\left(x_i = 1|y,z\right) := \hat{p}$, we can estimate $cov\left(x|y,z\right)$ by

$$\widetilde{K} = cov2cor\left(K + cI\right), \ c = 10^{-7}$$

$$cov\left(x|y,z\right)_{ij} \approx \widetilde{K}_{ij} \sqrt{\hat{p}_i\left(1 - \hat{p}_i\right) \hat{p}_j (1 - \hat{p}_j)}$$

The computation of approximated $T, \ E\left(T|y,z\right), \ var(T|y,z)$ is the same as non-GRM case except that $H = I - \frac{1}{n}\mathbf{1}\mathbf{1}^T, \ \hat{H} = \hat{\Sigma}^{-1} - \hat{\Sigma}^{-1} 1 \left(1^T \hat{\Sigma}^{-1} 1\right)^{-1} 1 \hat{\Sigma}^{-1}$, where $\hat{\Sigma} = \hat{\sigma}_g^2 K + \hat{\sigma}_e^2 I$ is the estimated covariance matrix in the LMM model

$$y \sim N(\alpha + \beta x + \gamma z + \delta(x \circ z), \ \sigma_g^2 K + \sigma_e^2 I)$$

**Method 2:**

Test against homoscedasticity:

Fit the null model

$$y|z \sim N(\alpha + \gamma z, \ \sigma_g^2 K + \sigma_e^2 I)$$

Fit models under alternative:

Let $I_0 = \{i : z_i = 0\}, \ I_1 = \{i : z_i = 1\}$

$$y_{I_0} \sim N(\alpha_0, \ \sigma_{g0}^2 K_{I_0, \ I_0} + \sigma_{e0}^2 I)$$

$$y_{I_1} \sim N(\alpha_1, \ \sigma_{g1}^2 K_{I_1, \ I_1} + \sigma_{e1}^2 I)$$

Conduct a chi-square test: if `pchisq(2*(llk1-llk0),2,lower.tail=F)< 0.15`, we reject the null.

If homoscedasticity is rejected, we fit the alternative model by `regress`

$$y|z \sim N(\alpha + \gamma z, \ \sigma_h^2 diag\left(z\right) + \sigma_g^2 K + \sigma_e^2 I)$$

Let $\hat{\Sigma} = \hat{\sigma}_h^2 diag\left(z\right) + \hat{\sigma}_g^2 K$, we estimate $y|x,z$ by

$$y|x,z \sim N(\alpha + \beta x + \gamma z, \ \sigma_b^2 \hat{\Sigma} + \sigma_e^2 I)$$

$$y_i|x_i, \ z_i \approx N(\hat{\alpha} + \hat{\beta} x_i + \hat{\gamma} z_i, \ \hat{\sigma}_b^2 \hat{\Sigma}_{ii} + \hat{\sigma}_e^2)$$

If homoscedasticity is not rejected, estimation of $y|x,z$ is the same as method 1.

Other parts are the same as method1.

**Method 3:**

For given $y$,$z$, test for heteroscedasticity as in method2.

If reject homoscedasticity:

Use `regress` to get the fitted variance component $\hat{\Sigma} = \hat{\sigma}_h^2 diag\,(z) + \hat{\sigma}_g^2 K$. For $E(T|y,z)$:

$p(y_i|x_i,z_i)$ in $E(x|y,z)$ and $var(x|y,z)$ are given by fitting the null model

$$y|x,z \sim N(\alpha + \beta x + \gamma z,\ \sigma_b^2 \hat{\Sigma} + \sigma_e^2 I)$$

For $var(T|y,z)$:

$p(y_i|x_i,z_i)$ in $E(x|y,z)$ is still given by fitting the null model

$$y|x,z \sim N(\alpha + \beta x + \gamma z,\ \sigma_b^2 \hat{\Sigma} + \sigma_e^2 I)$$

$p(y_i|x_i,z_i)$ in $var(x|y,z)$ is given by fitting the interaction model

$$y|x,z \sim N(\alpha + \beta x + \gamma z + \delta(x \circ z),\ \sigma_b^2 \hat{\Sigma} + \sigma_e^2 I)$$

If not reject homoscedasticity:

For $E(T|y,z)$:

$p(y_i|x_i,z_i)$ in $E(x|y,z)$ and $var(x|y,z)$ are given by fitting the null model

$$y|x,z \sim N(\alpha + \beta x + \gamma z,\ \sigma_b^2 K + \sigma_e^2 I)$$

For $var(T|y,z)$:

$p(y_i|x_i,z_i)$ in $E(x|y,z)$ is still given by fitting the null model

$$y|x,z \sim N(\alpha + \beta x + \gamma z,\ \sigma_b^2 K + \sigma_e^2 I)$$

$p(y_i|x_i,z_i)$ in $var(x|y,z)$ is given by fitting the interaction model

$$y|x,z \sim N(\alpha + \beta x + \gamma z + \delta(x \circ z),\ \sigma_b^2 K + \sigma_e^2 I)$$

Other parts are the same as method1 & 2.

#### Methods 1-3 using Normal approximation

Another, more general, approach to approximating the first and second conditional moments of $G_j$ is to assume a multivariate normal model for $(G_j, Y)|Z$. More specifically, we assume:

$$Y = \alpha + \gamma Z + \beta G_j + \delta(G_j - m_j)(Z - m_z) + \epsilon$$

and

$$\begin{pmatrix} G_j \\ \epsilon \end{pmatrix} |Z \sim N_2\left(\begin{pmatrix} \mu_{j|z} \\ 0 \end{pmatrix}, \begin{pmatrix} \sigma^2_{j|z} & 0 \\ 0 & \sigma^2_{\epsilon|z} \end{pmatrix}\right) \tag{5}$$

where $\mu_{j|z} = 1_n m_j + (Z - m_z)\eta_{j|z}$. This results in a model of the form

$$\begin{pmatrix} G_j \\ Y \end{pmatrix} |Z \sim N_2\left(\begin{pmatrix} \mu_{j|z} \\ \mu_{y|z} \end{pmatrix}, \begin{pmatrix} \sigma^2_{j|z} & (\beta + \delta(Z - m_z))\sigma^2_{j|z} \\ (\beta + \delta(Z - m_z))\sigma^2_{j|z} & v_{y|z} \end{pmatrix}\right). \tag{6}$$

Then we can compute the conditional distribution:

$$G_j|\,(Y,\ Z) \sim\ N(\mu_{j|z} + \frac{(\beta + \delta(Z - m_z))\,\sigma^2_{j|z}}{v_{y|z}}(y - \mu_{y|z}),\ \sigma^2_{j|z} - \frac{(\beta + \delta(Z - m_z))^2\,\sigma^4_{j|z}}{v_{y|z}})$$

**Method 1** We estimate $\mu_{y|z},\ v_{y|z}$ by fitting the ordinary linear regression

$$y|z \sim N(\alpha + \gamma z, \sigma^2 I)$$

Set $\delta = 0$ and estimate $\beta$ the parameters by

$$y|z, x \sim N(\alpha + \gamma z + \beta x, \sigma^2 I)$$

**Method 2** We check for heteroscedasticity of $y|z$ by fitting a linear mixed model

$$y|z \sim N(\alpha + \gamma z, \sigma_2^2 diag(z^2) + \sigma_1^2 diag(z) + \sigma_0^2 I)$$

and do a log-likelihood test against the null model in Method 1. If the null hypothesis is rejected, we estimate $\mu_{y|z},\ v_{y|z}$ by the fitted LMM, otherwise, we use the null model.

We then estimate $\beta$ by

$$y|z, x \sim N(\alpha + \gamma z + \beta x, \sigma^2 W)$$

where $W = I$ for the homoscedasticity case;
$W = diag(w_1, ..., w_n),\ w_i = \hat{\sigma}_2^2 z_i^2 + \hat{\sigma}_1^2 z_i + \hat{\sigma}_0^2$ for the heteroscedasticity case.

**Method 3** Similar to the Bernoulli methods, for $var(T|y, z)$, parameters in $var(x|y, z)$ are estimated by the alternative model

$$y|z, x \sim N(\alpha + \gamma z + \beta x + \delta(x \circ z), \sigma^2 W)$$

The GRM version works in a similar way, except an additional variance component $K$ for $y$.

#### Fast approximate Wald test

##### Fast approximate t-test

The basic idea is to regress out everything else except $y$ and the interaction term. Then the t-statistics for $m$ $x$'s can be computed at once via matrix multiplication.

Suppose we have phenotype $y$, SNPs $z,\ X = (x_1, x_2, ..., x_m)$ and covariates $u$. Let $z_c,\ X_c$ be the centered genotypes. Let $W = X_c \circ z_c = (x_{1c} \circ z_c, ..., x_{mc} \circ z_c)$ be the matrix of interactions. We want the t-statistics for each of the epistasis by fitting $m$ linear models

$$y \sim N(u\alpha_i + \gamma_i z_c + \beta_i x_{ic} + \delta_i (x_{ic} \circ z_c), \sigma_i^2 I)$$

**Step 1: regress $u,\ z_c$ out of $y,\ X_c, Xc \circ z_c$** Let $P$ be the matrix that projects to the subspace spanned by $(u, z_c)$. We let

$$y_r = y - Py$$

$$X_r = X_c - PXc$$

$$W_r = X_c \circ z_c - P(X_c \circ z_c)$$

**Step 2: regress each column of $X_r$ out of $y_r$ and each column of $W_r$** Since when the subspace only has 1 dimension, the projection matrix can be directly written as $\frac{1}{\|x_r\|^2} x_r x_r^T$ and the resulting variables can be computed by matrix operations in R. Let the results be $y_{rx}$ and $W_{rx}$

**Step 3: get the interaction t-statistics by regress each column of $W_{rx}$ on corresponding column of $y_{rx}$** Again, since each projection subspace has dimension 1, the result can be got by matrix operations in R

This fast method gets all $m$ t-statistics without running a loop of $m$ iterations.

**Fast approximate Wald test**

When fitting a linear mixed model, we can modify above method to get approximated test statistics. Suppose we want to fit the model

$$y \sim N(\alpha_i + \gamma_i z + \beta_i x_i + \delta_i (x_i \circ z), \Omega)$$

where $\Omega = \sigma_g^2 K + \sigma_e^2 I$. We could approximate it by a linear model by pre-multiply everything by $\Omega^{-1/2}$:

$$\Omega^{-1/2} y \sim N(\alpha_i (\Omega^{-1/2} \mathbf{1}) + \gamma_i (\Omega^{-1/2} z) + \beta_i (\Omega^{-1/2} x_i) + \delta_i \Omega^{-1/2} (x_i \circ z), I)$$

We could let $u = \Omega^{-1/2} \mathbf{1}$ be the new covariate and apply the fast t-statistics method to the new variables.

#### GCIF of more cases

##### $z$, $x$ normal

Figure S2 are histograms GCIF got by running simulation setting 6 500 times

**Fig S2.** $z$, $x$ **both normal**

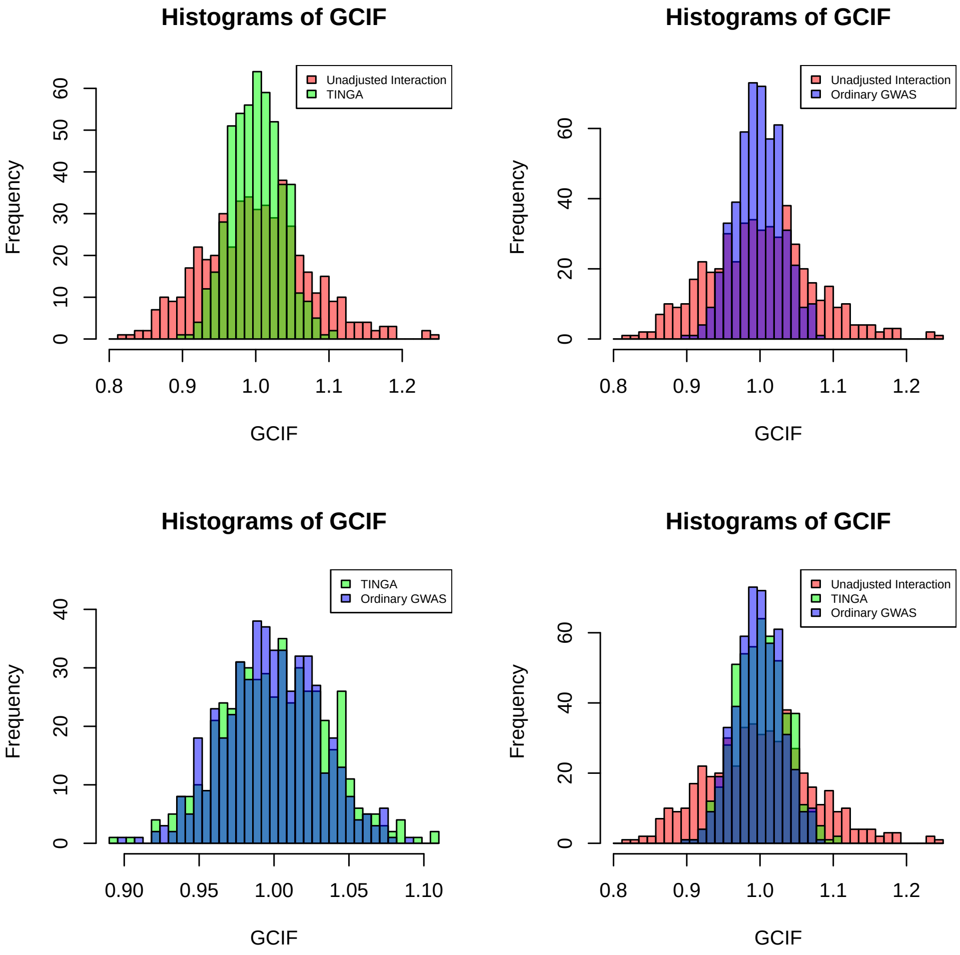

$z,\ x$ **binomial**

Figure S3 are histograms GCIF got by running simulation setting 7 500 times

**Fig S3.** $z,\ x$ **both binomial**

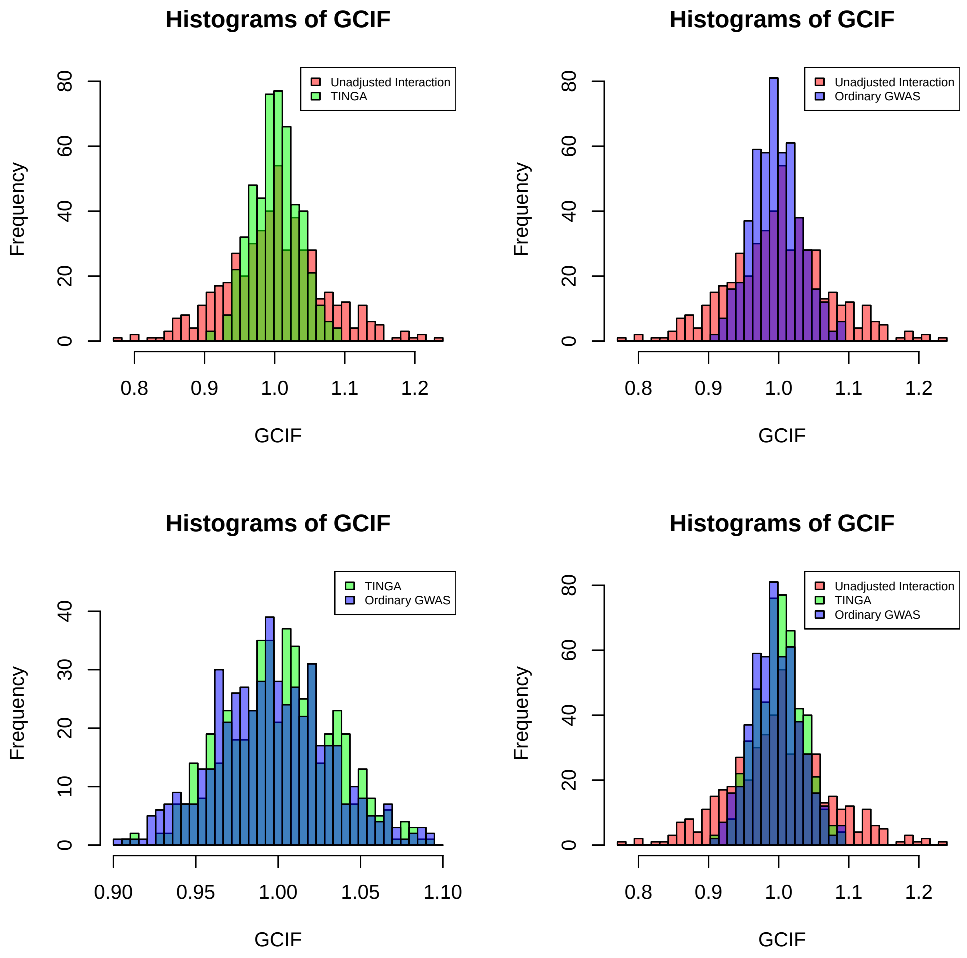

#### Type I error rates and power for more cases

**$z$ normal, $x$ Binomial** We run simulation setting 8 5000 times. Table S1 shows Type I error across 5000 replicates. Table S2 shows power.

**Table S1. Type I error at level 0.05** non-GRM, Normal approximation, $z$ normal, $x$ Binomial

| $x$ | Uncorrected | Method 1 | Method 2 | Method 3 |
|---|---|---|---|---|
| $x_1$ | 0.0534 | 0.0486 | 0.0494 | 0.0508 |
| $x_2$ | 0.0602 | 0.0564 | 0.0576 | 0.0586 |
| $x_3$ | 0.0576 | 0.0504 | 0.0508 | 0.0518 |

**Table S2. Power** non-GRM, Normal approximation, $z$ normal, $x$ Binomial

| p-value cutoff | Uncorrected | Method1 | Method2 | Method3 |
|---|---|---|---|---|
| $10^{-5}$ | 0.6818 | 0.6152 | 0.6382 | 0.6918 |
| $10^{-6}$ | 0.5036 | 0.4222 | 0.4454 | 0.5232 |

**$z$ normal, $x$ Bernoulli** We run simulation setting 9 5000 times. Table S3 shows Type I error across 5000 replicates. Table S4 shows power.

**Table S3. Type I error at level 0.05** non-GRM, Normal approximation, $z$ normal, $x$ Bernoulli

| $x$ | Uncorrected | Method1 | Method2 | Method3 |
|---|---|---|---|---|
| $x_1$ | 0.0544 | 0.0480 | 0.0482 | 0.0504 |
| $x_2$ | 0.0572 | 0.0532 | 0.0534 | 0.0546 |
| $x_3$ | 0.0536 | 0.0482 | 0.0498 | 0.0514 |

**Table S4. Power** non-GRM, Normal approximation, $z$ normal, $x$ Bernoulli

| p-value cutoff | Uncorrected | Method1 | Method2 | Method3 |
|---|---|---|---|---|
| $10^{-5}$ | 0.6896 | 0.6224 | 0.6436 | 0.7004 |
| $10^{-6}$ | 0.5204 | 0.4388 | 0.4678 | 0.5344 |

Since $x$'s are Bernoulli distributed, we could also use the Bernoulli version methods. Table S5 and S6 are the results (from the same simulated data as the normal approximation methods).

**Table S5. Type I error at level 0.05** non-GRM, Bernoulli methods, $z$ normal, $x$ Bernoulli

| $x$ | Method1 | Method2 | Method3 |
|---|---|---|---|
| $x_1$ | 0.0480 | 0.0480 | 0.0500 |
| $x_2$ | 0.0532 | 0.0532 | 0.0572 |
| $x_3$ | 0.0484 | 0.0504 | 0.0522 |

**$z$ Binomial, $x$ Binomial** We run simulation setting 10 5000 times. Table S7 shows Type I error across 5000 replicates. Table S8 shows power.

**Table S6. Power** non-GRM, Bernoulli methods, $z$ normal, $x$ Bernoulli

| p-value cutoff | Method1 | Method2 | Method3 |
|---|---|---|---|
| $10^{-5}$ | 0.6294 | 0.6528 | 0.7224 |
| $10^{-6}$ | 0.4476 | 0.4738 | 0.5646 |

**Table S7. Type I error at level 0.05** non-GRM, Normal approximation, $z$ Binomial, $x$ Binomial

| $x$ | Uncorrected | Method1 | Method2 | Method3 |
|---|---|---|---|---|
| $x_1$ | 0.0508 | 0.0474 | 0.0484 | 0.0500 |
| $x_2$ | 0.0574 | 0.0532 | 0.0552 | 0.0574 |
| $x_3$ | 0.0496 | 0.0468 | 0.0472 | 0.0488 |

**Table S8. Power** non-GRM, Normal approximation, $z$ Binomial, $x$ Binomial

| p-value cutoff | Uncorrected | Method1 | Method2 | Method3 |
|---|---|---|---|---|
| $10^{-5}$ | 0.6878 | 0.6324 | 0.6526 | 0.6974 |
| $10^{-6}$ | 0.5034 | 0.4376 | 0.4626 | 0.5314 |

#### Minimum cell count

For a pair of binary SNPs $(z, x)$, we can build a 2x2 table (Table S9) where each cell is the number of individuals with one possible combination:

**Table S9. Cell counts**

| | $z = 0$ | $z = 1$ |
|---|---|---|
| $x = 0$ | $\#(z, x) = (0, 0)$ | $\#(z, x) = (1, 0)$ |
| $x = 1$ | $\#(z, x) = (0, 1)$ | $\#(z, x) = (1, 1)$ |

Meaning of cell counts for a pair of Binary SNPs

Minimum cell count (MCC) is the smallest count in above table. If it is too small, it means there are very few samples with a certain combination of $z$ and $x$ value. This may make the estimation inaccurate.

#### Feast or Famine Effect in a Variety of Methods

We show that the feast or famine effect occurs across a wide range of GxE analysis methods, including but not limited to (1) testing interaction in a linear or linear mixed model (LMM) using standard approaches such as t-tests/Wald tests, likelihood ratio tests, or score tests; (2) doing a combined interaction-association test in a linear model or LMM using standard approaches such as F-tests or likelihood ratio tests; (3) testing interaction with multiple environments or multiple SNPs, where these are modeled as random effects in a LMM using standard approaches; (4) performing tests of interaction in a GWIS where significance is assessed using permutation of the trait residuals.

#### Supplemental Fig. 4

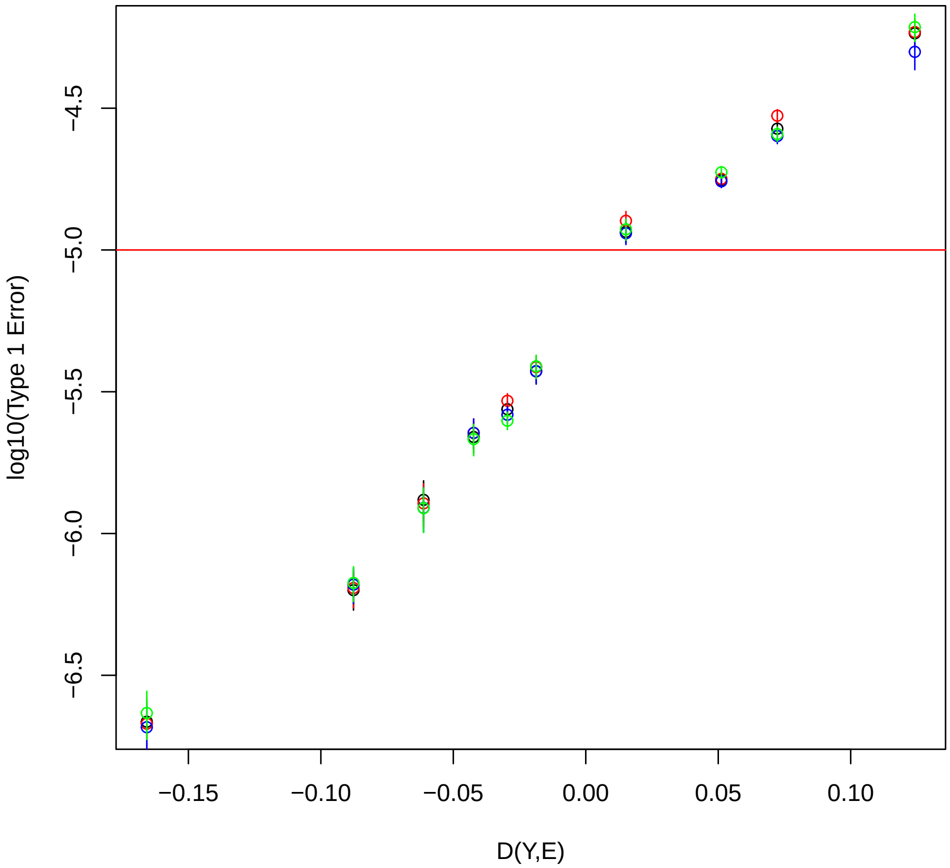


**Fig S4. In a GWIS, Type 1 error of t-test, score test, likelihood ratio test and permutation test in a linear model is well-predicted by a function of only the trait $Y$ and environment $E$** In each bin, type 1 error at level 1e-5 is based on 5 million replicates in each of 72 GWASs, for a total of 3.6e-8 replicates. 720 GWIS's were simulated under the null hypothesis assuming a linear model, and these GWISs were grouped into 10 bins based on their value of the predictor $D(Y, E)$, which is a function of only the observed values of the trait and environment variables. Type 1 error was combined across the GWIS's in each bin. Black points represent the t-test, blue points the score test, red points the likelihood ratio test, and green points the permutation test based on permuting the residuals of $Y$. Vertical segments represent standard errors on the type 1 error.

#### Supplemental Fig. 5

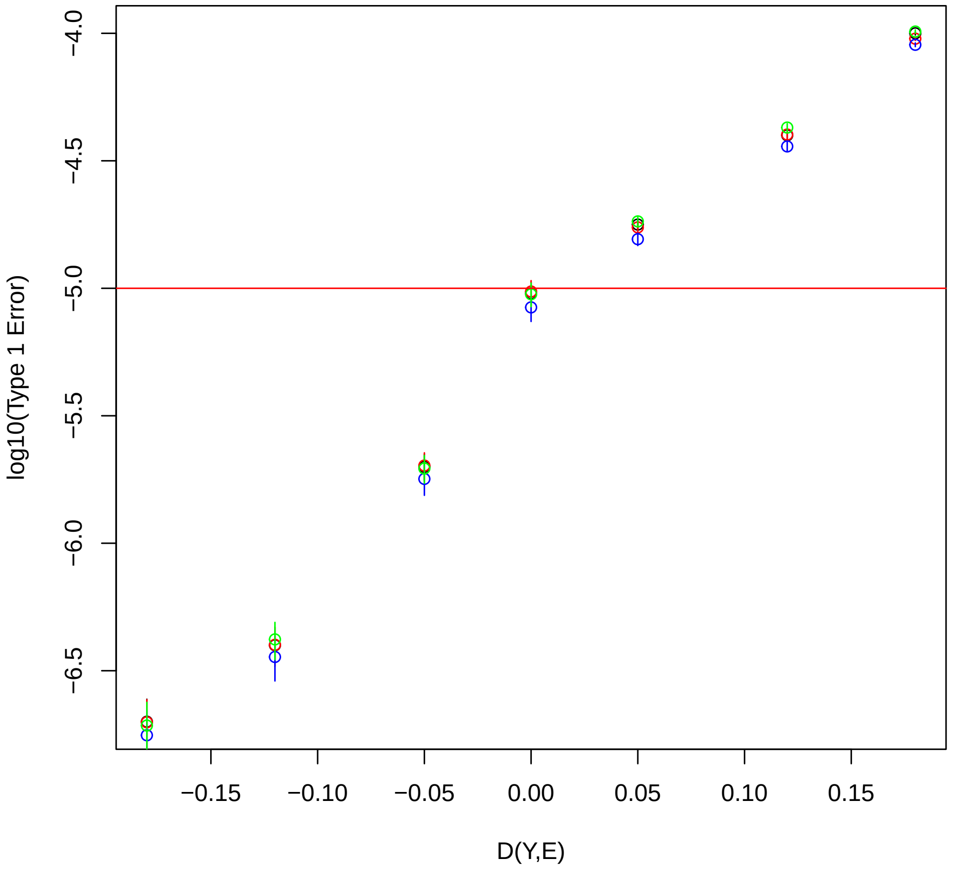


**Fig S5. In a GWIS, Type 1 error of Wald, score test, likelihood ratio test and permutation test in a linear mixed model is well-predicted by a function of only the trait $Y$ and environment $E$** In each bin, type 1 error at level 1e-5 is based on 5 million replicates in each of 50 GWASs, for a total of 2.5e-8 replicates. 490 GWIS's were simulated under the null hypothesis assuming a linear mixed model, and these GWISs were grouped into 7 bins based on their value of the predictor $D(Y, E)$, which is a function of only the observed values of the trait and environment variables. Type 1 error was combined across the GWIS's in each bin. Black points represent the Wald test, blue points the score test, red points the likelihood ratio test, and green points the permutation test based on permuting the decorrelated residuals of $Y$. Vertical segments represent standard errors on the type 1 error.

#### Supplemental Fig. 6

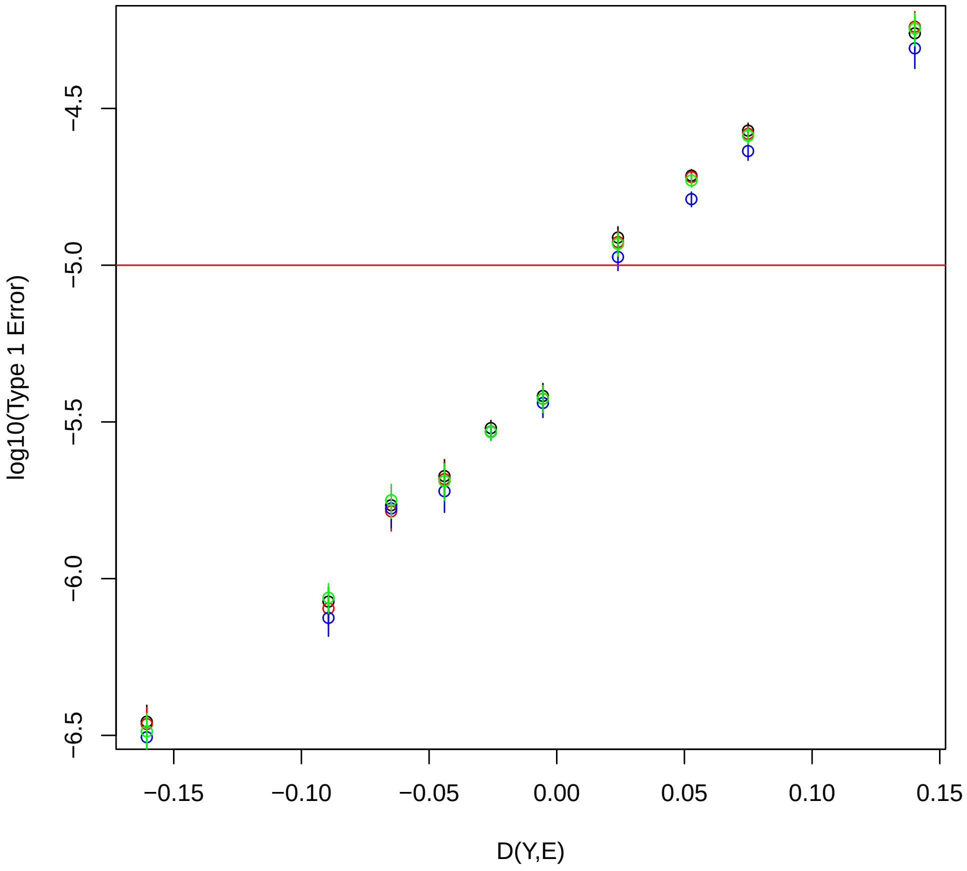


**Fig S6. In a GWIS, Type 1 error of joint association-interaction using F-test, score test, likelihood ratio test and permutation test in a linear model is well-predicted by a function of only the trait $Y$ and environment $E$** In each bin, type 1 error at level 1e-5 is based on 5 million replicates in each of 72 GWASs, for a total of 3.6e-8 replicates. 720 GWIS's were simulated under the null hypothesis assuming a mixed model, and these GWISs were grouped into 10 bins based on their value of the predictor $D(Y, E)$, which is a function of only the observed values of the trait and environment variables. Type 1 error was combined across the GWIS's in each bin. Black points represent the F-test, blue points the score test, red points the likelihood ratio test, and green points the permutation test based on permuting the residuals of $Y$. Vertical segments represent standard errors on the type 1 error. The same set of simulations were used as in Fig. S4, but different tests were performed.

#### Supplemental Fig. 7

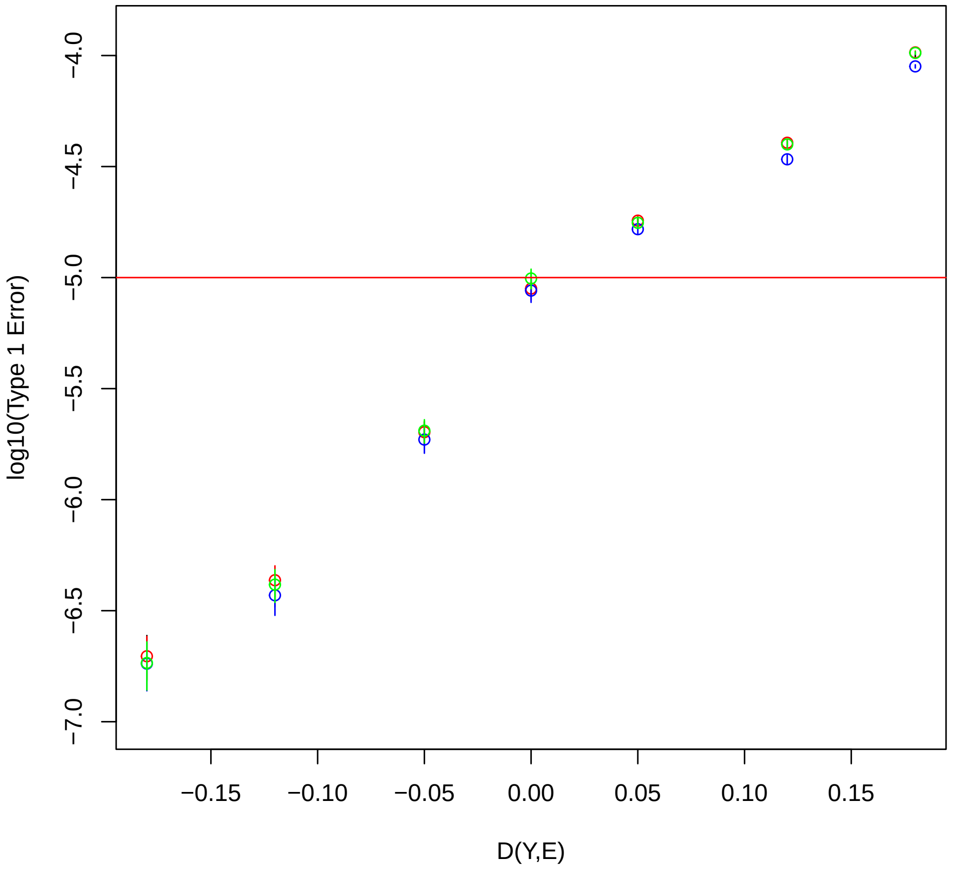


**Fig S7. In a GWIS, Type 1 error of joint association-interaction using score test, likelihood ratio test and permutation test in a linear mixed model is well-predicted by a function of only the trait $Y$ and environment $E$** In each bin, type 1 error at level 1e-5 is based on 5 million replicates in each of 70 GWASs, for a total of 2.5e-8 replicates. 490 GWIS's were simulated under the null hypothesis assuming a mixed model, and these GWISs were grouped into 7 bins based on their value of the predictor $D(Y, E)$, which is a function of only the observed values of the trait and environment variables. Type 1 error was combined across the GWIS's in each bin. Blue points the score test, red points the likelihood ratio test, and green points the permutation test based on permuting the decorrelated residuals of $Y$. Vertical segments represent standard errors on the type 1 error. The same set of simulations were used as in Fig. S5, but different tests were performed.

#### Supplemental Fig. 8

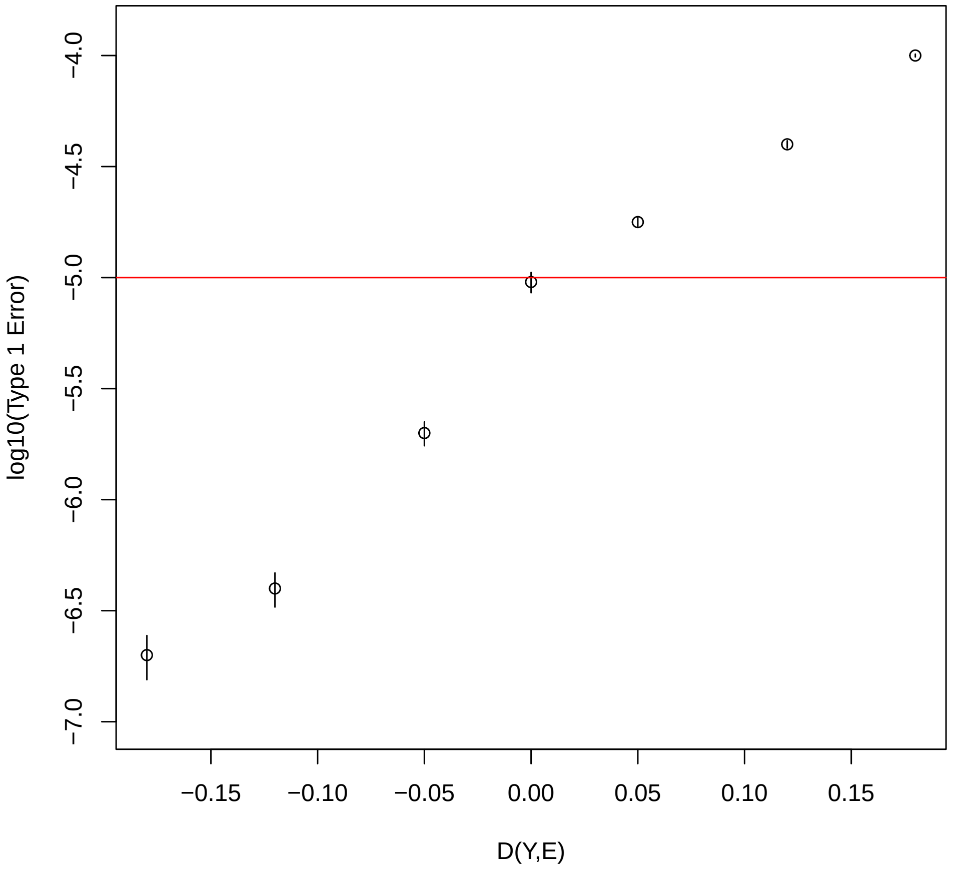


**Fig S8. In a GWIS, Type 1 error of multi-environment GxE is well-predicted by a function of only the trait $Y$ and environment $E$** The multi-environment GxE was tested as a random effect in a linear mixed model using the method of Moore et al. (2019)[2]. 10 environments were used, 5 of which were standard normal and 5 of which were Bernoulli(.2). In each simulation, the diagnostic ratio $D(Y,E)$ was calculated as the average of the co-kurtosis between $Y$ and each $E_j$, $j = 1, \ldots, 10$. In each bin, type 1 error at level 1e-5 is based on 5 million replicates in each of 70 GWASs, for a total of 2.5e-8 replicates. 490 GWIS's were simulated under the null hypothesis assuming a linear model, and these GWISs were grouped into 7 bins based on their value of the predictor $D(Y,E)$, which is a function of only the observed values of the trait and environment variables. Type 1 error was combined across the GWIS's in each bin. Vertical segments represent standard errors on the type 1 error.

#### Additional type 1 error problems that arise when testing interaction with multiple environments or multiple SNPs, where these are modeled as random effects in a LMM using standard approaches

In addition to the "feast or famine effect" shown in Fig. S8, there is another source of potential type 1 error problems that arises specifically when testing interaction with multiple environments or multiple SNPs, where these are modeled as random effects in a LMM using standard approaches[2, 3]. For example, in the multi-environment GxE test of Moore et al.[2], the LMM in which interaction is tested includes a marginal fixed effect for G, but the marginal effect of E is included only as a random effect, not a fixed effect, in the LMM. In this setting, as we show below, the type 1 error for interaction can be inflated when one or more of the environments are discrete and are associated with both $G$ and $Y$. (Technically, the problem arises when both (i) $\mathrm{Var}(E_i|G)$ depends on $G$ and (ii) $E_i$ has an effect on $Y$ beyond that explained by $G$. When $G$ is a genotype and $E_i$ is discrete, then association between $G$ and $E_i$ would commonly result in (i), so that is why it particularly tends to be a problem with discrete $E_i$.) As an example, we consider the following simulation setting. Suppose $G = (G_1, \ldots, G_n)^T$, is the centered and standardized genotype vector for a SNP with observed allele frequency 0.8. Suppose that we have $p = 3010$ environmental variables, $E_1, \ldots, E_{3010}$ where each $E_i$ is a vector of length $n$ whose elements were originally i.i.d. binomial(2,.8), where each $E_i$ is then centered and standardized. Suppose $E_1, \ldots, E_{10}$ each have correlation .25 with $G$, while $E_{11}, \ldots, E_{3010}$ are i.i.d. and independent of $(G, E_1, \ldots, E_{10})$. Suppose $Y$ is simulated as $Y = \sum_{i=1}^{10} E_i + \epsilon$, where $\epsilon \sim N_n(0, \sigma^2 I_n)$, with $\sigma^2 = 5$. Suppose we apply structLMM[2] to test for multi-environment interaction between $G$ and the multiple environments $E_1, \ldots, E_{3010}$. We take $n = 2000$ individuals in the sample, and repeat this experiment 100 times. We find that in 37% of the trials under the null hypothesis, the type 1 error was $< 0.05$, which is significantly different from the nominal level (p-value $= 1 \times 10^{-22}$ using exact binomial method), with 17% of the trails having type 1 error $< 0.01$ (p-value $3 \times 10^{-16}$ for difference from the nominal level), and 3% of the trials having type 1 error $< 0.001$ (p-value $1.5 \times 10^{-4}$ for difference from the nominal level). This example shows the potential for the variance component interaction test to have significantly inflated type 1 error in some settings.
